# Dual-wield NTPases: a novel protein family mined from AlphaFold DB

**DOI:** 10.1101/2023.02.19.529160

**Authors:** Koya Sakuma, Ryotaro Koike, Motonori Ota

**Author notes:** Email: K.S., M.O.

## Abstract

AlphaFold protein structure database (AlphaFold DB) archives a vast number of predicted models. We conducted systematic data mining against AlphaFold DB and discovered an uncharacterized P-loop NTPase family. The structure of the protein family was surprisingly novel, showing an atypical topology for P-loop NTPases, noticeable two-fold symmetry and two pairs of independent putative active sites. Our findings show that structural data mining is a powerful approach to identifying undiscovered protein families.

## Introduction

Characterizing protein structures is essential for understanding their function, and structures are typically solved by experimental approaches and deposited in the Protein Data Bank (PDB) (Burley et al. 2022). When the solved protein adopts a novel structure that appeared at the first time, the finding is usually reported by the researchers who determined it. However, more recently, public databases produced by state-of-the-art structure prediction, such as the AlphaFold protein structure database (AlphaFold DB) and ESM metagenomic Atlas (ESM Atlas), are changing this situation (Varadi et al. 2022; Lin et al. 2023). These databases are approximately three orders of magnitude larger than the PDB and contain numerous experimentally unsolved protein structures. Structural models never seen by human-being must be hiddenly archived there since the models were generated automatically by artificial intelligences and deposited without any human curations, providing opportunities for finding novel proteins based only on the structural information *in-silico*.

Dedicated data mining demands a clearly stated working hypothesis. While several groups have pursued intensive model classifications against AlphaFold DB (Bordin et al. 2023; Inigo Barrio Hernandez et al. 2023; Durairaj et al. 2023), this bird’s-eye approach could miss unique and intriguing proteins. To find these hidden gems, we defined a very specific database-search question: are there monomeric proteins that contain multiple phosphate-binding loops (P-loops) on a single continuous β-sheet? The P-loop or Walker-A motif is a local functional motif that recognizes phosphate groups and shared among P-loop NTPases, such as ATPases, GTPases and nucleotide kinases (NKs) (Walker et al. 1982; Saraste, Sibbald, and Wittinghofer 1990; Leipe, Koonin, and Aravind 2003; Leipe et al. 2002; Leipe, Koonin, and Aravind 2003). In general, one P-loop resides on a single continuous β-sheet of a three-layered α/β/α sandwich architecture. Our preliminary search against the PDB supported this observation because no structure has multiple P-loops in a β-sheet. However, the possibility that a single β-sheet possesses multiple P-loops should not be excluded. We hypothesized that such experimentally unobserved multiple-P-loop structures exist in AlphaFold DB and can be discovered via systematic data mining.

## Results

By computationally scanning more than 0.2 billion entries in AlphaFold DB version 4 (Varadi et al. 2022; Kim, Mirdita, and Steinegger 2023), we extracted 15,977 single-chained structures possessing multiple P-loops. We then analyzed the hydrogen-bond network and extracted 839 structures with multiple P-loops on a single continuous β-sheet (Frishman and Argos 1995). The structures were grouped into 11 clusters based on structural similarity (Van Kempen et al. 2023). As a result, we found an uncharacterized family of P-loop proteins, dual-wield P-loop NTPase (dwNTPase), as the largest cluster with 711 members. All structural models in this cluster were predicted with high confidence scores, i.e., the average predicted Local Distance Difference Test (pLDDT) was 94.27, indicating that the predictions were reliable (Figure supplement 1) (Jumper et al. 2021).

The overall architecture of dwNTPases was highly novel and showed noticeable two-fold symmetry. Figure 1a shows the structure of a representative dwNTPase from *Bacillus thuringiensis* (Bt. UniProt accession No.: A0A1Y0TWD8). Two P-loop domains are tightly packed and surrounded by two long bridging α-helices and two framing α-helices. Each of the domains comprises six β-strands, and twelve strands form a continuous single β-sheet, which reveals a previously unobserved single-sheeted dual-P-loop architecture (Figure 1b). Two β-hairpins from each domain form a pier-like four-stranded β-sheet to connect two domains. Two canonical P-loops independently form two putative ligand binding sites that penetrate through the molecule and resemble tunnels rather than pockets (Figure 1c). A search against the PDB clarified that no similar structures have been reported (Van Kempen et al. 2023; Minami et al. 2018). Similarly, a SwissProt subset of AlphaFold DB contained no similar structures (Van Kempen et al. 2023; Minami et al. 2018), indicating that the dwNTPase family has no reliable annotations manually verified by UniProt curators.

**Figure 1.**
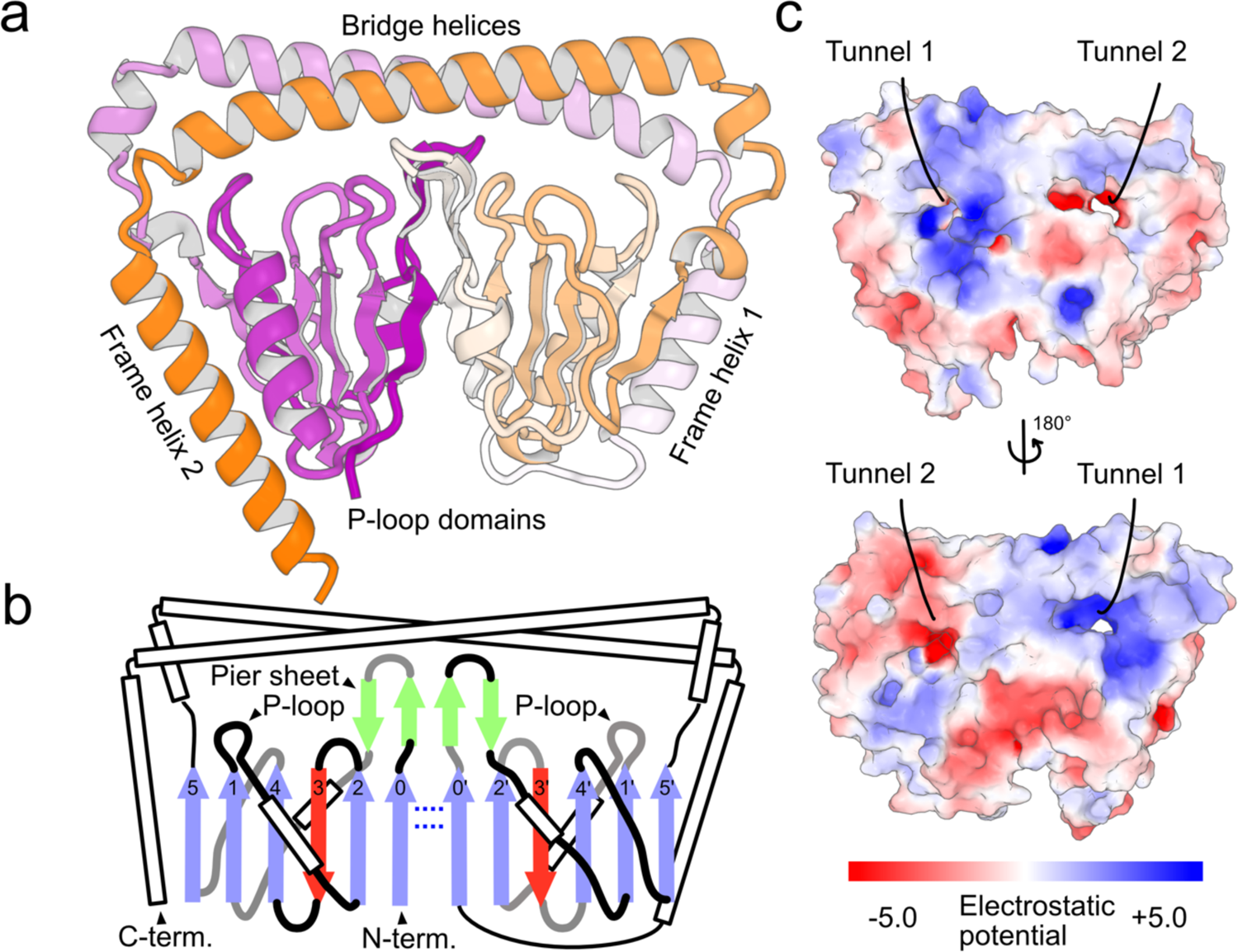
AlphaFold2 predicted model of a dual-wield NTPase structure (AF-A0A1Y0TWD8-F1-model_v4). **a**, Overall dwNTPase structure colored according to a purple-white-orange gradient from the N- to C-terminus. **b**, Topology diagram of dwNTPase. Blue and red arrows represent β-strands pointing up and down that form the large β-sheets in the P-loop domains. Green arrows represent the pier sheet. White rectangles are α-helices. Grey and black lines indicate junctions projecting behind and out of the β-sheets, respectively. Blue dotted lines represent hydrogen bonds connecting the two halves of the large β-sheet. **c**, The location and shape of the ligand binding tunnels. Color bar is at the bottom.

We found that the P-loop domain of dwNTPases was structurally atypical for a P-loop NTPase by searching against the PDB (Figure 2a) (Van Kempen et al. 2023; Minami et al. 2018). A crystal structure of mutual gliding-motility protein MglAa from *Myxococcus xanthus* (PDB ID: 6h35), a bacterial small and monomeric GTPase, was the only known P-loop NTPase that showed relevant structural similarity to the dwNTPase P-loop domain (Galicia et al. 2019). The P-loop domain of dwNTPase has an additional β-strand at the N-terminus (strand 0) compared to the MglAa structure (Figure 2b). Two strands constituting the pier sheet and a successive α-helix are also appended. In contrast, the domain lacks two C-terminal β-strands (strands 6 and 7) and some other surrounding secondary structural elements (SSEs). These unique arrangements of SSEs give rise to the atypical topology that does not resemble other P-loop NTPases (Figure supplement 2 and 3)(Minami et al. 2018; Chandonia et al. 2022). Furthermore, the P-loop domain has a long loop rather than a helix conserved in other P-loop NTPases (Figure supplement 4), which we named the switch loop (Figure 2a). These atypical features of the P-loop domain make it difficult to assign dwNTPase to known classes of P-loop NTPases.

**Figure 2.**
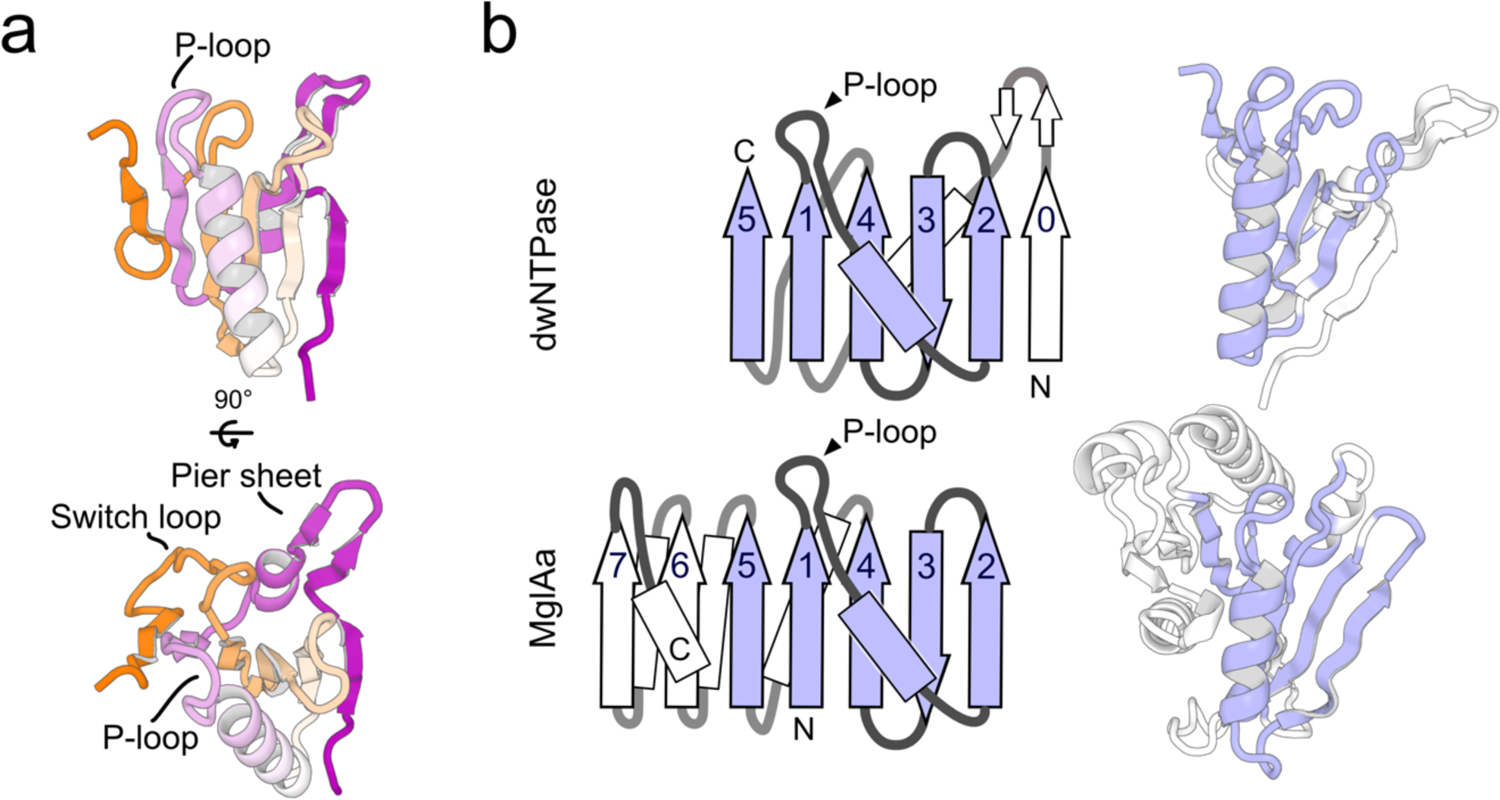
P-loop domain. **a**, Front and top views of the dwNTPase P-loop domain colored according to a purple-white-orange gradient from the N- to C-terminus. P-loop, switch loop and pier-sheet are indicated by labels. **b**, Topology diagrams and cartoon representations of dwNTPase P-loop domains and MglAa structure. Arrows and rectangles represent β-strands and α-helices. Secondary structures that align between two structures are colored blue.

Despite these novel features of dwNTPase, an iterative structure search by Foldseek against the entire AlphaFold DB revealed that 2,219 similar structures were deposited, most of which originated from bacteria in various Firmicutes (Table 1 and Supplement Data Table 1) (Varadi et al. 2022; Van Kempen et al. 2023). Similar searches against ESM Atlas culled by 30% sequence identity found 748 similar structures (Supplement Data Table 2) (Lin et al. 2023). We classified dwNTPase structures into six sub-classes based on the conservation of motifs and domains (Figure supplement 5). The bona fide dwNTPase structure with two P-loops intact (class 1) was the most abundant, suggesting functional constraints exist to conserve the two active P-loops. A BLAST search against the non-redundant database revealed that dwNTPase had been classified as the PRK06851 family protein in the NCBI conserved domain database (McGinnis and Madden 2004; J. Wang et al. 2023). Thus, we concluded that dwNTPases constitute a conserved protein family among bacteria.

**Table 1.**
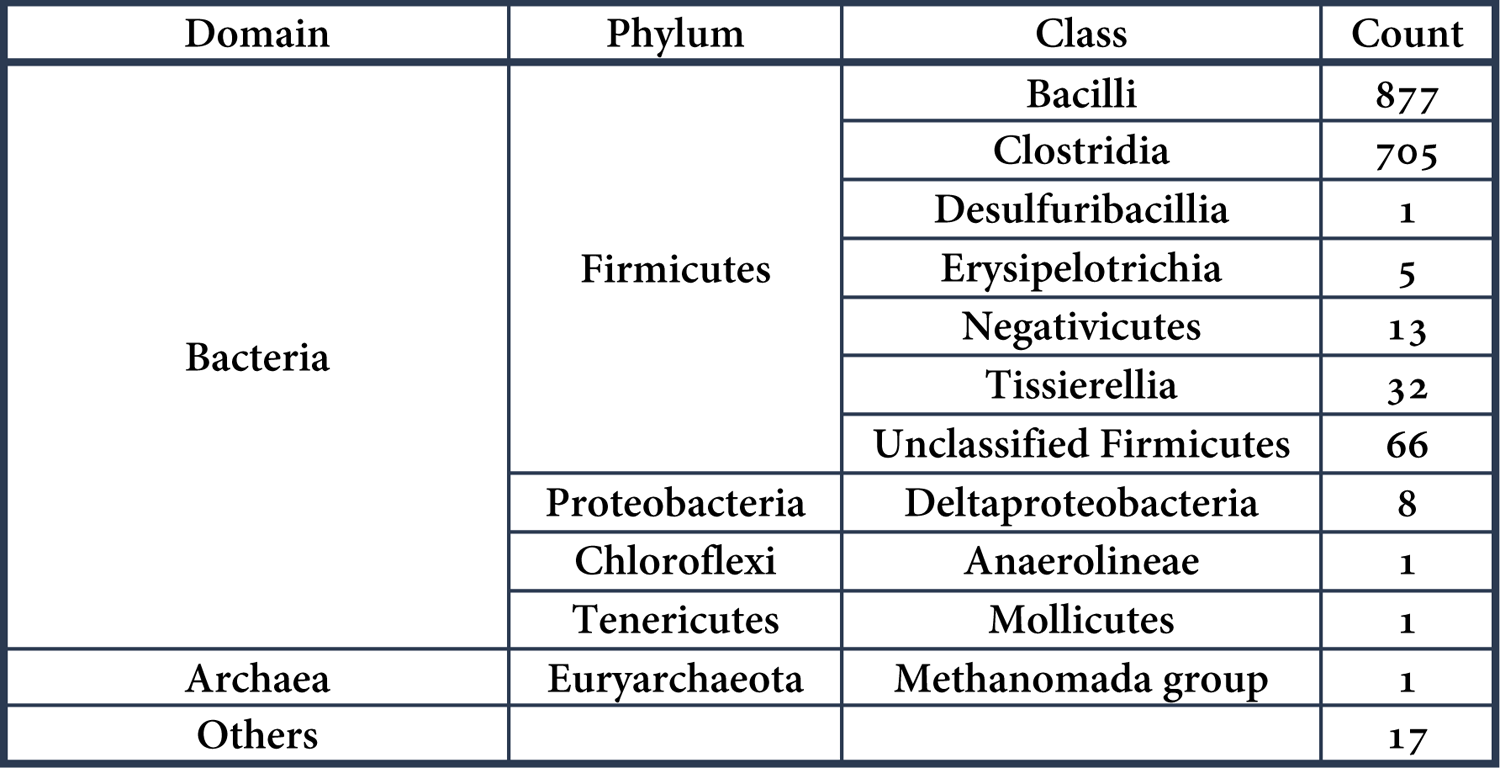
Phylogenetic classification of dwNTPases

| Domain | Phylum | Class | Count |
| --- | --- | --- | --- |
| Bacteria | Firmicutes | Bacilli | 877 |
|  |  | Clostridia | 705 |
|  |  | Desulfuribacillia | 1 |
|  |  | Erysipelotrichia | 5 |
|  |  | Negativicutes | 13 |
|  |  | Tissierellia | 32 |
|  |  | Unclassified Firmicutes | 66 |
|  | Proteobacteria | Deltaproteobacteria | 8 |
|  | Chloroflexi | Anaerolineae | 1 |
|  | Tenericutes | Mollicutes | 1 |
| Archaea | Euryarchaeota | Methanomada group | 1 |
| Others |  |  | 17 |

## Discussion

The molecular functions of dwNTPases were investigated by analyzing conserved residues. The most symmetric class of dwNTPases (class 1) possesses two clusters of conserved residues shared between both halves (Figure 3a and Figure supplement 6). We found Cys66/Cys248 (residue numbers follow Bt. dwNTPase), Asp74/Asp256, Asp87/Asp269 and His92/His274 formed putative metal binding sites. Molecular dynamics simulations of the Bt. dwNTPase structure complexed with two ATPs, two Mg^2+^ ions and two Zn^2+^ ions showed that the Zn^2+^ ions were stably coordinated by two aspartates and the γ-phosphate group of ATPs (Figure 3b) (Abraham et al. 2015; Huang et al. 2017), which resembles the active site structure of metal-dependent nucleotidyl-transfer enzymes (Figure 3c) (Yang 2008). The side chains of Cys66/Cys248 and His92/His274 remained unoccupied (Figure 3d), suggesting that they may have roles other than metal-binding. As the pair of cysteine and histidine residues are reminiscent of the catalytic triad/dyad in cysteine proteases (Figure 3e), we hypothesize that dwNTPases have additional hydrolase/ligase activity (Dodson and Wlodawer 1998).

**Figure 3.**
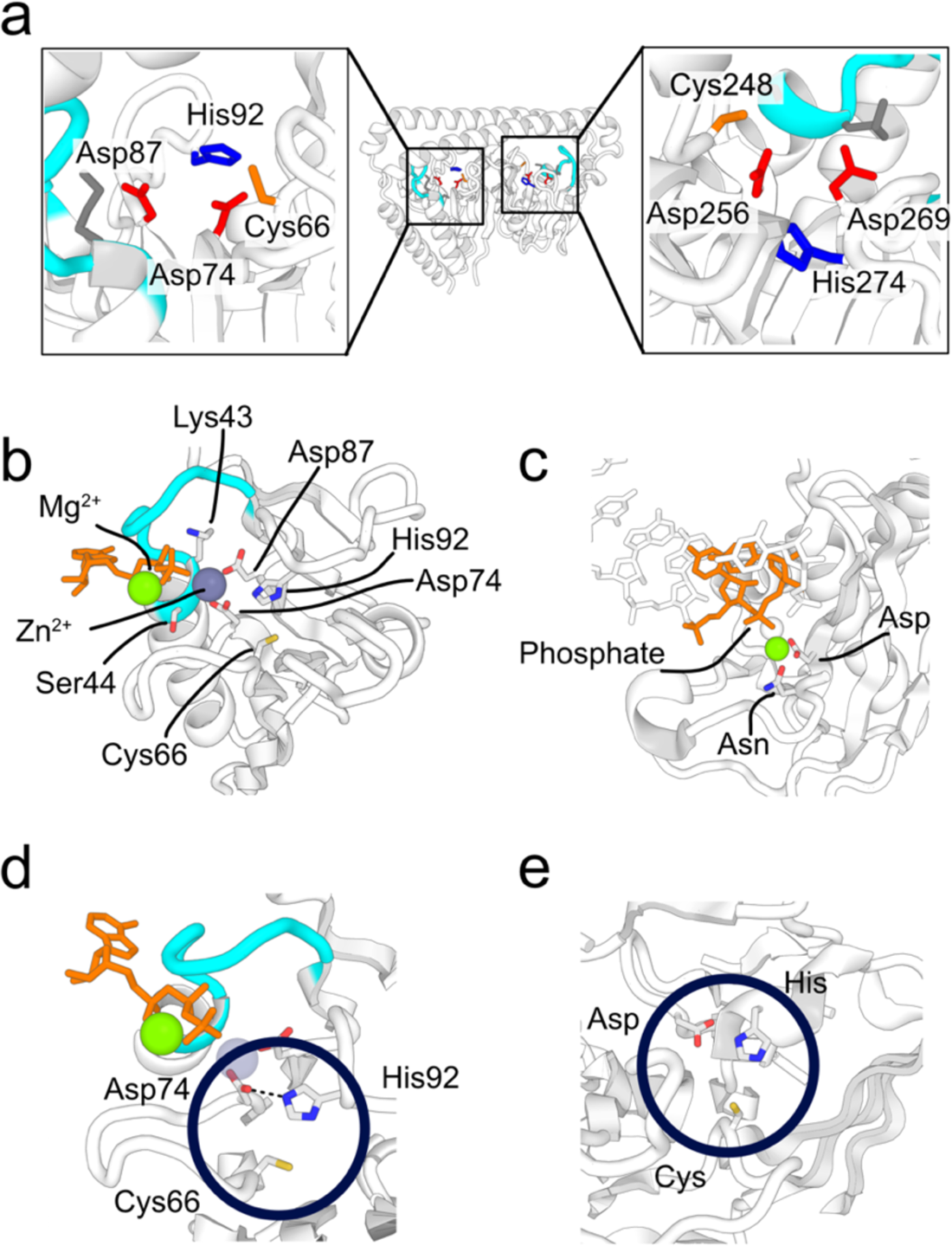
Putative functionally relevant residues. **a**, Conserved residues in the putative ligand binding tunnels. His, Cys and Asp are colored blue, orange and red, respectively. P-loops and their conserved residues are colored cyan and grey. **b**, Coordination of metal ions by two aspartate side chains observed in MD simulations. ATP is shown in orange stick representation. Side chains of relevant residues are shown as sticks and CPK coloring. Green and grey spheres represent Mg^2+^ and Zn^2+^ ions, respectively. The P-loop is colored cyan. **c**, The active site structure of RNase H (PDB ID: 1zbl). The side chain of metal coordinating amino acid residues Asp and Asn are shown as sticks and CPK coloring, where Asn is a mutation from Asp. Mg^2+^ ions are shown as spheres. The Mg^2+^ ion coordinating with the side chain of Asp and Asn is colored green. Nucleic acid residues that contact the Mg^2+^ ion are shown in orange. **d**, The catalytic triad-like side chain configuration observed during the MD simulations. The triad-like side chain cluster is circled. The black dotted line indicates the hydrogen bond between the side chains of His92 and Asp74, which HBPLUS detected. **e**, The active site structure of TEV protease (PDB ID: 1lvm). Side chains of the Cys-His-Asp catalytic triad are shown as sticks, CPK coloring and circled.

In addition to these conserved residues, we identified other regions characteristic of dwNTPases. First, each P-loop domain has conserved lysine residues (Lys36/Lys218) that precede the P-loops and interact with two switch loops. Because the switch loop partially conceals the ligand binding tunnels (Figure supplement 7a) and is highly flexible in MD simulations (Figure supplement 7b), the conserved lysine residues may play sensor-like roles to trigger NTPase activity, depending on the binding of other ligands to the tunnels. Additionally, the P-loops are surrounded by several charged or polar residues that support the recognition of NTPs and Mg^2+^ ions (Figure supplement 7c) and are not conserved in known P-loop NTPases (Leipe et al. 2002; Leipe, Koonin, and Aravind 2003).

The biological functions of dwNTPases remain elusive because their structures show limited homology to NTPases with known functions, indicating that dwNTPases are responsible for unique biological functions. The two-fold symmetry implies that the substrates of dwNTPases also possess two-fold symmetry, such as double-stranded DNA, or that the cleft between two P-loop domains recognizes ligand molecules in a similar manner to periplasmic heme-binding proteins (Figure supplement 8) (Mattle et al. 2010). The left half (residues 1–139 and 321–369) of the structure in Figure 1a is more positively charged than the right half (140–320), indicating that each half plays different functional roles (Figure 1c and Figure supplement 9). Because dwNTPases are distributed among various Firmicutes, especially among Bacillus (Table 1), their biological functions may be related to spore-formation, which is characteristic of the species. The evolutionary origin of dwNTPases is unknown. Although it is plausible that dwNTPases gained two-fold symmetry via gene duplication, the origin of the P-loop domains remains unclear. Detailed phylogenetic analysis may explain the evolution of P-loop NTPases, including dwNTPases (Leipe et al. 2002; Leipe, Koonin, and Aravind 2003). Structural and biochemical studies are required and should provide greater insight into the biological significance of the dwNTPase family.

In summary, we demonstrated that structural data mining based on specific working hypothesis can discover uncharacterized protein families and is a powerful approach to exploring dark proteomes (Taylor et al. 2009; Perdigão et al. 2015), the unwatched region of the protein universe, which will help and encourage the design of experimental studies.

We performed structural alignment of all 2219 structures against the representative dwNTPase structure. To ensure fragmented structures were excluded, 1843 structures showing TM-scores > 0.85 were selected. Entries with no phylogenetic information available in UniProt were ignored. The structures (1727 in total) were classified by their species. Others include environmental samples, metagenomes, unclassified bacteria and Firmicutes from environmental samples.

## Materials and methods

### Identification of structures containing P-loop-like fragments

AlphaFold DB (v4 UniProt) was downloaded from the Foldcomp database (Varadi et al. 2022; Kim, Mirdita, and Steinegger 2023). P-loop NTPase protein structures were extracted by converting the models into the sequences of ABEGO, where A, B, E and G, respectively, denote backbone dihedral angles (phi, psi) for α, β, left-handed β and left-handed α on the Ramachandran plot (Wintjens, Rooman, and Wodak, n.d.). O denotes other conformations unassignable on the Ramachandran plot, typically a cis-peptide conformation. Typical P-loop (Walker-A) motifs have conformations represented by EBBGAG or BBBGAG, both of which can be seen in the crystal structure of α and β subunits of bovine mitochondrial F1-ATPase (chain A and chain D of PDB ID: 1BMF) (Abrahams et al. 1994). Because the P-loop is a junction between a β-strand and an α-helix, we extended the ABEGO motifs to “BBBEBBGAGAAAAA” or “BBBBBBGAGAAAAA” and extracted all the structures containing any of them by sequence pattern matching. We then calculated the Cα root-mean-square deviations (RMSDs) of the matched substructures against the reference P-loop fragment (residues 166–179 of 1BMF, chain A) and filtered out substructures with RMSDs larger than 2.0 Å. We obtained 15,977 proteins containing multiple P-loop-like fragments and built a custom Foldcomp database for subsequent procedures (Kim, Mirdita, and Steinegger 2023; Steinegger and Söding 2017).

### Identification of dual-wield NTPases

Visual inspection revealed that most structures with multiple P-loop-like fragments within a single chain were tandem repeats of known P-loop NTPase domains connected by flexible linkers. Such proteins were excluded by analyzing structures using an in-house program that detects the β-sheet hydrogen-bond network and extracting structures possessing two P-loop-like fragments on a single continuous β-sheet (Frishman and Argos 1995). We obtained 839 structures possessing two P-loop-like fragments on a single β-sheet. These structures were clustered by TM-score calculations with Foldseek into 11 clusters (Van Kempen et al. 2023). The largest cluster contained 711 members, which corresponded to dual-wield NTPases. For these structures, we performed all-against-all structure alignment using MICAN and defined the structure with the largest average TM-score as the representative (AF-A0A1Y0TWD8-F1-model_v4) (Minami et al. 2018).

### Extraction of structures similar to dwNTPase from AlphaFold DB and ESM Atlas

We performed iterative structure searches using FoldSeek to enumerate as many structures that resemble dwNTPase as possible (Van Kempen et al. 2023). In the first stage, we performed a structure search against AlphaFold DB using all 711 structures initially mined from AlphaFold DB as queries. After removing overlapping structures, we obtained 1377 structures. Using these structures as seeds, we again performed a Foldseek search and obtained 135 new non-overlapping structures. The third iteration of Foldseek search yielded some non-specific hits. Therefore, we stopped this iteration, manually selected similar structures and discarded the remaining structures. Consequently, we obtained 2219 dwNTPase structures from AlphaFold DB. Similarly, we performed structural searches against the highquality_clust30 subset of ESM-atlas using 711 structures found in AlphaFold DB as queries and obtained 748 structures with a TM-score larger than 0.5 (Lin et al. 2023; Van Kempen et al. 2023; Xu and Zhang 2010).

### Whole structure search against the PDB and Swiss-Prot subset of AlphaFold DB

To assess the novelty of the dwNTPase structure and gain insights into the function, we performed structural searches against PDB100 and the Swiss-Prot subset of AlphaFold DB (version 4) using the Foldseek server in the TM-align mode and the representative structure as the query (Burley et al. 2022; Varadi et al. 2022; Van Kempen et al. 2023). No relevant (TM-score ≧ 0.5) hit was found among these databases. We used MICAN to perform rigorous one-against-all searches without pre-filtering; however, no similar (TM-score ≧ 0.5) structures were found among the PDB (2023-09-Jan) and the Swiss-Prot subset of AlphaFold DB (version 2) (Minami et al. 2018).

### Domain structure search against the PDB, Swiss-Prot subset of AlphaFold DB and SCOPe

We searched structures similar to the P-loop domain of the representative structure (residues 1–110) against the PDB100 and the Swiss-Prot subset of AlphaFold DB by using the Foldseek server (Van Kempen et al. 2023). No relevant hit was found. We used MICAN to perform a rigorous structure search without pre-filtering against the PDB (2023-09-Jan) and the Swiss-Prot subset of AlphaFold DB (version 2) (Minami et al. 2018). We obtained 358 and 2931 relevant hits (TM-score ≧ 0.5) from the PDB and Swiss-Prot. We performed clustering by mmseq2 with sequence identity set at 35% and obtained 15 and 137 clusters (Steinegger and Söding 2017). The alignments were checked by visual inspection of all cluster representatives. We found that some structures showed similar topology to the P-loop domain of dwNTPase: 6h35, Q1DB04 and Q9UBK7 from the PDB and Swiss-Prot, which are annotated as GTPase or GTP-binding proteins (Galicia et al. 2019). The remaining hits showed RecA-like topology and were not topologically identical to dwNTPase because the RecA-like topology has an all-parallel β-sheet, whereas dwNTPases have anti-parallel-containing β-sheets. Similarly, we performed structural comparisons against domain structures classified as G-proteins (SCOP concise classification string (SCCS): c.37.8), NKs (c.37.1) and RecA-like proteins (c.37.11) in the SCOPe version 2.08 using MICAN (Minami et al. 2018; Chandonia et al. 2022). The groups of G-proteins, NKs and RecA-like proteins contained 255, 212 and 118 parsed domain structures, respectively, and we selected the structures that showed the highest TM-score in the group for visualization (Figure supplement 2).

### Calculation of sequence identities between two halves of dwNTPase structures

We selected 1,903 structures with more than 340 residues from the set of dwNTPases extracted from AlphaFold DB. A structure was self-aligned by MICAN in the rewiring mode, which ignores the sequential order of SSEs (Minami et al. 2018). The sequence identity was calculated based on the second-best alignment by MICAN.

### Identification of putative catalytic residues and a side chain pattern search against the PDB

The potential function of dwNTPases was examined by performing a sequence search and alignment to identify conserved residues by HHblits against UniRef30_2022_02 (Remmert et al. 2012; Suzek et al. 2015). After three iterations, 2687 sequences were extracted from the database. To exclude fragmented sequences most likely originating from partial matches to the P-loop consensus motif, we removed aligned sequences with more than ten gaps against the representative sequence and obtained a Multiple Sequence Alignment (MSA) with 138 sequences. From this MSA, the site-wise entropy of the alignments was calculated to identify conserved residues, and the top 10 residues around the two tunnels were listed. We defined tunnel 1 as residues 61–100 and tunnel 2 as residues 243–282. From tunnel 1, residues 62, 66, 67, 69, 73, 74, 75, 87, 88 and 100 were identified. From tunnel 2, residues 244, 246, 247, 248, 252, 255, 256, 261, 263 and 274 were identified. According to the orientation of side chains toward the tunnel, we selected Cys66, Ser67, Asp74 and Asp87 as candidates for probable functional residues in tunnel 1. Similarly, Cys248, Asp256 and His274 were selected for tunnel 2. Considering the symmetry of the dwNTPase structure, Cys66/Cys248, Asp74/Asp256, Asp87/Asp269 and His92/His274 were considered clusters of functional residues in tunnels 1 and 2. We performed a side chain pattern search against the PDB using the strucmotif-search program to determine whether protein structures possessed similar side chain configurations (Bittrich, Burley, and Rose 2020). The set of residues Cys66, Asp74, Asp87 and His92 in the representative structure were selected as queries, and a search was performed against all structures in the PDB (2022-28-12), with the threshold for the structure similarity set to 1.0 Å. The side chain pattern search gave no hits and indicated that the putative catalytic residues have a novel configuration of conserved residues.

### Docking of ATP, Mg and Zn

We transplanted ligand structures from existing PDB structures to model the complex structures. The P-loop region of an ATPase crystal structure (PDB ID: 6j18) was superposed to the P-loop of the representative structure by MICAN in PyMOL, and the ATP and Mg^2+^ models were extracted (Minami et al. 2018; S. Wang et al. 2020; Schrodinger 2015). Similarly, His125 from a zinc finger motif (PDB ID: 2hgh) was superposed to His92 and His274, and the coordinating Zn^2+^ ions were extracted (Lee et al. 2006). The extracted ligand molecules were merged with the representative structure.

### MD simulations

MD simulations were performed by Gromacs version 2022.04 with the charmm36 force field (Abraham et al. 2015; Huang et al. 2017). The size of simulation boxes was determined by the molecule size with margins of 13 Å. After *in-vacuo* energy minimization to remove steric clashes, the protein-ligand complex was solvated by the TIP3P water model with 0.1 M NaCl, and the system was neutralized by adding additional Na^+^ or Cl^-^ ions, depending on the total charge of the protein and ligands. The energy was minimized by the steepest descent and equilibrated by 100 ps NVT and NPT simulations with harmonic restraints on the non-hydrogen atoms. The temperature and pressure of the system were controlled to 300 K and 1 bar by the V-rescale thermostat and Parrinello-Rahman barostat. Electrostatic interactions were computed by the particle mesh Ewald method, and bonds involving hydrogen atoms were constrained by the LINCS algorithm. For each docked model, we performed 20 trajectories of 100 ns simulations with a 2 fs time step.

### Figure preparation

The images of molecular structures were created by PyMOL and Mol* viewer (Schrodinger 2015; Sehnal et al. 2021). The surface electrostatic potential was calculated by the PyMOL APBS plugin (Jurrus et al. 2018). Hydrogen bonds were detected by HBPLUS and visualized by PyMOL (McDonald and Thornton 1994). Secondary structure elements were assigned by DSSP and illustrated by ESPript (Kabsch and Sander 1983; Robert and Gouet 2014).

## Author contributions

K.S. designed and initiated the study, performed structure mining, structure searches, structure comparisons, complex modeling, MD simulations, interpreted data and prepared and wrote the manuscript. R.K. and M.O. interpreted data and wrote the manuscript.

## Acknowledgements

K.S. would like to thank George Chikenji for providing the program to analyze hydrogen-bonding patterns in β-sheets, Milot Mirdita for providing instructions on building the custom Foldcomp databases, Naoya Kobayashi and Shintaro Minami for their suggestions on extracting functionally relevant residues to infer functions from structures, Nobu C. Shirai for his constructive comments on the manuscript, and Shigeo S. Sugano and Ryo Ozuka for discussing the feasibility and significance of structure-based protein mining, which inspired K.S. to design this study. Structure searches by Foldseek and MD simulations were carried out on the supercomputer “Flow” at the Information Technology Center, Nagoya University. This study was supported by KAKENHI grant numbers JP21H00394 to R.K. and 20H05932 to M.O. We thank Edanz (https://jp.edanz.com/ac) for editing a draft of this manuscript.

---

**Figure supplement 1.**
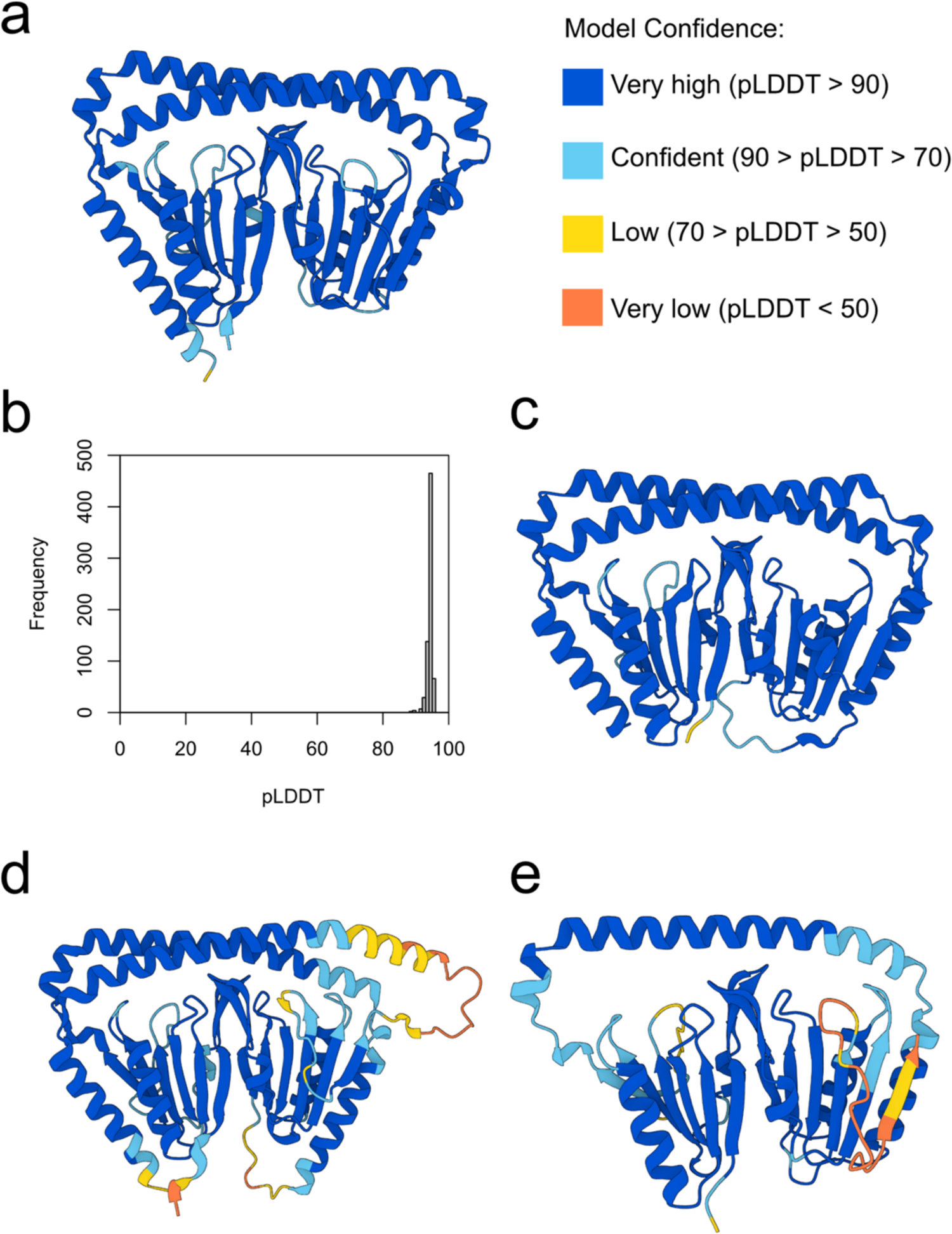
Model confidence of initially mined dwNTPase structures. **a**, The cartoon shows the representative dwNTPase structure (UniProt accession No.: A0A1Y0TWD8). The cartoon is colored according to the residue-wise values of pLDDT. The coloring scheme is shown on the right of the structure, which is the same as displayed in AlphaFold DB. **b**, Distribution of total pLDDT values among the 711 dwNTPase structures initially mined from AlphaFold DB. **c–e**, The most confident, second worst and worst predictions among the 711 dwNTPase structures (A0A7V6TLZ4, A0A7C6J2V5 and W4RL53) gave average pLDDTs of 95.67, 88.43 and 88.01, respectively. Structures were created by Mol* viewer.

**Figure supplement 2.**
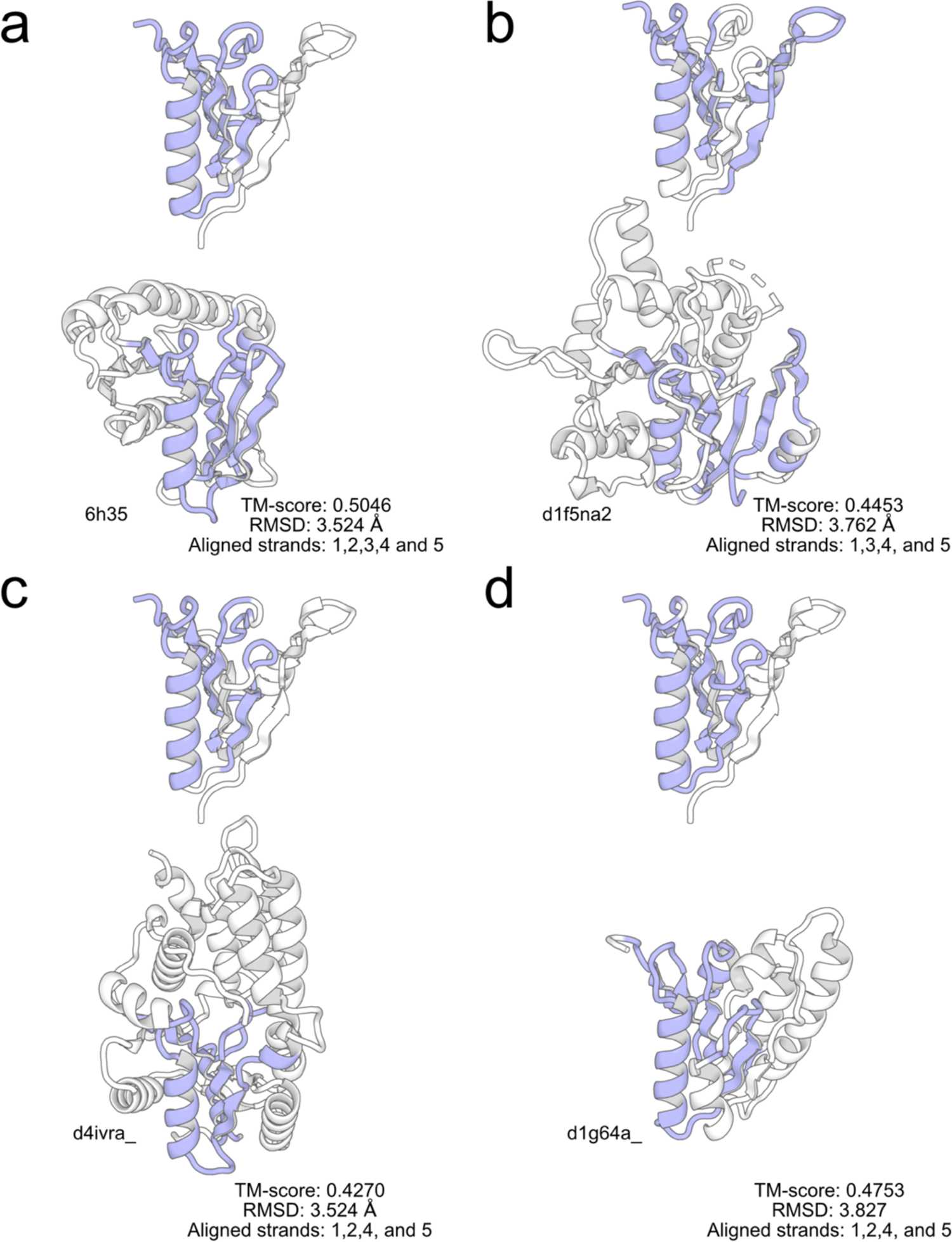
Structure of the dwNTPase P-loop domain compared with other representative P-loop NTPase protein structures. Aligned structures of the dwNTPase P-loop domain and known NTPases. **a**, MglAa, (**b**) G-proteins, (**c**) NKs and (**d**) Rec-A like. The P-loop domain of dwNTPase and known NTPase are shown on the top and bottom of the panels, respectively. The latter three structures were selected from c.37.8, c.37.1 and c.37.11 SCCSs in SCOPe that showed the highest similarity to the dwNTPase P-loop domain. The aligned regions are colored blue. The PDB or SCOPe IDs are shown below the structures. The RMSD, TM-score and list of aligned strands in the dwNTPase P-loop domain are summarized at the bottom of the panels.

**Figure supplement 3.**
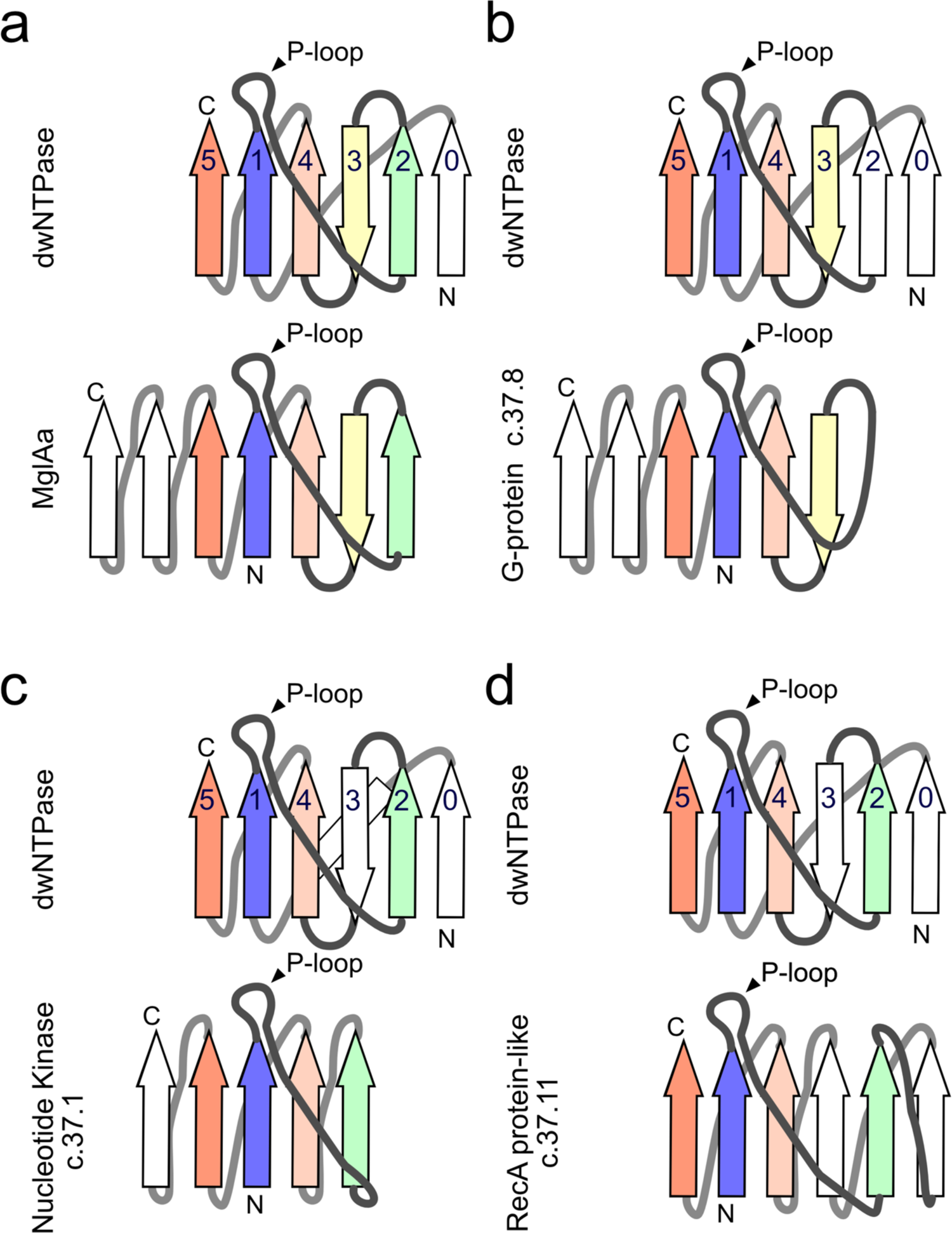
Topology diagram of the dwNTPase P-loop domain compared with other representative P-loop NTPases. **a**, Comparisons with MglAa, (**b**) G-proteins, (**c**) NKs and (**d**) RecA-like proteins. Arrows represent strands. Grey and black lines indicate junctions projecting behind and out of the β-sheet. The pair of aligned strands are colored in the same colors. Unaligned strands are colored white. Helices are omitted for clarity.

**Figure supplement 4.**
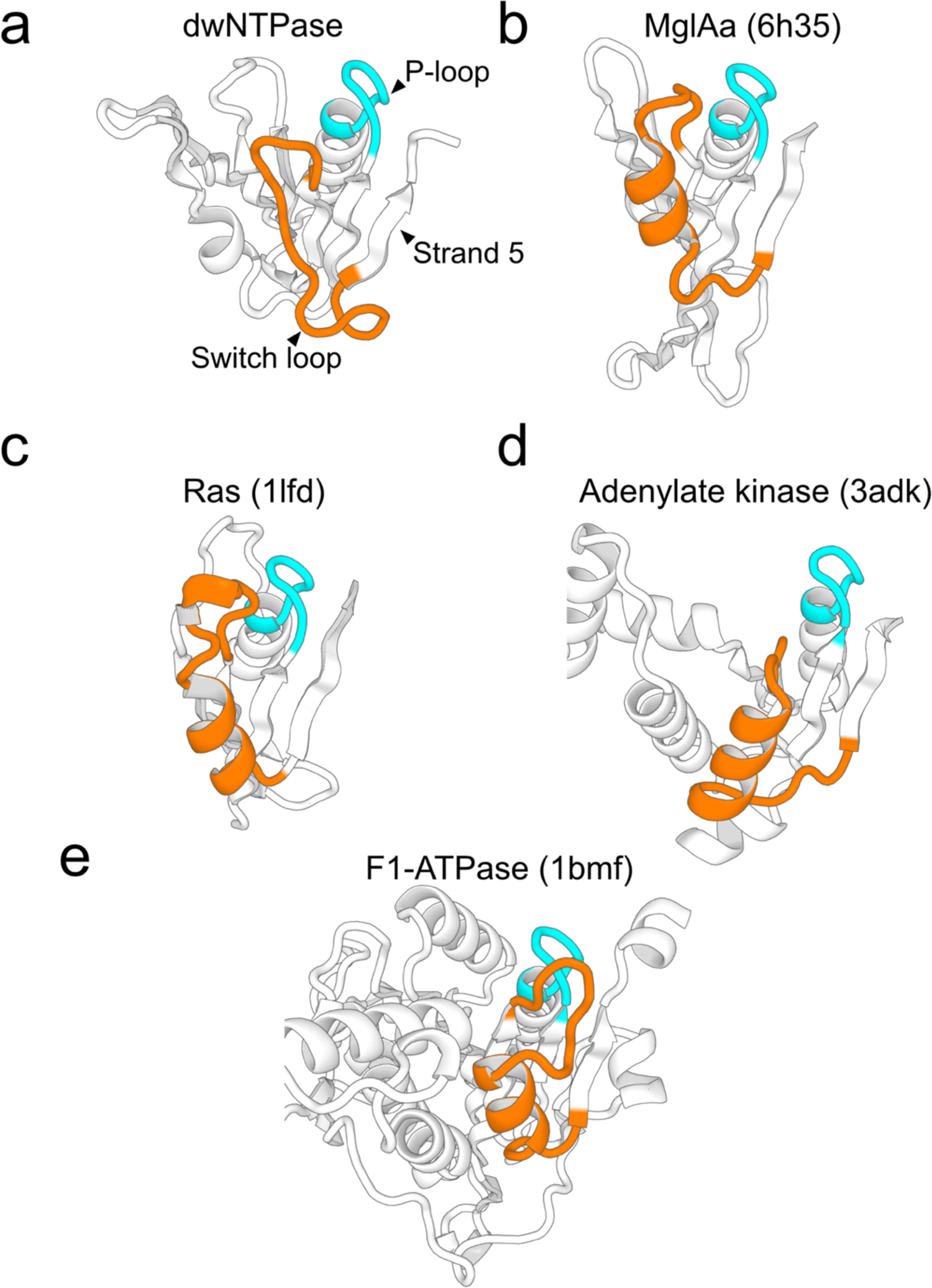
Comparison of the switch loop with the helical region conserved among P-loop NTPases. Typical P-loop NTPase structures are compared to the dwNTPase P-loop domain (**a**). MglAa (**b**), Ras (**c**), adenylate-kinase (**d**) and F1-ATPase (**e**) are selected from G-proteins (SCCS: c.37.8), NKs (c.37.1) and Rec-A like protein (c.37.11) from SCOPe. The switch loop or corresponding helical regions are orange, and the P-loops are cyan. C-terminal portions of the structures that follow strand 5 are omitted for clarity. Protein names and PDB IDs are shown above the panels.

**Figure supplement 5.**
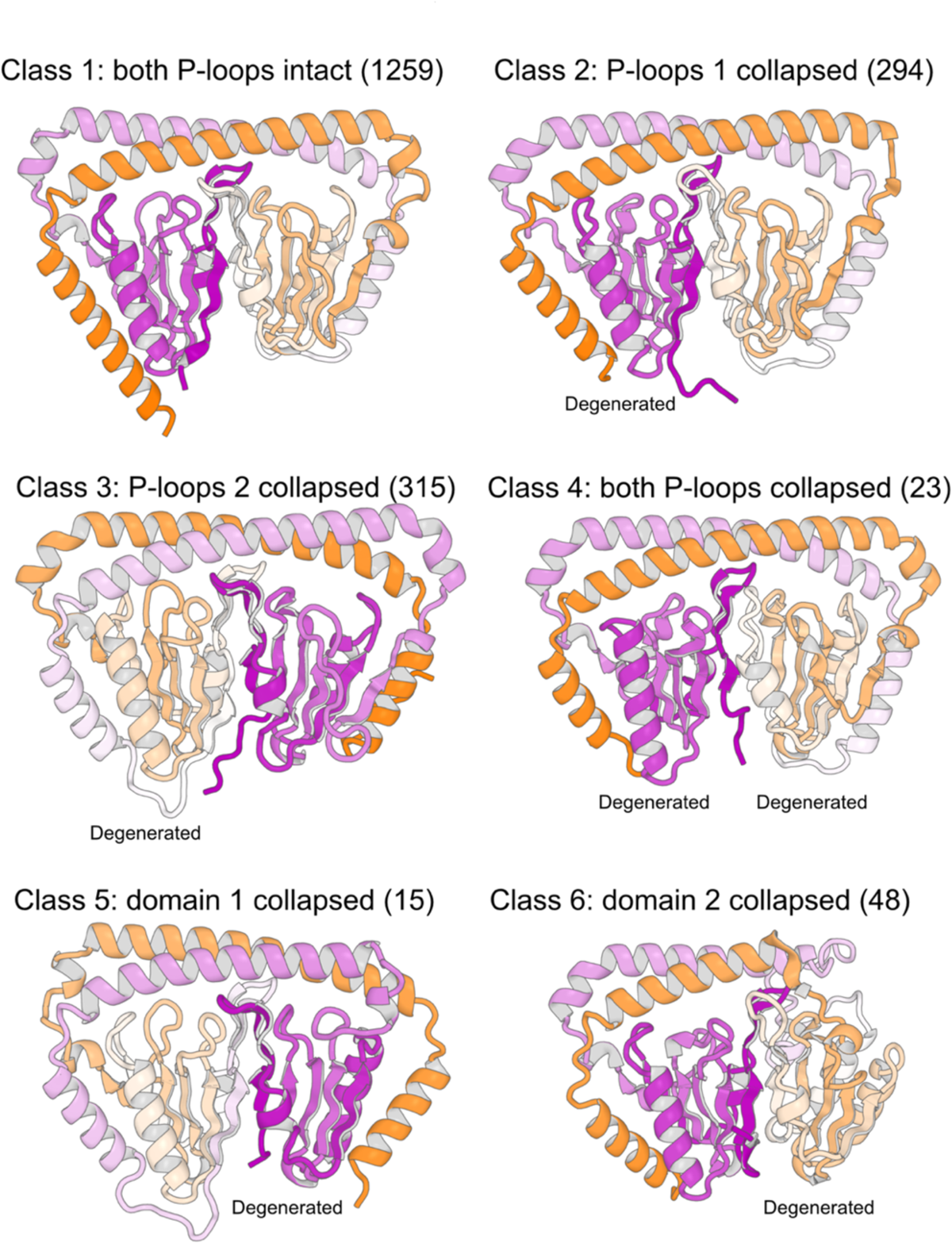
Variations in the dwNTPase structure. Depending on the conservation of P-loops and domains, the dwNTPase family can be classified into six classes. The class 1 structure (e.g., Bt. dwNTPase, Uniport accession No. A0A1Y0TWD8) preserves both P-loops intact. The class 2 structure (A0A1T4Y6S3) has P-loop 1 degenerate and P-loop 2 intact. The class 3 structure (A0A1Y4J6T6) has P-loop 1 intact and P-loop 2 degenerate. The class 4 structure (A0A1I3I7D1) has both P-loops degenerate. The class 5 structure (A0A1Y4EVI7) has domain 1 degenerate and domain 2 intact. The class 6 structure (A0A101VWI5) has domain 1 intact and domain 2 degenerate. The number of structures that belong to respective classes is shown in parentheses. The structures are colored according to a purple-white-orange gradient from the N- to C-terminus. Note that the structures were rotated to show the collapsed region in front.

**Figure supplement 6.**
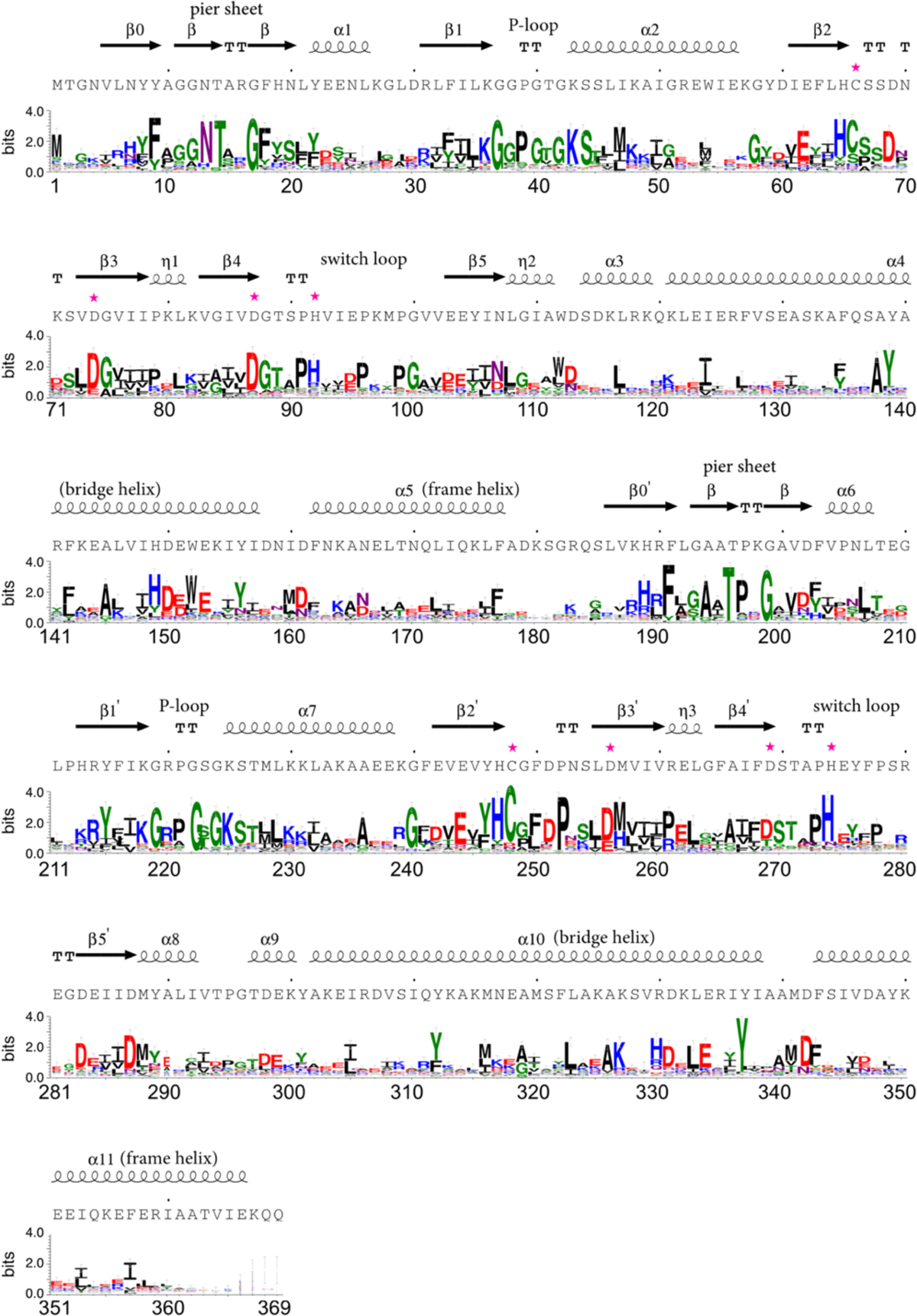
Sequence logo of dwNTPase. Using HHblits, we performed a sequence search against the UniRef30 database and constructed multiple sequence alignments. Black, blue, green and red represent hydrophobic, positively charged, polar and negatively charged residues. The amino acid sequence of the representative structure (Bt. dwNTPase) and the secondary structures assigned by DSSP are shown. The vertical axis represents the bit score, and the horizontal axis represents the residue index. Putative functional residues discussed in the main text are indicated by purple stars.

**Figure supplement 7.**
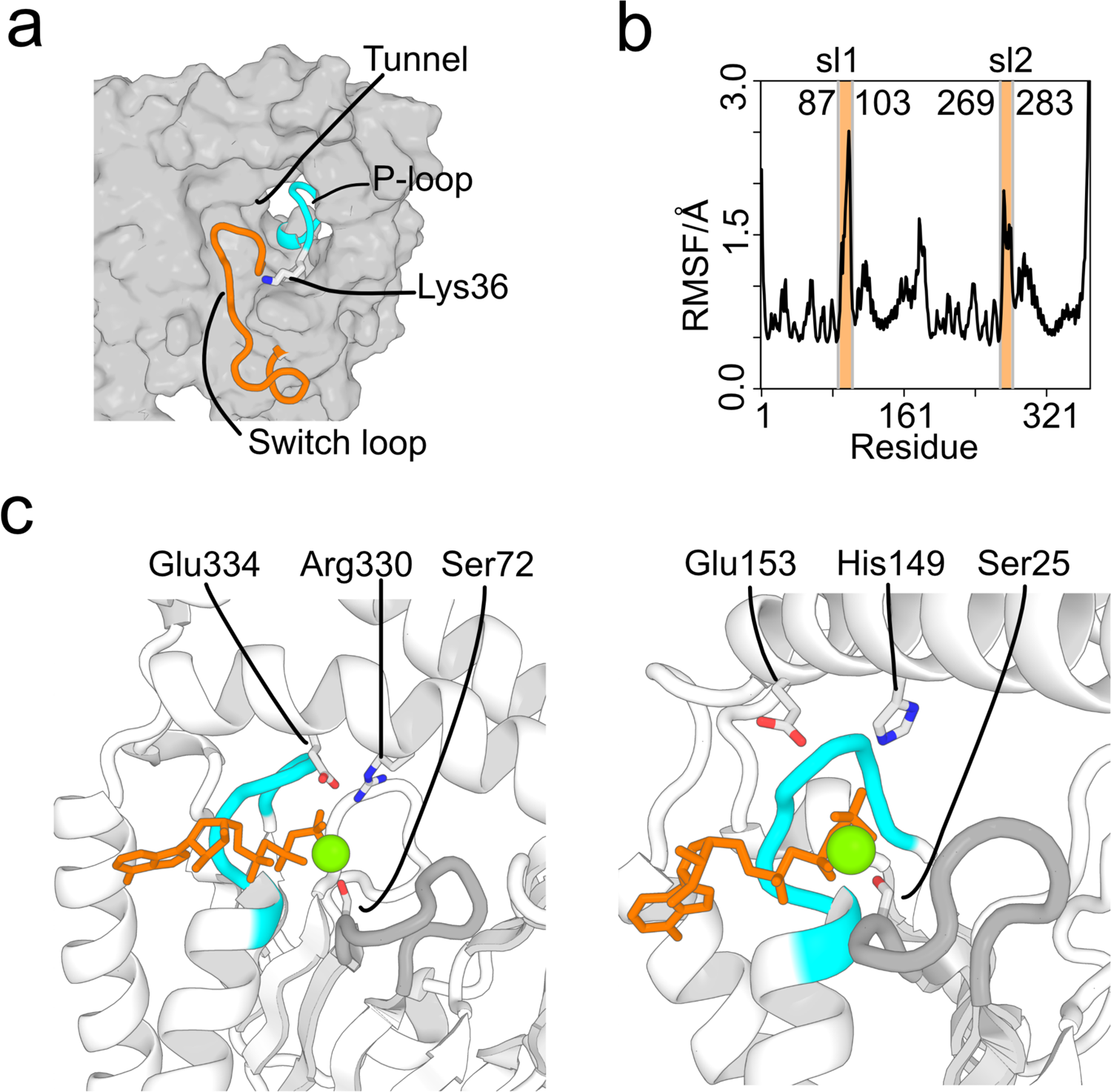
Other characteristic residues and substructures found in dwNTPases. **a**, Location of the switch loop relative to the tunnel and the hydrogen bond formed between conserved lysine and glutamate residues. The switch loop and P-loop are shown in orange and cyan cartoon, and the rest of the structure is a grey surface. The side chain of Lys36 is shown. **b**, Root-mean-square fluctuation (RMSF) of the Cα atoms during 20 trajectories of 100 ns MD simulations. The positions of switch loops are indicated by orange bars and labels reading sl1/sl2. The numbers above the plot show the residue index of the start/end of the switch loops. **c**, Conserved residues around P-loops supporting the recognition of ATP molecules. The side chain atoms of conserved residue recognizing ATPs or Mg^2+^ ions are shown as sticks. P-loops, ATPs and Mg^2+^ ions are colored cyan, orange and green, respectively. Additional loops unobserved in other P-loop proteins are colored grey.

**Figure supplement 8.**
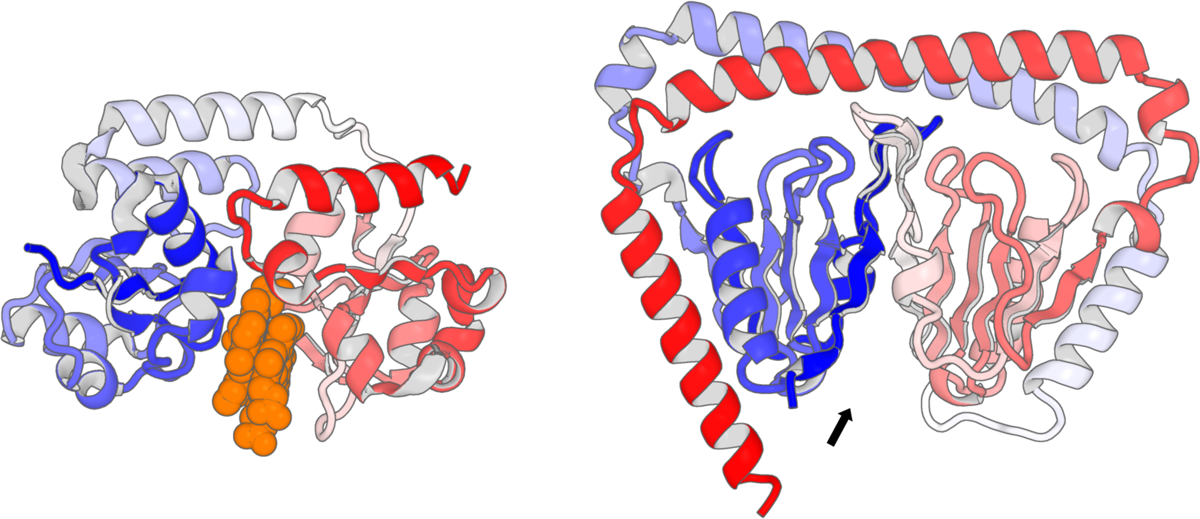
Comparison with periplasmic heme-binding proteins. (Left) A crystal structure of periplasmic heme-binding protein HmuT (PDB ID: 3nu1) liganded with two heme molecules represented in orange CPK. (Right) The structure of dwNTPase is shown at the same scale. The cleft between two P-loop domains found in dwNTPase is indicated by an arrow. The chain is shown in cartoon representation colored according to a blue-white-red gradient from the N- to C-terminus.

**Figure supplement 9.**
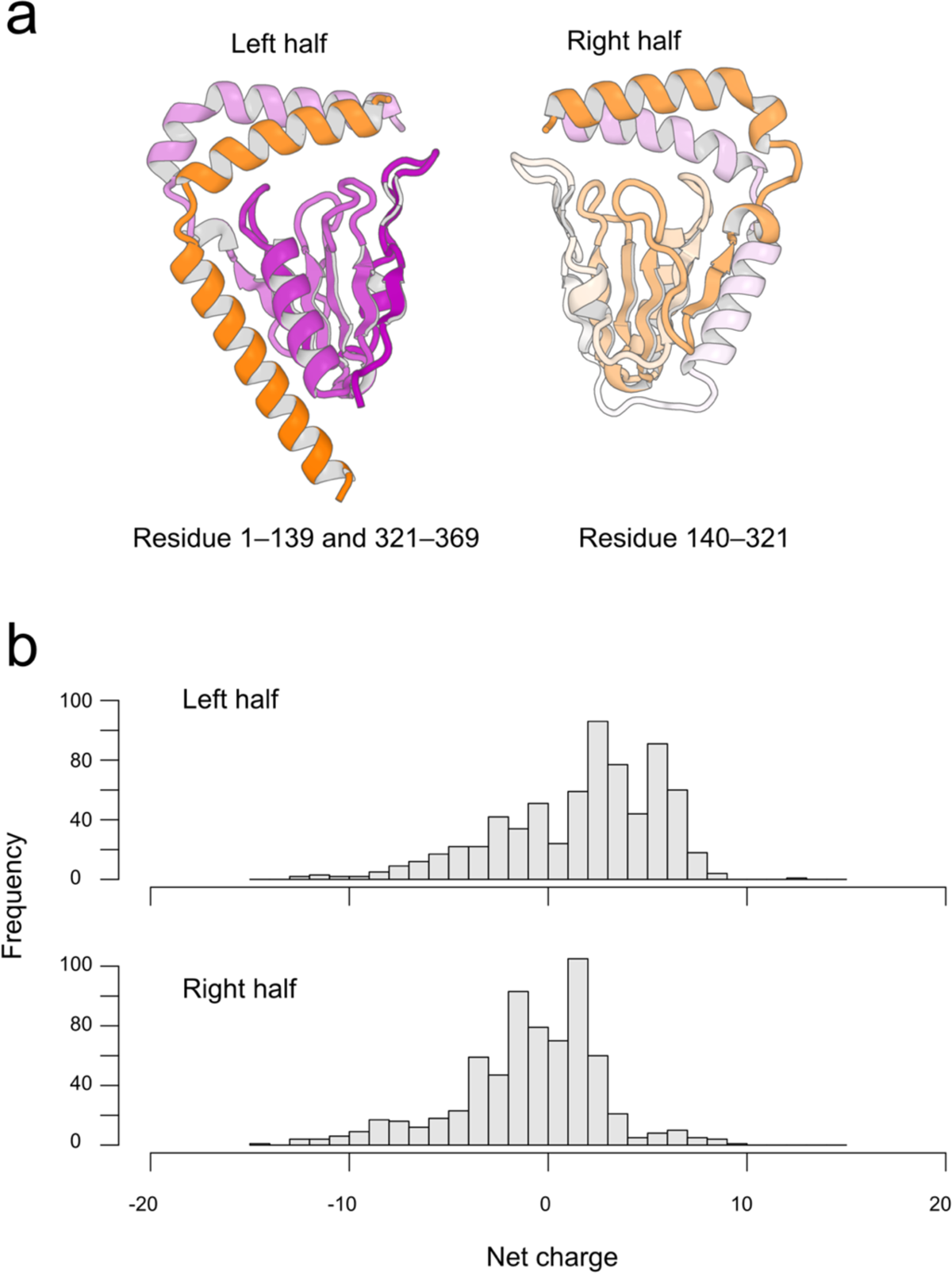
Distribution of the net charge in the left and right halves of the dwNTPase structure. **a**, The representative structure was divided into left and right halves and used as a template to find corresponding halves in other structures. The structures are colored according to a purple-white-orange gradient from the N- to C-terminus. **b**, From the initially mined 711 dwNTPase structures, 707 structures with all secondary structures intact were selected. The structures were aligned to the left half of the representative structure, and the aligned regions were assigned as left halves. The remaining regions of the structures were assigned as the right halves. In calculating net charges, each Arg, His and Lys residue was counted to have a +1 charge, and each Asp and Glu residue was counted to have a –1 charge.

## Notes

### Competing Interest Statement

The authors have declared no competing interest.

### Summary of Updates

(0) Modified abstract. (1) The main text was extended. (2) Table 1 and figure 3 were moved from extended material to main. (3) Figures 7,8 and 9 were revived.

