## Supplement Data Table1 for "Dual-wield NTPases: a novel protein family mined from AlphaFold DB"

**Supplement Data Table 1:** **List of AlphaFold DB entries structurally related to dwNTPase family.** First column stores the Uniprot accession codes of the protein, and the second column stores the organism or resource names.

| A0A011VR05 | Ruminococcus albus SY3 |
| --- | --- |
| A0A023NZU8 | Bacillus bombysepticus str. Wang |
| A0A031WEG6 | Clostridioides difficile |
| A0A037ZA99 | Clostridium tetanomorphum DSM 665 |
| A0A072NEZ6 | Bacillus azotoformans MEV2011 |
| A0A072XU63 | Clostridium botulinum C/D str. BKT12695 |
| A0A072Y634 | Clostridium sp. K25 |
| A0A072YM95 | Clostridium novyi B str. NCTC 9691 |
| A0A073KAF6 | Bacillus manliponensis |
| A0A073KAV0 | Bacillus gaemokensis |
| A0A075JVS8 | Virgibacillus sp. SK37 |
| A0A075R584 | Brevibacillus laterosporus LMG 15441 |
| A0A077J9P5 | Bacillus sp. X1(2014) |
| A0A078KVF2 | [Clostridium] cellulosi |
| A0A084IXZ6 | Bacillus mycoides |
| A0A084JBV3 | Clostridium sulfidigenes |
| A0A085LC24 | Peptococcaceae bacterium SCADC1_2_3 |
| A0A089IKM2 | Paenibacillus sp. FSL H7-0737 |
| A0A089KC26 | Paenibacillus sp. FSL R7-0273 |
| A0A089KST6 | Paenibacillus sp. FSL R7-0331 |
| A0A090ITZ3 | Caldibacillus thermoamylovorans |
| A0A090YS29 | Bacillus clarus |
| A0A096B6R2 | Flavonifractor plautii 1_3_50AFAA |
| A0A096B6U9 | Flavonifractor plautii 1_3_50AFAA |
| A0A096BFL9 | Caloranaerobacter azorensis H53214 |
| A0A098AYH1 | Desulfitobacterium hafniense |
| A0A098B810 | Desulfitobacterium hafniense |
| A0A098F568 | Bacillus sp. B-jedd |
| A0A098F754 | Peribacillus simplex |
| A0A099RZW2 | Desulfosporosinus sp. HMP52 |
| A0A099S8A8 | Clostridium sp. HMP27 |
| A0A099SC38 | Clostridium sp. HMP27 |
| A0A0A0I967 | Clostridium botulinum C/D str. DC5 |
| A0A0A0IB26 | Clostridium novyi A str. 4552 |
| A0A0A0IJB9 | Clostridium haemolyticum NCTC 8350 |
| A0A0A0IUT0 | Clostridium novyi A str. 4570 |
| A0A0A1MWX8 | Oceanobacillus oncorhynchi |
| A0A0A2TJ00 | Desulfosporosinus sp. Tol-M |
| A0A0A2TS74 | Desulfosporosinus sp. Tol-M |
| A0A0A2VHB7 | Pontibacillus chungwhensis BH030062 |
| A0A0A3I689 | Lysinibacillus manganicus DSM 26584 |
| A0A0A3IIX0 | Lysinibacillus boronitolerans JCM 21713 = 10a = NBRC 103108 |
| A0A0A3IM15 | Lysinibacillus sinduriensis BLB-1 = JCM 15800 |
| A0A0A3J5T7 | Lysinibacillus massiliensis 4400831 = CIP 108448 = CCUG 49529 |
| A0A0A5GE29 | Pontibacillus marinus BH030004 = DSM 16465 |
| A0A0A7FXV3 | Clostridium baratii str. Sullivan |
| A0A0A8JGW3 | Bacillus sp. (strain OxB-1) |
| A0A0A8X6A0 | Bacillus selenatarsenatis SF-1 |
| A0A0B0HMM4 | Paenibacillus sp. P1XP2 |
| A0A0B0HQQ8 | Paenibacillus sp. P1XP2 |
| A0A0B0IK09 | Alkalihalobacillus okhensis |
| A0A0B1Y1A2 | Lysinibacillus sp. A1 |
| A0A0B3VZH0 | Terrisporobacter othiniensis |
| A0A0B4WCA3 | Clostridium botulinum Prevot_594 |
| A0A0B5X8Z0 | Bacillus thuringiensis |
| A0A0B7MFU2 | Syntrophaceticus schinkii |
| A0A0B7MIF4 | Syntrophaceticus schinkii |
| A0A0B7MJC7 | Syntrophaceticus schinkii |
| A0A0C1UFI3 | Clostridium argentinense CDC 2741 |
| A0A0C2RR63 | Jeotgalibacillus campisalis |
| A0A0C2TRM5 | Bacillus badius |
| A0A0C2UQT9 | Cohnella kolymensis |
| A0A0C2VZ26 | Jeotgalibacillus soli |
| A0A0C3HBH1 | Clostridium botulinum |
| A0A0C5BZN8 | Weizmannia coagulans |
| A0A0C7GBA6 | Paeniclostridium sordellii |
| A0A0C7N6A9 | Moorella glycerini |
| A0A0C7NLI7 | Moorella glycerini |
| A0A0C7NNC1 | Moorella glycerini |
| A0A0D0EGA2 | Caldibacillus thermoamylovorans |
| A0A0D0FPH4 | Caldibacillus thermoamylovorans |
| A0A0D0QXF9 | Bacillus sp. L_1B0_5 |
| A0A0D1BPS2 | Clostridium botulinum B2 450 |
| A0A0D1R4D3 | Bacillus thuringiensis Sbt003 |
| A0A0D3VI41 | Paenibacillus sp. IHBB 10380 |
| A0A0D7WTV6 | Paenibacillus terrae |
| A0A0D8I5Y8 | Clostridium aceticum |
| A0A0D8J352 | Ruthenibacterium lactatiformans |
| A0A0E1L182 | Clostridium botulinum CDC_1436 |
| A0A0E1MKX4 | Bacillus cereus |
| A0A0E3JRD0 | Clostridium scatologenes |
| A0A0E3W385 | Syntrophomonas zehnderi OL-4 |
| A0A0E4GWT5 | Syntrophomonas zehnderi OL-4 |
| A0A0F2JK31 | Desulfosporosinus sp. I2 |
| A0A0F2NB27 | Peptococcaceae bacterium BRH_c4a |
| A0A0F2PJY4 | Peptococcaceae bacterium BRH_c8a |
| A0A0F2PRR1 | Peptococcaceae bacterium BRH_c4b |
| A0A0F2Q015 | Clostridiaceae bacterium BRH_c20a |
| A0A0F2S8L7 | Peptococcaceae bacterium BRH_c23 |
| A0A0F2SMQ4 | Peptococcaceae bacterium BRH_c23 |
| A0A0F3FS45 | Clostridium baratii |
| A0A0F5I727 | Quasibacillus thermotolerans |
| A0A0F5RKY0 | Bacillus sp. UMTAT18 |
| A0A0F6G0J2 | Bacillus thuringiensis subsp. kurstaki |
| A0A0F6Y0I8 | Brevibacillus laterosporus |
| A0A0F7RJU5 | Bacillus anthracis |
| A0A0G8C3G0 | Bacillus wiedmannii |
| A0A0G8E5A0 | Bacillus cereus |
| A0A0G8F8F2 | Bacillus cereus |
| A0A0G9LEG5 | Clostridium sp. C8 |
| A0A0H2YUK9 | Clostridium perfringens (strain ATCC 13124 / DSM 756 / JCM 1290 / NCIMB 6125 / NCTC 8237 / Type A) |
| A0A0H3NFU8 | Clostridioides difficile (strain CD196) |
| A0A0H5SV22 | Herbinix hemicellulosilytica |
| A0A0J1DLS7 | Peptococcaceae bacterium 1109 |
| A0A0J1FUB6 | Desulfosporosinus acididurans |
| A0A0J1HRD9 | Bacillus anthracis |
| A0A0J1IJI5 | Desulfosporosinus acididurans |
| A0A0J5GZ52 | Bacillus sp. LL01 |
| A0A0J5VYY5 | Cytobacillus firmus |
| A0A0J6L130 | Bacillus sp. LK2 |
| A0A0J6Z9S3 | Bacillus cereus |
| A0A0J7DES4 | Bacillus cereus |
| A0A0J7GWC3 | Bacillus cereus |
| A0A0J8D4K4 | Clostridium cylindrosporum DSM 605 |
| A0A0K0GAZ8 | Bacilli bacterium VT-13-104 |
| A0A0K0Q6V2 | Bacillus thuringiensis |
| A0A0K8J594 | Herbinix luporum |
| A0A0K9F4F4 | Lysinibacillus xylanilyticus |
| A0A0K9GZ84 | Peribacillus loiseleuriae |
| A0A0K9H9Y9 | Bacillus sp. FJAT-27231 |
| A0A0K9MEL7 | Bacillus sp. FJAT-27238 |
| A0A0K9YXI0 | Brevibacillus reuszeri |
| A0A0L0WD40 | Gottschalkia purinilytica |
| A0A0L6JUT0 | Pseudobacteroides cellulosolvens ATCC 35603 = DSM 2933 |
| A0A0L6VZ10 | Thermincola ferriacetica |
| A0A0L6ZAH9 | Clostridium homopropionicum DSM 5847 |
| A0A0L7NS99 | Clostridium botulinum |
| A0A0L9YAX5 | Clostridium botulinum |
| A0A0M0G936 | Sporosarcina globispora |
| A0A0M0GPI9 | Bacillus marisflavi |
| A0A0M0LLZ8 | Viridibacillus arvi |
| A0A0M0WCR2 | Bacillus sp. FJAT-21945 |
| A0A0M0WVJ1 | Lysinibacillus sp. FJAT-14745 |
| A0A0M1IYQ9 | Clostridium sp. L74 |
| A0A0M1N1C9 | Paenibacillus solani |
| A0A0M1NSP8 | Bacillus sp. FJAT-22058 |
| A0A0M1URX7 | Paeniclostridium sordellii |
| A0A0M2PCR3 | Bacillus sp. SA1-12 |
| A0A0M2SZL0 | Mesobacillus campisalis |
| A0A0M2U5Z7 | Clostridiales bacterium PH28_bin88 |
| A0A0M2VWH4 | Paenibacillus sp. DMB20 |
| A0A0M3DJ91 | Paraclostridium benzoelyticum |
| A0A0M4G2T7 | Bacillus sp. FJAT-18017 |
| A0A0M4GHP4 | Bacillus sp. FJAT-22090 |
| A0A0M8PRN4 | Lysinibacillus sp. FJAT-14222 |
| A0A0M9GSA4 | Bacillus sp. CHD6a |
| A0A0M9WYR3 | Lysinibacillus contaminans |
| A0A0N0CX44 | Lysinibacillus macroides |
| A0A0N0M8S6 | Oceanobacillus caeni |
| A0A0N1HD10 | Clostridioides difficile |
| A0A0P6VY21 | Rossellomorea vietnamensis |
| A0A0P7JZU5 | Lysinibacillus sp. ZYM-1 |
| A0A0P8VTG0 | Caloranaerobacter sp. TR13 |
| A0A0Q0R9D7 | Bacillus thuringiensis |
| A0A0Q3QUY6 | Cytobacillus solani |
| A0A0Q3S4L7 | Psychrobacillus sp. FJAT-21963 |
| A0A0Q3T811 | Brevibacillus choshinensis |
| A0A0Q3W0U6 | Bacillus sp. FJAT-25509 |
| A0A0Q3WSD7 | Bacillus shackletonii |
| A0A0Q6KXT2 | Bacillus sp. Leaf406 |
| A0A0Q9VYZ9 | Bacillus sp. Soil768D1 |
| A0A0Q9Y4M1 | Virgibacillus soli |
| A0A0R3K1I6 | Caloramator mitchellensis |
| A0A0S2W210 | Intestinimonas butyriciproducens |
| A0A0S6U621 | Clostridium botulinum B str. Osaka05 |
| A0A0S6U9Q5 | Moorella thermoacetica Y72 |
| A0A0S6UH05 | Moorella thermoacetica Y72 |
| A0A0U1KNY7 | Paraliobacillus sp. PM-2 |
| A0A0U1L3G4 | Sporomusa ovata |
| A0A0U1NX13 | Neobacillus massiliamazoniensis |
| A0A0U2UAW9 | Paenibacillus naphthalenovorans |
| A0A0U2UQH7 | Paenibacillus sp. 32O-W |
| A0A0U3WB32 | Lentibacillus amyloliquefaciens |
| A0A0U9H622 | Oceanobacillus picturae |
| A0A0V8HJY0 | Bacillus enclensis |
| A0A0W1AXZ5 | Paenibacillus etheri |
| A0A0W1JHD8 | Desulfitobacterium hafniense |
| A0A0W1JPB7 | Desulfitobacterium hafniense |
| A0A0W7TVM7 | Ruthenibacterium lactatiformans |
| A0A0W7YLD2 | Lysinibacillus sp. F5 |
| A0A0W8E4K5 | hydrocarbon metagenome |
| A0A0W8E642 | hydrocarbon metagenome |
| A0A0X8D2G5 | Aneurinibacillus sp. XH2 |
| A0A0X8G2A8 | Turicibacter sp. H121 |
| A0A101F7Q9 | Thermoanaerobacterales bacterium 50_218 |
| A0A101FQZ0 | Clostridia bacterium 62_21 |
| A0A101FR57 | Clostridia bacterium 62_21 |
| A0A101GVR0 | Desulfotomaculum sp. 46_80 |
| A0A101HT37 | Pelotomaculum thermopropionicum |
| A0A101V9B3 | Desulfitibacter sp. BRH_c19 |
| A0A101VTK5 | Gracilibacter sp. BRH_c7a |
| A0A101VWI5 | Gracilibacter sp. BRH_c7a |
| A0A101WDA9 | Desulfosporosinus sp. BRH_c37 |
| A0A101Y1Q0 | Paenibacillus sp. DMB5 |
| A0A109G0D1 | Bacillus mycoides |
| A0A117KXD0 | Clostridia bacterium 41_269 |
| A0A117S2R7 | Desulfitibacter sp. BRH_c19 |
| A0A120GQ95 | Peribacillus simplex |
| A0A125YDJ8 | Clostridioides difficile ATCC 9689 = DSM 1296 |
| A0A127DDQ3 | Peribacillus simplex |
| A0A127EJE1 | Clostridium perfringens |
| A0A127W314 | Sporosarcina psychrophila |
| A0A133KH74 | Weizmannia coagulans |
| A0A133MV32 | Clostridium perfringens |
| A0A135L4N3 | Tepidibacillus decaturensis |
| A0A135WG74 | Sporosarcina sp. HYO08 |
| A0A136BHT8 | Bacillus cereus |
| A0A140L102 | Fervidicola ferrireducens |
| A0A140LB51 | Thermotalea metallivorans |
| A0A143HCD7 | Rummeliibacillus stabekisii |
| A0A143ZUA6 | Eubacteriaceae bacterium CHKCI005 |
| A0A150BFB0 | Bacillus cereus |
| A0A150BLI9 | Bacillus cereus |
| A0A150D114 | Bacillus cereus |
| A0A150E2M5 | Bacillus cereus |
| A0A150EXW3 | Bacillus cereus |
| A0A150FRV8 | [Clostridium] paradoxum JW-YL-7 = DSM 7308 |
| A0A150JV47 | Weizmannia coagulans |
| A0A150JY23 | Weizmannia coagulans |
| A0A150KAW3 | Weizmannia coagulans |
| A0A150L6D1 | Bacillus sporothermodurans |
| A0A151AP40 | Clostridium colicanis DSM 13634 |
| A0A151AXY0 | Moorella mulderi DSM 14980 |
| A0A151AZJ8 | Moorella mulderi DSM 14980 |
| A0A151B040 | Clostridium tepidiprofundi DSM 19306 |
| A0A151UTC6 | Bacillus cereus |
| A0A154BVE5 | Anaerosporomusa subterranea |
| A0A158RNF5 | Bacillus cereus (strain 03BB102) |
| A0A160H875 | Bacillus cereus |
| A0A160IK67 | Fictibacillus phosphorivorans |
| A0A160MHH3 | Bacillus oceanisediminis 2691 |
| A0A161QQ05 | Bacillus cereus |
| A0A161RGG3 | Bhargavaea cecembensis |
| A0A161XFK1 | Clostridium magnum DSM 2767 |
| A0A162M8B1 | Thermovenabulum gondwanense |
| A0A162U197 | Clostridium magnum DSM 2767 |
| A0A163QB21 | Fictibacillus phosphorivorans |
| A0A164MFU9 | Bacillus cereus |
| A0A165J5J0 | Bacillus marisflavi |
| A0A167ASF5 | Paenibacillus crassostreae |
| A0A168DKC2 | Paenibacillus macquariensis |
| A0A168KZD2 | Paenibacillus antarcticus |
| A0A169ZHB1 | Paenibacillus glacialis |
| A0A173QZX0 | Turicibacter sanguinis |
| A0A173RAA6 | Faecalibacterium prausnitzii |
| A0A173WID7 | Turicibacter sanguinis |
| A0A173WRV5 | Clostridium ventriculi |
| A0A173Z3K4 | Clostridium disporicum |
| A0A174BH00 | Faecalibacterium prausnitzii |
| A0A174FC78 | Clostridium paraputrificum |
| A0A174GRL5 | Clostridium disporicum |
| A0A174MRJ7 | Anaerotruncus colihominis |
| A0A174NG55 | Anaerotruncus colihominis |
| A0A174SS44 | Clostridium baratii |
| A0A174VYZ3 | Flavonifractor plautii |
| A0A174WPP5 | Flavonifractor plautii |
| A0A175LR66 | Clostridium botulinum B2 433 |
| A0A177KLS5 | Domibacillus aminovorans |
| A0A177KZN5 | Domibacillus aminovorans |
| A0A177XSF6 | Brevibacillus sp. SKDU10 |
| A0A177ZJ27 | Lederbergia galactosidilyticus |
| A0A179T4F9 | Metabacillus litoralis |
| A0A193CMG2 | Bacillus thuringiensis serovar coreanensis |
| A0A1A5X1Q3 | Brevibacillus sp. WF146 |
| A0A1A5YUR3 | Paenibacillus oryzae |
| A0A1B1KZS3 | Bacillus thuringiensis |
| A0A1B1YDN0 | Thermoclostridium stercorarium subsp. thermolacticum DSM 2910 |
| A0A1B1YKT7 | Thermoclostridium stercorarium subsp. leptospartum DSM 9219 |
| A0A1B1YZX8 | Fictibacillus arsenicus |
| A0A1B2E1A4 | Paenibacillus ihbetae |
| A0A1B3XVP1 | Peribacillus muralis |
| A0A1B7LBR9 | Desulfotomaculum copahuensis |
| A0A1B7LND2 | Candidatus Arthromitus sp. SFB-turkey |
| A0A1B8WEU6 | Bacillus sp. FJAT-27264 |
| A0A1B8WIC8 | Bacillus sp. FJAT-26390 |
| A0A1B9AER0 | Bacillus sp. FJAT-27225 |
| A0A1B9AYB3 | Bacillus wudalianchiensis |
| A0A1C0AAB8 | Orenia metallireducens |
| A0A1C2XR73 | Dehalobacter sp. TeCB1 |
| A0A1C2Y2N9 | Dehalobacter sp. TeCB1 |
| A0A1C3F700 | Desulfosporosinus sp. BG |
| A0A1C3T2Z1 | Bacillus mycoides |
| A0A1C3ZGP9 | Bacillus cereus |
| A0A1C3ZK95 | Bacillus mycoides |
| A0A1C3ZL92 | Bacillus wiedmannii |
| A0A1C3ZNB1 | Bacillus thuringiensis |
| A0A1C5L3M5 | uncultured Ruminococcus sp |
| A0A1C5LVQ8 | uncultured Faecalibacterium sp |
| A0A1C5PDC5 | uncultured Oscillibacter sp |
| A0A1C5PER6 | uncultured Clostridium sp |
| A0A1C5PYK2 | uncultured Faecalibacterium sp |
| A0A1C5SL51 | uncultured Eubacterium sp |
| A0A1C5URW7 | uncultured Clostridium sp |
| A0A1C5W3J4 | uncultured Ruminococcus sp |
| A0A1C5WEL9 | uncultured Clostridium sp |
| A0A1C6AI66 | uncultured Flavonifractor sp |
| A0A1C6AIX5 | uncultured Flavonifractor sp |
| A0A1C6BAZ6 | uncultured Flavonifractor sp |
| A0A1C6BC28 | uncultured Flavonifractor sp |
| A0A1C6BD87 | uncultured Ruminococcus sp |
| A0A1C6BJ78 | uncultured Clostridium sp |
| A0A1C6BPE7 | uncultured Flavonifractor sp |
| A0A1C6C6A7 | uncultured Ruminococcus sp |
| A0A1C6C7X7 | uncultured Clostridium sp |
| A0A1C6C8D7 | uncultured Ruminococcus sp |
| A0A1C6CCN1 | uncultured Clostridium sp |
| A0A1C6D0U7 | uncultured Ruminococcus sp |
| A0A1C6F277 | uncultured Clostridium sp |
| A0A1C6FK51 | uncultured Clostridium sp |
| A0A1C6FLL0 | uncultured Anaerotruncus sp |
| A0A1C6FLY0 | uncultured Clostridium sp |
| A0A1C6H0S5 | uncultured Flavonifractor sp |
| A0A1C6HFU5 | uncultured Flavonifractor sp |
| A0A1C6HG51 | uncultured Flavonifractor sp |
| A0A1C6I316 | uncultured Ruminococcus sp |
| A0A1C6IIH8 | uncultured Oscillibacter sp |
| A0A1C6JLW8 | uncultured Eubacterium sp |
| A0A1C6W9W4 | Bacillus wiedmannii |
| A0A1C7FJT2 | Flavonifractor plautii |
| A0A1C9BMK6 | Bacillus thuringiensis Bt18247 |
| A0A1D3N109 | Bacillus mycoides |
| A0A1D3QAF9 | Bacillus cereus |
| A0A1D7XAR7 | Moorella thermoacetica |
| A0A1D7XC76 | Moorella thermoacetica |
| A0A1D7XN80 | Clostridium taeniosporum |
| A0A1D8GCD5 | Geosporobacter ferrireducens |
| A0A1D8GCE3 | Geosporobacter ferrireducens |
| A0A1D8JJK0 | Sporosarcina ureilytica |
| A0A1D9FQP5 | Clostridium formicaceticum |
| A0A1E3BXQ2 | Clostridium sp. Bc-iso-3 |
| A0A1E4LH44 | Clostridium sp. SCN 57-10 |
| A0A1E4R2F3 | Lysinibacillus fusiformis |
| A0A1E5G5R6 | Desulfuribacillus alkaliarsenatis |
| A0A1E5K3E5 | Oceanobacillus sp. E9 |
| A0A1E8BDF3 | Bacillus mycoides |
| A0A1E8BV13 | Bacillus mycoides |
| A0A1E8F1G9 | Clostridium acetireducens DSM 10703 |
| A0A1F8U479 | Clostridiales bacterium GWB2_37_7 |
| A0A1F8UM10 | Clostridiales bacterium GWD2_32_59 |
| A0A1G4EEX6 | Bacillus mycoides |
| A0A1G5C0U1 | Alkaliphilus peptidifermentans DSM 18978 |
| A0A1G5HY52 | Desulfoluna spongiiphila |
| A0A1G6DDP0 | Ruminococcaceae bacterium FB2012 |
| A0A1G6KYH8 | Pelagirhabdus alkalitolerans |
| A0A1G7DNX2 | Bhargavaea beijingensis |
| A0A1G7MCD1 | Sporolituus thermophilus DSM 23256 |
| A0A1G7TAP9 | Fontibacillus panacisegetis |
| A0A1G7USJ3 | Desulfosporosinus hippei DSM 8344 |
| A0A1G7Y1D2 | Aneurinibacillus thermoaerophilus |
| A0A1G8GQA3 | Alteribacillus persepolensis |
| A0A1G8MRA4 | Alteribacillus bidgolensis |
| A0A1G8RPF8 | Natribacillus halophilus |
| A0A1G9AD17 | Paenibacillus typhae |
| A0A1G9BBN1 | Natronincola ferrireducens |
| A0A1G9FM54 | Clostridium cochlearium |
| A0A1G9RMI9 | Romboutsia lituseburensis DSM 797 |
| A0A1G9WXA1 | Tenuibacillus multivorans |
| A0A1G9XUL1 | Bacillus sp. OK048 |
| A0A1H0A0L6 | Psychrobacillus sp. OK028 |
| A0A1H0AAX9 | Acetanaerobacterium elongatum |
| A0A1H0Q2G8 | Clostridium gasigenes |
| A0A1H0TBK8 | Litchfieldia salsus |
| A0A1H0ZBE8 | Virgibacillus salinus |
| A0A1H2T661 | Tepidimicrobium xylanilyticum |
| A0A1H3QW75 | Bacillus sp. 166amftsu |
| A0A1H3RSI3 | Proteiniborus ethanoligenes |
| A0A1H3V0B1 | Evansella caseinilytica |
| A0A1H3XM84 | Thalassobacillus cyri |
| A0A1H5TB50 | Caloramator fervidus |
| A0A1H6B9T0 | Bacillus sp. ok061 |
| A0A1H6TDH0 | Bhargavaea ginsengi |
| A0A1H7MK05 | Ruminococcus albus |
| A0A1H7N6W8 | Paenibacillus sp. cl141a |
| A0A1H7YNT9 | Hydrogenoanaerobacterium saccharovorans |
| A0A1H7YQC1 | Hydrogenoanaerobacterium saccharovorans |
| A0A1H8BB33 | Candidatus Frackibacter sp. WG12 |
| A0A1H8GM21 | Mesobacillus persicus |
| A0A1H9E2K8 | Piscibacillus halophilus |
| A0A1H9HI04 | Lysinibacillus fusiformis |
| A0A1H9TG16 | Salipaludibacillus aurantiacus |
| A0A1H9TZI7 | Psychrobacillus sp. OK032 |
| A0A1I0BNV3 | Oceanobacillus limi |
| A0A1I0CFN1 | Anaerobranca gottschalkii DSM 13577 |
| A0A1I0CQ85 | Salinibacillus kushneri |
| A0A1I0DKG8 | [Clostridium] polysaccharolyticum |
| A0A1I0EAX8 | Natronincola peptidivorans |
| A0A1I0WGY8 | Lentibacillus halodurans |
| A0A1I0ZY24 | Clostridium frigidicarnis |
| A0A1I1H869 | Bacillus sp. OV322 |
| A0A1I1PB18 | Clostridium uliginosum |
| A0A1I1PZW9 | Ruminococcus albus |
| A0A1I1Q201 | Bacillus sp. 491mf |
| A0A1I1UFT1 | Bacillus sp. OV194 |
| A0A1I1WPY2 | Lentibacillus persicus |
| A0A1I2CR33 | Alteribacillus iranensis |
| A0A1I2EJD4 | Bacillus sp. OV194 |
| A0A1I2N521 | Desulfotomaculum arcticum DSM 17038 |
| A0A1I3I7D1 | Ruminococcaceae bacterium D5 |
| A0A1I3L7E6 | Brevibacillus centrosporus |
| A0A1I3S7X9 | Terrisporobacter glycolicus |
| A0A1I4LC20 | Bacillus sp. 5mfcol3.1 |
| A0A1I4PFL3 | Paenibacillus sp. 1_12 |
| A0A1I5HXL5 | Anaerocolumna aminovalerica |
| A0A1I5ML01 | Oscillibacter sp. PC13 |
| A0A1I5XIS3 | Psychrobacillus psychrotolerans |
| A0A1I5Y3L6 | Caldicoprobacter faecalis |
| A0A1I6C6U5 | Bacillus sp. cl95 |
| A0A1I6DPK0 | Desulfallas geothermicus DSM 3669 |
| A0A1I6SB48 | Halolactibacillus miurensis |
| A0A1I6W0X2 | Bacillus sp. 103mf |
| A0A1I7JA14 | Clostridium sp. DSM 8431 |
| A0A1J1D1X8 | Clostridium sporogenes |
| A0A1J5NNT6 | Moorella thermoacetica |
| A0A1J5NQY3 | Moorella thermoacetica |
| A0A1J6WUG5 | Bacillus aquimaris |
| A0A1J9UTK0 | Bacillus albus |
| A0A1J9VXU3 | Bacillus paramycoides |
| A0A1J9WCS4 | Bacillus anthracis |
| A0A1J9XDM1 | Bacillus cereus |
| A0A1K1NBX6 | Ruminococcus sp. YE71 |
| A0A1K1NXA5 | Paenibacillus sp. UNCCL117 |
| A0A1L3MR54 | Bacillus weihaiensis |
| A0A1L3NEL7 | Clostridium sporogenes |
| A0A1M2UH20 | Bacillus cereus |
| A0A1M2ZNC5 | Clostridiales bacterium 43-6 |
| A0A1M4M7T2 | Proteiniborus sp. DW1 |
| A0A1M4NBF5 | Clostridium sp. N3C |
| A0A1M4S8A9 | Tissierella praeacuta DSM 18095 |
| A0A1M4VAW7 | Caloramator proteoclasticus DSM 10124 |
| A0A1M4XQW6 | Clostridium fallax |
| A0A1M4ZT88 | Desulfotomaculum putei DSM 12395 |
| A0A1M5BA76 | Caloramator proteoclasticus DSM 10124 |
| A0A1M5BNT5 | Desulfofundulus australicus DSM 11792 |
| A0A1M5CIN4 | Caldanaerobius fijiensis DSM 17918 |
| A0A1M5FEF6 | Ornithinibacillus halophilus |
| A0A1M5KLM9 | Thermosyntropha lipolytica DSM 11003 |
| A0A1M5R855 | Desulfosporosinus lacus DSM 15449 |
| A0A1M5RDV9 | Asaccharospora irregularis DSM 2635 |
| A0A1M5RTW9 | Thermosyntropha lipolytica DSM 11003 |
| A0A1M5T741 | Tepidibacter thalassicus DSM 15285 |
| A0A1M5UUA6 | Sporanaerobacter acetigenes DSM 13106 |
| A0A1M5VU38 | Caloranaerobacter azorensis DSM 13643 |
| A0A1M5W4Z4 | Desulfosporosinus lacus DSM 15449 |
| A0A1M5XVW7 | Clostridium collagenovorans DSM 3089 |
| A0A1M6E8G4 | Clostridium intestinale DSM 6191 |
| A0A1M6EWW6 | Lutispora thermophila DSM 19022 |
| A0A1M6JJI5 | Clostridium amylolyticum |
| A0A1M6JJZ0 | Desulfofundulus thermosubterraneus DSM 16057 |
| A0A1M6KEF1 | Thermoclostridium caenicola |
| A0A1M6LSB9 | Caminicella sporogenes DSM 14501 |
| A0A1M6M5J8 | Paramaledivibacter caminithermalis DSM 15212 |
| A0A1M6MFZ1 | Tepidibacter formicigenes DSM 15518 |
| A0A1M6MLV0 | Anaerobranca californiensis DSM 14826 |
| A0A1M6N6A3 | Geosporobacter subterraneus DSM 17957 |
| A0A1M6Q1U2 | Geosporobacter subterraneus DSM 17957 |
| A0A1M6QUV4 | Clostridium cavendishii DSM 21758 |
| A0A1M6RTD8 | Hathewaya proteolytica DSM 3090 |
| A0A1M6UIB8 | Desulfotomaculum aeronauticum DSM 10349 |
| A0A1M7HWB5 | Anaerosporobacter mobilis DSM 15930 |
| A0A1M7LCH2 | Caldanaerovirga acetigignens |
| A0A1M7TBT0 | Desulfitobacterium chlororespirans DSM 11544 |
| A0A1M7UMW6 | Desulfitobacterium chlororespirans DSM 11544 |
| A0A1N7ADS6 | Peribacillus simplex |
| A0A1N7CX82 | Paenibacillus macquariensis |
| A0A1N7E780 | Bacillus cereus |
| A0A1Q5P5M7 | Domibacillus mangrovi |
| A0A1Q6KNQ7 | Clostridiales bacterium 52_15 |
| A0A1Q6QCJ8 | Firmicutes bacterium CAG:129_59_24 |
| A0A1Q6RC60 | Firmicutes bacterium CAG:24053_14 |
| A0A1Q6RFB9 | Oscillibacter sp. 57_20 |
| A0A1Q6RFH2 | Oscillibacter sp. 57_20 |
| A0A1Q6SIP3 | Firmicutes bacterium CAG:272_52_7 |
| A0A1Q6TI18 | Ruminococcus sp. 37_24 |
| A0A1Q8Q1G0 | Domibacillus antri |
| A0A1Q8R323 | Desulfosporosinus sp. OL |
| A0A1Q8UX18 | Alkalihalobacillus pseudofirmus |
| A0A1Q9I3V8 | Bacillus cereus |
| A0A1Q9PPU2 | Alkalihalobacillus pseudofirmus |
| A0A1Q9Q3V9 | Bacillus sp. MRMR6 |
| A0A1R0WYB3 | Paenibacillus odorifer |
| A0A1R0XT94 | Paenibacillus odorifer |
| A0A1R0Y423 | Paenibacillus odorifer |
| A0A1R0YXC6 | Paenibacillus odorifer |
| A0A1R0ZAD2 | Paenibacillus odorifer |
| A0A1R1A975 | Paenibacillus lautus |
| A0A1R1APE4 | Paenibacillus sp. FSL A5-0031 |
| A0A1R1D0W9 | Paenibacillus sp. FSL H8-0548 |
| A0A1R1EPP2 | Paenibacillus rhizosphaerae |
| A0A1R1ETF9 | Paenibacillus sp. FSL R5-0490 |
| A0A1R1GPK8 | Paenibacillus sp. FSL R7-0337 |
| A0A1S1FML9 | Bacillus sp. HMSC76G11 |
| A0A1S1YGZ1 | Cytobacillus oceanisediminis |
| A0A1S2F9E4 | Paenibacillus sp. LC231 |
| A0A1S2LJR3 | Anaerobacillus arseniciselenatis |
| A0A1S2LMJ2 | Anaerobacillus alkalilacustris |
| A0A1S2M2M6 | Anaerobacillus alkalidiazotrophicus |
| A0A1S2R4N1 | Bacillus sp. MUM 13 |
| A0A1S2REB7 | Bacillus sp. MUM 116 |
| A0A1S6IZB5 | Desulfotomaculum ferrireducens |
| A0A1S8G6G6 | Bacillus mycoides |
| A0A1S9CD67 | Epulopiscium sp. AS2M-Bin001 |
| A0A1S9CJQ8 | Epulopiscium sp. Nele67-Bin005 |
| A0A1S9I655 | Clostridium tepidum |
| A0A1S9TRK3 | Bacillus cereus |
| A0A1S9V6Y2 | Bacillus cereus |
| A0A1S9XRV9 | Bacillus mycoides |
| A0A1T2P9U8 | Bacillus cereus |
| A0A1T2PLB9 | Bacillus cereus |
| A0A1T2T2H4 | Bacillus cereus |
| A0A1T2X1Y8 | Paenibacillus selenitireducens |
| A0A1T3V387 | Bacillus anthracis |
| A0A1T4KQ78 | Garciella nitratireducens DSM 15102 |
| A0A1T4M1C6 | Carboxydocella sporoproducens DSM 16521 |
| A0A1T4N5I1 | Selenihalanaerobacter shriftii |
| A0A1T4W911 | Clostridium sp. USBA 49 |
| A0A1T4XEZ7 | Caloramator quimbayensis |
| A0A1T4Y6S3 | Sporosarcina newyorkensis |
| A0A1T4ZVG8 | Lysinibacillus sp. AC-3 |
| A0A1T5KRS1 | Maledivibacter halophilus |
| A0A1T5LV26 | Maledivibacter halophilus |
| A0A1U6JH99 | Bacillus sp. V-88 |
| A0A1U7M820 | Tissierella creatinophila DSM 6911 |
| A0A1U7MMM7 | Sporomusa sphaeroides DSM 2875 |
| A0A1U7PJX6 | Edaphobacillus lindanitolerans |
| A0A1U9K7A3 | Novibacillus thermophilus |
| A0A1V1HY46 | Romboutsia ilealis |
| A0A1V2A6A5 | Domibacillus epiphyticus |
| A0A1V2HDL6 | Bacillus cereus |
| A0A1V2M959 | Epulopiscium sp. SCG-C07WGA-EpuloA2 |
| A0A1V2SKR6 | Bacillus sp. VT-16-64 |
| A0A1V2YCU8 | Epulopiscium sp. Nuni2H_MBin003 |
| A0A1V2YJR5 | Epulopiscium sp. Nuni2H_MBin003 |
| A0A1V2YJY4 | Epulopiscium sp. Nuni2H_MBin001 |
| A0A1V3G4H6 | Fictibacillus arsenicus |
| A0A1V4I8L2 | [Clostridium] thermoalcaliphilum |
| A0A1V4SRX6 | Ruminiclostridium hungatei |
| A0A1V4SU42 | Clostridium thermobutyricum DSM 4928 |
| A0A1V4VLK6 | Pelotomaculum sp. PtaB.Bin117 |
| A0A1V4VUH2 | Pelotomaculum sp. PtaB.Bin104 |
| A0A1V5BQQ0 | Pelotomaculum sp. PtaU1.Bin065 |
| A0A1V5BTT9 | Pelotomaculum sp. PtaU1.Bin065 |
| A0A1V5KFX2 | Firmicutes bacterium ADurb.Bin467 |
| A0A1V5L119 | Firmicutes bacterium ADurb.Bin456 |
| A0A1V5MLC7 | Firmicutes bacterium ADurb.Bin419 |
| A0A1V5S7A8 | Firmicutes bacterium ADurb.Bin300 |
| A0A1V5TZH5 | Firmicutes bacterium ADurb.Bin248 |
| A0A1V5XFX1 | Firmicutes bacterium ADurb.Bin193 |
| A0A1V5YBX9 | Firmicutes bacterium ADurb.Bin182 |
| A0A1V9IQ78 | Clostridium sporogenes |
| A0A1V9W066 | Bacillus sp. CDB3 |
| A0A1W1UHE8 | Desulfonispora thiosulfatigenes DSM 11270 |
| A0A1W1UHI0 | Desulfonispora thiosulfatigenes DSM 11270 |
| A0A1W1W2L7 | Thermanaeromonas toyohensis ToBE |
| A0A1W1W793 | Sulfobacillus thermosulfidooxidans (strain DSM 9293 / VKM B-1269 / AT-1) |
| A0A1W1Z1E7 | Papillibacter cinnamivorans DSM 12816 |
| A0A1W2BSZ8 | Sporomusa malonica |
| A0A1W2GW40 | Bacillus sp. JKS001846 |
| A0A1W6A2X6 | Bacillus mycoides |
| A0A1X3MIN9 | Bacillus toyonensis |
| A0A1X7G627 | Paenibacillus uliginis N3/975 |
| A0A1X9MAY4 | Alkalihalobacillus krulwichiae |
| A0A1Y0CSK5 | Bacillus horikoshii |
| A0A1Y0TWD8 | Bacillus thuringiensis |
| A0A1Y2T509 | Symbiobacterium thermophilum |
| A0A1Y3ML41 | Bacillus pseudomycoides |
| A0A1Y3RJ62 | Flavonifractor sp. An91 |
| A0A1Y3RN37 | Flavonifractor sp. An9 |
| A0A1Y3RU35 | Gemmiger sp. An87 |
| A0A1Y3SNW4 | Flavonifractor sp. An82 |
| A0A1Y3SVF6 | Pseudoflavonifractor sp. An85 |
| A0A1Y3TAG8 | Faecalibacterium sp. An77 |
| A0A1Y3WVM8 | Faecalibacterium sp. An58 |
| A0A1Y3XHS1 | Flavonifractor sp. An52 |
| A0A1Y3XKU1 | Gemmiger sp. An50 |
| A0A1Y3YMJ5 | Pseudoflavonifractor sp. An44 |
| A0A1Y3ZTH2 | Flavonifractor sp. An4 |
| A0A1Y4C6G8 | Flavonifractor sp. An306 |
| A0A1Y4EVI7 | Anaeromassilibacillus sp. An250 |
| A0A1Y4FPP0 | Flavonifractor plautii |
| A0A1Y4HEB1 | Anaerofilum sp. An201 |
| A0A1Y4HZM4 | Anaeromassilibacillus sp. An200 |
| A0A1Y4ISQ3 | Gemmiger sp. An194 |
| A0A1Y4J6T6 | Faecalibacterium sp. An192 |
| A0A1Y4KBK2 | Pseudoflavonifractor sp. An187 |
| A0A1Y4LQJ8 | Pseudoflavonifractor sp. An184 |
| A0A1Y4M3H2 | Pseudoflavonifractor sp. An176 |
| A0A1Y4RWX2 | Flavonifractor sp. An135 |
| A0A1Y4T4A1 | Faecalibacterium sp. An121 |
| A0A1Y4TCI6 | Faecalibacterium sp. An122 |
| A0A1Y4TNC5 | Gemmiger sp. An120 |
| A0A1Y4UF12 | Flavonifractor sp. An112 |
| A0A1Y4WLT8 | Flavonifractor sp. An10 |
| A0A1Y4WM03 | Flavonifractor sp. An100 |
| A0A1Y4X3Z3 | Brevibacillus brevis |
| A0A1Y5K2Z6 | Paenibacillus sp. MY03 |
| A0A1Y5Z8A0 | Bacillus mobilis |
| A0A1Y5ZW25 | Bacillus cereus |
| A0A1Y6E7Z6 | Bacillus sp. OV166 |
| A0A1Z5HR28 | Calderihabitans maritimus |
| A0A1Z5HRM8 | Calderihabitans maritimus |
| A0A212J6A5 | uncultured Eubacteriales bacterium |
| A0A212LQJ6 | uncultured Sporomusa sp |
| A0A220MBR1 | Brevibacillus formosus |
| A0A220U462 | Virgibacillus phasianinus |
| A0A221BRN2 | Bacillus cereus |
| A0A221MAN7 | Virgibacillus necropolis |
| A0A223EHG3 | Peribacillus simplex NBRC 15720 = DSM 1321 |
| A0A223KWL1 | Bacillus cohnii |
| A0A226BW94 | Natranaerobius trueperi |
| A0A226QXI9 | Bacillus sp. M13(2017) |
| A0A229MA73 | Bacillus sp. KbaL1 |
| A0A231I5E7 | Bacillus thuringiensis |
| A0A231REP0 | Cohnella sp. CIP 111063 |
| A0A231W5H4 | Bacillus sp. OG2 |
| A0A235FBB7 | Fictibacillus aquaticus |
| A0A239HN74 | Bacillus sp. OK838 |
| A0A239JX23 | Anaerovirgula multivorans |
| A0A242WES8 | Bacillus thuringiensis serovar mexicanensis |
| A0A242WFM2 | Bacillus thuringiensis serovar cameroun |
| A0A242XPL1 | Bacillus thuringiensis serovar guiyangiensis |
| A0A242Y852 | Bacillus thuringiensis serovar novosibirsk |
| A0A242Z661 | Bacillus wiedmannii |
| A0A242ZR82 | Bacillus thuringiensis serovar kim |
| A0A243AJR2 | Bacillus thuringiensis serovar navarrensis |
| A0A243AYX9 | Bacillus thuringiensis serovar poloniensis |
| A0A243BHM2 | Bacillus thuringiensis serovar pingluonsis |
| A0A243C540 | Bacillus thuringiensis serovar yosoo |
| A0A243D230 | Bacillus thuringiensis serovar vazensis |
| A0A243D8H4 | Bacillus thuringiensis serovar subtoxicus |
| A0A243DZM3 | Bacillus thuringiensis subsp. darmstadiensis |
| A0A243E7Y0 | Bacillus thuringiensis serovar toumanoffi |
| A0A243F0A6 | Bacillus thuringiensis subsp. kumamotoensis |
| A0A243GLN5 | Bacillus thuringiensis subsp. finitimus |
| A0A243IW53 | Bacillus thuringiensis subsp. konkukian |
| A0A243JCV4 | Bacillus thuringiensis serovar pirenaica |
| A0A243KSL6 | Bacillus thuringiensis subsp. higo |
| A0A243KST8 | Bacillus thuringiensis serovar argentinensis |
| A0A243L1P4 | Bacillus thuringiensis serovar iberica |
| A0A243LH44 | Bacillus thuringiensis subsp. jegathesan |
| A0A243MNK8 | Bacillus thuringiensis serovar zhaodongensis |
| A0A243NFL2 | Bacillus thuringiensis subsp. medellin |
| A0A246PPY5 | Bacillus sp. K2I17 |
| A0A249X6P6 | Bacillus cereus |
| A0A259TBR9 | Paenibacillus sp. XY044 |
| A0A259UC19 | Sporomusa acidovorans DSM 3132 |
| A0A259UJE8 | Sporomusa silvacetica DSM 10669 |
| A0A261QKA1 | Bacillaceae bacterium SAS-127 |
| A0A263BSZ4 | Lottiidibacillus patelloidae |
| A0A263BVX5 | Lottiidibacillus patelloidae |
| A0A264DDD2 | Paenibacillus sp. VTT E-133280 |
| A0A264EBE1 | Paenibacillus odorifer |
| A0A265NAG4 | Virgibacillus indicus |
| A0A265QAQ8 | Tissierella sp. P1 |
| A0A267MHD8 | Anaeromicrobium sediminis |
| A0A267MHP5 | Anaeromicrobium sediminis |
| A0A267MJL2 | Anaeromicrobium sediminis |
| A0A267TPE6 | Caldibacillus hisashii |
| A0A268E5V2 | Bacillus sp. 7586-K |
| A0A268IBL0 | Bacillus sp. 7504-2 |
| A0A268IVB9 | Bacillus sp. 7894-2 |
| A0A268KAD0 | Bacillus sp. 7884-1 |
| A0A270AKB4 | Peribacillus simplex |
| A0A271M5K9 | Bacillaceae bacterium SAOS 7 |
| A0A285CIQ7 | Bacillus oleivorans |
| A0A285GKD8 | Orenia metallireducens |
| A0A285STC0 | Ureibacillus xyleni |
| A0A285UCY4 | Ureibacillus acetophenoni |
| A0A291BG03 | Brevibacillus brevis X23 |
| A0A291JGS9 | Staphylococcus nepalensis |
| A0A291TDQ9 | Faecalibacterium prausnitzii |
| A0A2A2IH90 | Virgibacillus profundi |
| A0A2A2PDV7 | Bacillus toyonensis |
| A0A2A4BH91 | Peribacillus simplex |
| A0A2A5LA92 | Paenibacillus lautus |
| A0A2A6Z975 | Faecalibacterium prausnitzii |
| A0A2A6ZHL0 | Faecalibacterium prausnitzii |
| A0A2A6ZWI1 | Faecalibacterium prausnitzii |
| A0A2A7AE65 | Faecalibacterium prausnitzii |
| A0A2A7AQI1 | Faecalibacterium prausnitzii |
| A0A2A7AWL6 | Faecalibacterium prausnitzii |
| A0A2A7B591 | Faecalibacterium prausnitzii |
| A0A2A7BEP3 | Faecalibacterium prausnitzii |
| A0A2A7BV03 | Bacillus wiedmannii |
| A0A2A7DEM5 | Bacillus anthracis |
| A0A2A7EDJ2 | Bacillus sp. AFS094611 |
| A0A2A7HAI2 | Bacillus sp. AFS098217 |
| A0A2A7HTZ5 | Bacillus cereus |
| A0A2A7IH20 | Bacillus sp. AFS096315 |
| A0A2A7II50 | Bacillus cereus |
| A0A2A7WZZ4 | Bacillus sp. AFS002410 |
| A0A2A7X994 | Bacillus wiedmannii |
| A0A2A7ZLW7 | Bacillus cereus |
| A0A2A8AWH7 | Bacillus wiedmannii |
| A0A2A8E333 | Bacillus cereus |
| A0A2A8FQB8 | Bacillus sp. AFS026049 |
| A0A2A8G170 | Bacillus wiedmannii |
| A0A2A8HLR4 | Bacillus toyonensis |
| A0A2A8ILY2 | Bacillus sp. AFS006103 |
| A0A2A8ISM0 | Bacillus cereus |
| A0A2A8MS79 | Bacillus sp. AFS001701 |
| A0A2A8NJR9 | Bacillus thuringiensis |
| A0A2A8PSJ6 | Bacillus cereus |
| A0A2A8S2B7 | Bacillus cereus |
| A0A2A8SBK7 | Bacillus sp. AFS018417 |
| A0A2A8TRY3 | Bacillus sp. AFS017274 |
| A0A2A8UBR5 | Bacillus cereus |
| A0A2A8UTJ5 | Bacillus sp. AFS015896 |
| A0A2A8UZM5 | Bacillus sp. AFS015802 |
| A0A2A9A748 | Bacillus cereus |
| A0A2A9BY70 | Bacillus sp. es.034 |
| A0A2A9RE12 | Bacillus sp. AFS088145 |
| A0A2A9XA34 | Bacillus cereus |
| A0A2B0C3G2 | Bacillus cereus |
| A0A2B0L4X9 | Bacillus cereus |
| A0A2B0M9N4 | Bacillus cereus |
| A0A2B0WYF9 | Bacillus cereus |
| A0A2B0Y802 | Bacillus anthracis |
| A0A2B1DDL6 | Bacillus cereus |
| A0A2B1DVW1 | Bacillus cereus |
| A0A2B1KQ32 | Bacillus cereus |
| A0A2B1RFM2 | Bacillus cereus |
| A0A2B1VEH4 | Bacillus cereus |
| A0A2B2C870 | Bacillus sp. AFS073361 |
| A0A2B2GL24 | Bacillus cereus |
| A0A2B2LTU8 | Bacillus cereus |
| A0A2B3EJV2 | Bacillus thuringiensis |
| A0A2B3LAL9 | Bacillus thuringiensis |
| A0A2B3UBJ6 | Bacillus cereus |
| A0A2B4FGT8 | Bacillus sp. AFS059628 |
| A0A2B4LEL1 | Bacillus cereus |
| A0A2B4P2J7 | Bacillus sp. AFS075960 |
| A0A2B4XUJ2 | Bacillus mycoides |
| A0A2B5IUJ3 | Bacillus wiedmannii |
| A0A2B5J402 | Bacillus wiedmannii |
| A0A2B5KD50 | Bacillus wiedmannii |
| A0A2B5XRJ4 | Bacillus wiedmannii |
| A0A2B6CB44 | Bacillus anthracis |
| A0A2B6NIU4 | Bacillus toyonensis |
| A0A2B6QHE4 | Bacillus toyonensis |
| A0A2B6R361 | Bacillus pseudomycoides |
| A0A2B8G019 | Bacillus cereus |
| A0A2B8ITP8 | Bacillus anthracis |
| A0A2B8LWR8 | Bacillus cereus |
| A0A2B8U1N7 | Bacillus sp. AFS055030 |
| A0A2B9AHC7 | Bacillus sp. AFS053548 |
| A0A2B9BAK0 | Bacillus cereus |
| A0A2B9DKF8 | Bacillus cereus |
| A0A2B9ELR3 | Bacillus cereus |
| A0A2B9MXX8 | Bacillus cereus |
| A0A2B9PD52 | Bacillus thuringiensis |
| A0A2B9PKS3 | Bacillus cereus |
| A0A2B9UAJ3 | Bacillus cereus |
| A0A2B9XDW2 | Bacillus thuringiensis |
| A0A2C0CGH1 | Bacillus cereus |
| A0A2C0ZJI7 | Bacillus sp. AFS041924 |
| A0A2C1AHU8 | Bacillus cereus |
| A0A2C1D606 | Bacillus cereus |
| A0A2C1L0U2 | Bacillus sp. AFS040349 |
| A0A2C1YG79 | Bacillus cereus |
| A0A2C1YWT2 | Bacillus sp. AFS037270 |
| A0A2C2ACV3 | Bacillus cereus |
| A0A2C2C3G3 | Bacillus cereus |
| A0A2C2UWY0 | Bacillus cereus |
| A0A2C2VD57 | Bacillus sp. AFS031507 |
| A0A2C3G960 | Bacillus anthracis |
| A0A2C4Q0S2 | Bacillus wiedmannii |
| A0A2C4QUS1 | Bacillus toyonensis |
| A0A2C9YWB3 | Bacillus thuringiensis subsp. kyushuensis |
| A0A2D1SZN4 | Solibacillus sp. R5-41 |
| A0A2G2M9A1 | Alkaliphilus sp |
| A0A2G3PYK1 | Lachnospiraceae bacterium |
| A0A2G5W133 | Sporosarcina sp. P10 |
| A0A2G5WL10 | Sporosarcina sp. P13 |
| A0A2G5WS38 | Sporosarcina sp. P16a |
| A0A2G5X4D5 | Sporosarcina sp. P16b |
| A0A2G5XCZ7 | Sporosarcina sp. P17b |
| A0A2G5XJ69 | Sporosarcina sp. P19 |
| A0A2G5ZBA1 | Sporosarcina sp. P29 |
| A0A2G6AQU1 | Sporosarcina sp. P34 |
| A0A2G6B9H5 | Sporosarcina sp. P3 |
| A0A2G6QCC1 | Bacillus fungorum |
| A0A2G7HGT3 | Clostridium combesii |
| A0A2H3M8S4 | Bacillus pseudomycoides |
| A0A2H3QV93 | Bacillus sp. AFS012607 |
| A0A2I0R7A2 | Bacillus cereus Rock4-18 |
| A0A2I0V659 | Lysinibacillus fusiformis |
| A0A2I4NHD5 | Clostridium botulinum |
| A0A2J4JRY0 | Faecalibacterium prausnitzii |
| A0A2J9DL65 | Bacillus thuringiensis |
| A0A2K2FMQ3 | Pseudoclostridium thermosuccinogenes |
| A0A2K8TKH8 | Bacillus cereus |
| A0A2K8ZED8 | Bacillus sp. HBCD-sjtu |
| A0A2K9E3N6 | Acetivibrio saccincola |
| A0A2K9MUU7 | Clostridium sporogenes |
| A0A2K9P4M5 | Monoglobus pectinilyticus |
| A0A2L1GLY3 | Desulfobulbus oralis |
| A0A2L2XDB8 | Desulfocucumis palustris |
| A0A2M9ML56 | Paenibacillus sp. GM2FR |
| A0A2M9P0K1 | Bacillus sp. mrc49 |
| A0A2M9P1F7 | Bacillus sp. mrc49 |
| A0A2M9P1G9 | Bacillus sp. mrc49 |
| A0A2M9Q2R8 | Lysinibacillus xylanilyticus |
| A0A2M9X824 | Bacillus cereus |
| A0A2N0F8U6 | Viridibacillus sp. OK051 |
| A0A2N0Y8R8 | Bacillus sp. BA3 |
| A0A2N1JVA8 | Bacillus sp. SN10 |
| A0A2N2AZS9 | Firmicutes bacterium HGW-Firmicutes-7 |
| A0A2N2BZ01 | Firmicutes bacterium HGW-Firmicutes-21 |
| A0A2N2CV14 | Firmicutes bacterium HGW-Firmicutes-16 |
| A0A2N2D3E3 | Firmicutes bacterium HGW-Firmicutes-15 |
| A0A2N2D566 | Firmicutes bacterium HGW-Firmicutes-15 |
| A0A2N2DCM7 | Firmicutes bacterium HGW-Firmicutes-14 |
| A0A2N2DLY1 | Firmicutes bacterium HGW-Firmicutes-13 |
| A0A2N2E3B8 | Firmicutes bacterium HGW-Firmicutes-12 |
| A0A2N2EG20 | Firmicutes bacterium HGW-Firmicutes-1 |
| A0A2N3LJI1 | Bacillus camelliae |
| A0A2N3NQM5 | Bacillus sp. BI3 |
| A0A2N5FJF8 | Bacillus sp. UMB0893 |
| A0A2N5FY00 | Bacillus sp. UMB0728 |
| A0A2N5GE75 | Bacillus sp. V3-13 |
| A0A2N5GH83 | Bacillus canaveralius |
| A0A2N5GZV2 | Bacillus sp. T33-2 |
| A0A2N5HEA0 | Neobacillus cucumis |
| A0A2N5I912 | Bacillus sp. M6-12 |
| A0A2N5M5S1 | Peribacillus deserti |
| A0A2N5MNK4 | Bacillus sp. V5-8f |
| A0A2N6RLW4 | Bacillus sp. UMB0899 |
| A0A2P1THR6 | Clostridium botulinum |
| A0A2P1WPY9 | Oceanobacillus iheyensis |
| A0A2P2BMP6 | Romboutsia hominis |
| A0A2P7UJU8 | Brevibacillus fortis |
| A0A2R4N1Y7 | Carboxydocella thermautotrophica |
| A0A2R5EV86 | Paenibacillus agaridevorans |
| A0A2S0JGS4 | Lysinibacillus sp. B2A1 |
| A0A2S0K5C2 | Lysinibacillus sphaericus |
| A0A2S1A6H4 | Bacillus cytotoxicus |
| A0A2S3QGA7 | Sulfobacillus sp. hq2 |
| A0A2S5D5A1 | Lysinibacillus sphaericus |
| A0A2S5G9Y4 | Jeotgalibacillus proteolyticus |
| A0A2S5I3X3 | Brevibacillus laterosporus |
| A0A2S6G021 | Clostridium algidicarnis DSM 15099 |
| A0A2S8RDK9 | Acetivibrio saccincola |
| A0A2S8ULQ8 | Bacillus sp. MYb209 |
| A0A2S8VTN1 | Bacillus sp. MYb78 |
| A0A2S9H9I8 | Bacillus cereus |
| A0A2S9HTJ4 | Bacillus sp. MYb56 |
| A0A2S9Y3T3 | Bacillus sp. M21 |
| A0A2T0ATZ0 | Clostridium thermopalmarium DSM 5974 |
| A0A2T0AVH7 | Moorella humiferrea |
| A0A2T0B2S3 | Clostridium liquoris |
| A0A2T0BND5 | Clostridium luticellarii |
| A0A2T0DZ65 | Bacillus toyonensis |
| A0A2T0EY66 | Bacillus thuringiensis |
| A0A2T2WSB6 | Sulfobacillus thermosulfidooxidans |
| A0A2T2X9B7 | Sulfobacillus benefaciens |
| A0A2T2XCW4 | Sulfobacillus benefaciens |
| A0A2T4SAZ4 | Staphylococcus nepalensis |
| A0A2T5UND8 | Bacillus sp. OV752 |
| A0A2T6EGR9 | Bacillus sporothermodurans |
| A0A2T6JSF6 | Paenisporosarcina sp. OV554 |
| A0A2T7YM27 | Bacillus thuringiensis |
| A0A2U1CD65 | Intestinimonas butyriciproducens |
| A0A2U1K0A2 | Pueribacillus theae |
| A0A2U3L9L7 | Candidatus Desulfosporosinus infrequens |
| A0A2U8DXJ4 | Clostridium drakei |
| A0A2U8ECB0 | Caldibacillus thermoamylovorans |
| A0A2V2CFN9 | Clostridia bacterium |
| A0A2V2D263 | Clostridiales bacterium |
| A0A2V2DZA7 | Clostridiales bacterium |
| A0A2V2E2X6 | Clostridiales bacterium |
| A0A2V2EVA5 | Clostridiales bacterium |
| A0A2V2FAV6 | Oscillospiraceae bacterium |
| A0A2V2FZ12 | Clostridiales Family XIII bacterium |
| A0A2V2G642 | Clostridiales bacterium |
| A0A2V2GP49 | Oscillospiraceae bacterium |
| A0A2V3A2Z5 | Cytobacillus oceanisediminis |
| A0A2V3W5G7 | Pseudogracilibacillus auburnensis |
| A0A2W0H6T2 | Bacillus lacisalsi |
| A0A2W1NRL8 | Paenibacillus xerothermodurans |
| A0A2W4K387 | Firmicutes bacterium |
| A0A2W4KCH9 | Firmicutes bacterium |
| A0A2W4KN14 | Firmicutes bacterium |
| A0A2W4MLX1 | Caldicoprobacter oshimai |
| A0A2W4NA88 | Firmicutes bacterium |
| A0A2W4QJI6 | Firmicutes bacterium |
| A0A2W6N1B4 | Clostridium perfringens |
| A0A2W7N3B1 | Psychrobacillus insolitus |
| A0A2X0YUK1 | Lysinibacillus capsici |
| A0A2X2WD83 | Clostridium cochlearium |
| A0A2X2WFW3 | Clostridium perfringens |
| A0A2X2Y040 | Clostridium perfringens |
| A0A2X4ZEY1 | Lederbergia lentus |
| A0A2Z4MEI0 | Brevibacillus brevis |
| A0A2Z4W7Y4 | Clostridiaceae bacterium 14S0207 |
| A0A316LC94 | Clostridiales bacterium |
| A0A316N0M8 | Oscillospiraceae bacterium |
| A0A316PHE9 | Oscillospiraceae bacterium |
| A0A316Q1U9 | Clostridiales bacterium |
| A0A316Q8Q5 | Clostridiales bacterium |
| A0A316QIS8 | Clostridiales bacterium |
| A0A316QL15 | Clostridiales bacterium |
| A0A316RHL2 | Oscillospiraceae bacterium |
| A0A316RWG6 | Oscillospiraceae bacterium |
| A0A316T704 | Massilioclostridium sp |
| A0A317KT67 | Gracilibacillus dipsosauri |
| A0A317TQ04 | Clostridium perfringens |
| A0A318THP6 | Ureibacillus chungkukjangi |
| A0A323TDH3 | Salipaludibacillus keqinensis |
| A0A327S649 | Bacillus sp. YR335 |
| A0A328KTD5 | Bacillus sp. SRB_8 |
| A0A328LFT4 | Bacillus sp. SRB_331 |
| A0A328TZK2 | Paenibacillus montanisoli |
| A0A328UM82 | Hydrogeniiclostidium mannosilyticum |
| A0A328WBS4 | Paenibacillus lautus |
| A0A329L399 | Paenibacillus sp. YN15 |
| A0A329MK52 | Paenibacillus contaminans |
| A0A329TG41 | Faecalibacterium prausnitzii |
| A0A329TU74 | Faecalibacterium prausnitzii |
| A0A329TYW0 | Faecalibacterium prausnitzii |
| A0A329U8K4 | Faecalibacterium prausnitzii |
| A0A329U9F6 | Faecalibacterium prausnitzii |
| A0A329UL90 | Faecalibacterium prausnitzii |
| A0A329UXQ3 | Faecalibacterium prausnitzii |
| A0A336QS83 | Clostridium perfringens |
| A0A343J9Y3 | Clostridium isatidis |
| A0A345BZ74 | Salicibibacter kimchii |
| A0A345P196 | Sporosarcina sp. PTS2304 |
| A0A345PJ59 | Oceanobacillus zhaokaii |
| A0A345X2R3 | Bacillus sp. COPE52 |
| A0A347V4E7 | Bacillus thuringiensis LM1212 |
| A0A348AFU3 | Methylomusa anaerophila |
| A0A348P5U0 | Oscillospiraceae bacterium |
| A0A348Z7V0 | Clostridium sp. |
| A0A349DBV8 | Oscillospiraceae bacterium |
| A0A349HQI6 | Clostridiales bacterium |
| A0A349PV89 | Oscillibacter sp |
| A0A349Q832 | Clostridiales bacterium |
| A0A349U4N8 | Desulfotomaculum sp |
| A0A349YM96 | Lachnospiraceae bacterium |
| A0A350BFC8 | Firmicutes bacterium |
| A0A350NP44 | Firmicutes bacterium |
| A0A350WZJ4 | Firmicutes bacterium |
| A0A351ED52 | Oscillospiraceae bacterium |
| A0A351F9J4 | Oscillospiraceae bacterium |
| A0A351J1F0 | Clostridiales bacterium |
| A0A351J6K4 | Firmicutes bacterium |
| A0A351KH25 | Clostridiaceae bacterium |
| A0A351QW08 | Clostridium sp. |
| A0A352CXV3 | Ruminococcus sp |
| A0A352IKU0 | Clostridiales bacterium |
| A0A352NFS8 | Pelotomaculum sp |
| A0A352RLV5 | Oscillibacter sp |
| A0A352SVM3 | Clostridiales bacterium |
| A0A352UNA3 | Clostridiales bacterium |
| A0A353EWN2 | Clostridiales bacterium |
| A0A353HAX4 | Clostridiales bacterium |
| A0A353K333 | Clostridiaceae bacterium |
| A0A353M1U8 | Firmicutes bacterium |
| A0A353MQ25 | Firmicutes bacterium |
| A0A353QJM9 | Firmicutes bacterium |
| A0A353T297 | Clostridiales bacterium |
| A0A354FE88 | Peptococcaceae bacterium |
| A0A354FFD0 | Peptococcaceae bacterium |
| A0A354HTE7 | Firmicutes bacterium |
| A0A354KV82 | Terrisporobacter glycolicus |
| A0A354MXH6 | Clostridiales bacterium |
| A0A354YWL8 | Syntrophomonas wolfei |
| A0A354YZH7 | Syntrophomonas wolfei |
| A0A355BIC4 | Firmicutes bacterium |
| A0A355D522 | Clostridium sp. |
| A0A355FSA5 | Firmicutes bacterium |
| A0A355GQV5 | Firmicutes bacterium |
| A0A355KU56 | Oscillospiraceae bacterium |
| A0A355S2K7 | Clostridiaceae bacterium |
| A0A355SCQ2 | Clostridiaceae bacterium |
| A0A356B4R0 | Clostridiales bacterium |
| A0A356BUG1 | Firmicutes bacterium |
| A0A356CZ28 | Ruminococcus sp |
| A0A356GG05 | Clostridiales bacterium |
| A0A356P5S4 | Desulfosporosinus sp |
| A0A356PU05 | Oscillospiraceae bacterium |
| A0A356U822 | Syntrophomonas sp |
| A0A356UC60 | Syntrophomonas sp |
| A0A356UH50 | Desulfotomaculum sp |
| A0A356XG36 | Clostridiales bacterium |
| A0A356Z544 | Syntrophomonas sp |
| A0A356ZD95 | Syntrophomonas sp |
| A0A357AMG0 | Ruminiclostridium sp |
| A0A357AXI3 | Clostridiales bacterium |
| A0A357CYZ1 | Clostridiales bacterium |
| A0A357D4I5 | Firmicutes bacterium |
| A0A357M3W6 | Paenibacillus sp. |
| A0A357R5N9 | Firmicutes bacterium |
| A0A357T4X7 | Firmicutes bacterium |
| A0A357TE93 | Peptococcaceae bacterium |
| A0A357WTF0 | Oscillospiraceae bacterium |
| A0A357WU76 | Oscillospiraceae bacterium |
| A0A358LJQ9 | Oscillospiraceae bacterium |
| A0A358M392 | Clostridiales bacterium |
| A0A358PZD7 | Desulfosporosinus sp |
| A0A358Q0A3 | Desulfosporosinus sp |
| A0A358QZN1 | Desulfotomaculum sp |
| A0A358RK06 | Clostridiales bacterium |
| A0A358TZ47 | Desulfosporosinus sp |
| A0A358U2Q8 | Desulfosporosinus sp |
| A0A359B8Z7 | Desulfotomaculum sp |
| A0A359CPP1 | Clostridiaceae bacterium |
| A0A366F230 | Bacillus aquimaris |
| A0A366G4Y6 | Bacillus sp. DB-2 |
| A0A366ICG1 | Alkalibaculum bacchi |
| A0A366JYS9 | Cytobacillus firmus |
| A0A366XV32 | Bacillus taeanensis |
| A0A368WKU2 | Bacillus sp. NFR08 |
| A0A369BER6 | Fontibacillus phaseoli |
| A0A369CT49 | Bacillus sp. AG102 |
| A0A370GHS8 | Falsibacillus pallidus |
| A0A371IVE6 | Romboutsia maritimum |
| A0A371J1F8 | Romboutsia weinsteinii |
| A0A371P7N4 | Paenibacillus paeoniae |
| A0A371SGS9 | Bacillus sp. HNG |
| A0A372LAQ3 | Bacillus glennii |
| A0A372LNH2 | Bacillus saganii |
| A0A372V9C9 | Subdoligranulum sp. AM16-9 |
| A0A372VBR2 | Subdoligranulum sp. AM16-9 |
| A0A372X1Q7 | Subdoligranulum sp. AM23-21AC |
| A0A372X241 | Subdoligranulum sp. AM23-21AC |
| A0A373LBB0 | Ruminococcus sp. AF37-20 |
| A0A373MU10 | Ruminococcus sp. AF34-12 |
| A0A373NBN2 | Faecalibacterium sp. OF04-11AC |
| A0A373Q021 | Ruminococcus sp. AM54-1NS |
| A0A373SIT7 | Ruminococcus sp. AF25-19 |
| A0A373UCK2 | Ruminococcus sp. AF21-11 |
| A0A373V6B2 | Ruminococcus sp. AF19-15 |
| A0A373VX01 | Ruminococcus sp. AF18-29 |
| A0A373WES9 | Ruminococcus sp. AF17-6 |
| A0A373XXP2 | Ruminococcus sp. AF16-50 |
| A0A374B7G4 | Ruminococcus sp. AM47-2BH |
| A0A374BTQ5 | Ruminococcus sp. AM43-6 |
| A0A374EVK4 | Ruminococcus sp. AM31-15AC |
| A0A374G9L4 | Ruminococcus sp. AM28-13 |
| A0A374HKU9 | Ruminococcus sp. TF12-2 |
| A0A380BE16 | Sporosarcina pasteurii |
| A0A380Y8S7 | Cytobacillus firmus |
| A0A381J0P5 | Clostridium perfringens |
| A0A381J564 | Clostridium putrefaciens |
| A0A385NVZ8 | Bacillus sp. Y1 |
| A0A385T4M6 | Brevibacillus laterosporus |
| A0A385TWI6 | Paenibacillus lautus |
| A0A385YSV1 | Paenisporosarcina sp. K2R23-3 |
| A0A386PGB6 | Clostridium septicum |
| A0A386XPA5 | Ethanoligenens harbinense |
| A0A386YMD8 | Clostridium novyi |
| A0A386ZX63 | Bacillus thuringiensis |
| A0A396LHN8 | Faecalibacterium sp. OF03-6AC |
| A0A396SLF4 | Lysinibacillus yapensis |
| A0A398BBJ0 | Mesobacillus zeae |
| A0A398BI76 | Peribacillus asahii |
| A0A3A0SB19 | Staphylococcus nepalensis |
| A0A3A1QVE5 | Bacillus salacetis |
| A0A3A1UV74 | Paenibacillus nanensis |
| A0A3A4U4F7 | Firmicutes bacterium |
| A0A3A5IIS4 | Bacillus sp. PK3_68 |
| A0A3A6CP95 | Faecalibacterium sp. AF27-11BH |
| A0A3A6EDQ9 | Faecalibacterium sp. AM43-5AT |
| A0A3A6EEW4 | Subdoligranulum sp. AF14-43 |
| A0A3A6EVM9 | Subdoligranulum sp. AF14-43 |
| A0A3A6JQ26 | Faecalibacterium sp. AF10-46 |
| A0A3A6JX76 | Subdoligranulum sp. OF01-18 |
| A0A3A6JXB8 | Subdoligranulum sp. OF01-18 |
| A0A3A6MN37 | Candidatus Desulforudis sp |
| A0A3A6N4I8 | Ammonifex sp |
| A0A3A6N9D5 | Dethiobacter sp |
| A0A3A6PL26 | Paenibacillus pinisoli |
| A0A3A8YVX8 | bacterium 1xD42-67 |
| A0A3A8YX15 | bacterium 1xD42-67 |
| A0A3A9FU49 | bacterium 1XD42-8 |
| A0A3A9J3Q2 | Anaerotruncus sp. 1XD22-93 |
| A0A3A9JB24 | Anaerotruncus sp. 1XD22-93 |
| A0A3A9JV21 | Thermoanaerobacteraceae bacterium SP2 |
| A0A3B0CBH9 | Paenibacillus ginsengarvi |
| A0A3B8HTT2 | Syntrophomonas sp |
| A0A3B8HZH0 | Syntrophomonas sp |
| A0A3B8I0G7 | Syntrophomonas sp |
| A0A3B8JBJ8 | Ruminiclostridium sp |
| A0A3B8K2L7 | Firmicutes bacterium |
| A0A3B8K5S8 | Firmicutes bacterium |
| A0A3B8NMQ4 | Peptococcaceae bacterium |
| A0A3B8NQ33 | Peptococcaceae bacterium |
| A0A3B8SB28 | Lachnospiraceae bacterium |
| A0A3B9M453 | Peptococcaceae bacterium |
| A0A3B9PWM8 | Clostridiales bacterium UBA9856 |
| A0A3B9QBZ8 | Clostridiales bacterium UBA9857 |
| A0A3B9SPM9 | Peptococcaceae bacterium |
| A0A3B9SQ47 | Peptococcaceae bacterium |
| A0A3B9SSG2 | Desulfotomaculum sp |
| A0A3B9SZX9 | Ruminococcus sp |
| A0A3B9UZB9 | Clostridium sp. |
| A0A3C0D3A0 | Faecalibacterium sp |
| A0A3C0H8A7 | Firmicutes bacterium |
| A0A3C0IKT5 | Ruminococcus sp |
| A0A3C0J364 | Firmicutes bacterium |
| A0A3C0NNV8 | Clostridiales bacterium |
| A0A3C0SRU3 | Clostridium sp. |
| A0A3C0WAL9 | Oscillospiraceae bacterium |
| A0A3C0X2F4 | Oscillospiraceae bacterium |
| A0A3C1I6C7 | Ornithinibacillus sp |
| A0A3C1K354 | Clostridiales bacterium |
| A0A3C1LXN5 | Oscillospiraceae bacterium |
| A0A3C1QLW9 | Firmicutes bacterium |
| A0A3C1QM67 | Firmicutes bacterium |
| A0A3C2DWK9 | Oscillospiraceae bacterium |
| A0A3C2DX09 | Oscillospiraceae bacterium |
| A0A3C2EN61 | Clostridiales bacterium |
| A0A3D0E9Y1 | Bacillus sp. |
| A0A3D0M0Q5 | Oscillospiraceae bacterium |
| A0A3D0M2D9 | Oscillospiraceae bacterium |
| A0A3D0XR26 | Oscillospiraceae bacterium |
| A0A3D0YB63 | Clostridiales bacterium |
| A0A3D0Z0V5 | Clostridiales bacterium |
| A0A3D0Z4R0 | Oscillospiraceae bacterium |
| A0A3D1FEY1 | Firmicutes bacterium |
| A0A3D1HXL8 | Oscillospiraceae bacterium |
| A0A3D1HYL3 | Oscillospiraceae bacterium |
| A0A3D1HZW6 | Oscillospiraceae bacterium |
| A0A3D1JTZ2 | Clostridiales bacterium |
| A0A3D1LB76 | Clostridiales bacterium |
| A0A3D1Q831 | Syntrophomonas sp |
| A0A3D1QFN2 | Syntrophomonas sp |
| A0A3D1SZ61 | Firmicutes bacterium |
| A0A3D1VMT5 | Clostridiales bacterium |
| A0A3D1X695 | Oscillibacter sp |
| A0A3D1XC91 | Clostridiales bacterium |
| A0A3D2A8W4 | Oscillospiraceae bacterium |
| A0A3D2CLU3 | Clostridiales bacterium |
| A0A3D2FZK3 | Clostridiales bacterium |
| A0A3D2N3Y6 | Ruminococcus sp |
| A0A3D2Q295 | Desulfotomaculum sp |
| A0A3D2QHE0 | Oscillospiraceae bacterium |
| A0A3D2XCI8 | Lachnoclostridium phytofermentans |
| A0A3D3AG35 | Clostridiaceae bacterium |
| A0A3D3Y2G2 | Oscillospiraceae bacterium |
| A0A3D3Y3K9 | Oscillospiraceae bacterium |
| A0A3D4D8Z5 | Oscillibacter sp |
| A0A3D4EET5 | Clostridiales bacterium |
| A0A3D4FIA8 | Clostridium sp. |
| A0A3D4JLU0 | Ruminococcus sp |
| A0A3D4LJ30 | Clostridiales bacterium |
| A0A3D4MVC5 | Clostridiales bacterium |
| A0A3D4T6A9 | Oscillospiraceae bacterium |
| A0A3D4W0A2 | Faecalibacterium sp |
| A0A3D5LNG6 | Oscillospiraceae bacterium |
| A0A3D5MNI4 | Clostridiales bacterium |
| A0A3D5MVP4 | Clostridium sp. |
| A0A3D5TL14 | Oscillospiraceae bacterium |
| A0A3D5U1F3 | Bacillus sp. |
| A0A3D5UD81 | Clostridiales bacterium |
| A0A3D5VJD4 | Firmicutes bacterium |
| A0A3D5WU24 | Clostridiales bacterium |
| A0A3D6BH55 | Clostridiales bacterium |
| A0A3D8GTM4 | Bacillus piezotolerans |
| A0A3D8PHT3 | Oceanobacillus chungangensis |
| A0A3D8PLJ7 | Oceanobacillus arenosus |
| A0A3D8YTM7 | Sporosarcina sp. BI001-red |
| A0A3D9GUW2 | Paenibacillus sp. VMFN-D1 |
| A0A3D9I4L6 | Cohnella phaseoli |
| A0A3D9UYD7 | Bacillus mycoides |
| A0A3E0KJF7 | Firmicutes bacterium |
| A0A3E0R8V1 | Brevibacillus sp |
| A0A3E2B443 | Evtepia gabavorous |
| A0A3E2JQL9 | Bacillus sp. V59.32b |
| A0A3E2T3K5 | Harryflintia acetispora |
| A0A3E2TCZ2 | Faecalibacterium prausnitzii |
| A0A3E2TTR2 | Faecalibacterium prausnitzii |
| A0A3E2U976 | Faecalibacterium prausnitzii |
| A0A3E2V306 | Faecalibacterium prausnitzii |
| A0A3E2XB04 | Faecalibacterium prausnitzii |
| A0A3F2ZV25 | Clostridium botulinum (strain 657 / Type Ba4) |
| A0A3F3JUB5 | Faecalibacterium prausnitzii |
| A0A3F3S3X6 | Tissierella praeacuta |
| A0A3G1KTZ2 | Candidatus Formimonas warabiya |
| A0A3G2R5F0 | Biomaibacter acetigenes |
| A0A3G3K1F8 | Cohnella candidum |
| A0A3G5UIN2 | Bacillus sp. FDAARGOS_527 |
| A0A3L7JTA0 | Falsibacillus albus |
| A0A3M7TUW7 | Bacillus sp. KQ-3 |
| A0A3M8ALJ3 | Brevibacillus agri |
| A0A3M8B1I5 | Brevibacillus gelatini |
| A0A3M8C1D3 | Brevibacillus invocatus |
| A0A3M8C567 | Brevibacillus panacihumi |
| A0A3M8D8Z4 | Brevibacillus fluminis |
| A0A3M8D921 | Brevibacillus nitrificans |
| A0A3M8H7F6 | Lysinibacillus halotolerans |
| A0A3N1XR06 | Mobilisporobacter senegalensis |
| A0A3N5AWX6 | Thermodesulfitimonas autotrophica |
| A0A3N5B993 | Aquisalibacillus elongatus |
| A0A3N5ZQX6 | Rummeliibacillus sp. TYF005 |
| A0A3N9UI22 | Lysinibacillus composti |
| A0A3P1B841 | Bacillus pacificus |
| A0A3P1ZI32 | Desulfovibrio sp. OH1186_COT-070 |
| A0A3P5WVG9 | Filibacter tadaridae |
| A0A3Q9B4M5 | Halocella sp. SP3-1 |
| A0A3Q9HNW8 | Anoxybacter fermentans |
| A0A3Q9HUB1 | Anoxybacter fermentans |
| A0A3Q9I8I0 | Paenibacillus lutimineralis |
| A0A3Q9QW48 | Neobacillus mesonae |
| A0A3Q9SE28 | [Brevibacterium] frigoritolerans |
| A0A3R6P0K6 | Faecalibacterium sp. AF28-13AC |
| A0A3R9D996 | Bacillus sp. |
| A0A3R9EA12 | Mesobacillus subterraneus |
| A0A3S0HP48 | Lysinibacillus telephonicus |
| A0A3S0WAW7 | Peribacillus cavernae |
| A0A3S0YIS1 | Clostridium perfringens |
| A0A3S1D9W1 | Paenibacillus zeisoli |
| A0A3S1DQS3 | Paenibacillus anaericanus |
| A0A3S1EPZ1 | Bacillus sp. VKPM B-3276 |
| A0A3S4S661 | Bacillus freudenreichii |
| A0A3S6Z3G0 | Sulfobacillus thermotolerans |
| A0A3S8RQT2 | Paenibacillus lentus |
| A0A3S9T4D7 | Bacillus thuringiensis |
| A0A3T0KYT2 | Peribacillus asahii |
| A0A3T1D5T5 | Cohnella abietis |
| A0A401UQV4 | Clostridium tagluense |
| A0A402GM91 | Bacillus cereus |
| A0A410MUG7 | Lysinibacillus sphaericus |
| A0A410PLB1 | Clostridium sp. JN-9 |
| A0A410PVX8 | Aminipila sp. JN-18 |
| A0A412AXQ3 | [Clostridium] leptum |
| A0A413D9G4 | Faecalibacterium prausnitzii |
| A0A413G7T5 | Anaerotruncus sp. AF02-27 |
| A0A415E3I5 | Emergencia timonensis |
| A0A416R2V9 | Pseudoflavonifractor sp. AF19-9AC |
| A0A416Z6U4 | Ruminococcaceae bacterium AF10-16 |
| A0A417HNF5 | Ruminococcaceae bacterium AM28-23LB |
| A0A417N7R6 | Ruminococcaceae bacterium TF06-43 |
| A0A417UNK9 | Faecalibacterium sp. OM04-11BH |
| A0A417YD98 | Oceanobacillus profundus |
| A0A417YT94 | Neobacillus notoginsengisoli |
| A0A418IN22 | Staphylococcus xylosus |
| A0A418SZG8 | Paenibacillus sp. 1011MAR3C5 |
| A0A419G0M4 | Peptococcaceae bacterium |
| A0A419SQN1 | Ammoniphilus oxalaticus |
| A0A419T4I6 | Thermohalobacter berrensis |
| A0A419T4J7 | Thermohalobacter berrensis |
| A0A420H172 | Bacillus toyonensis |
| A0A424YCT7 | Candidatus Syntrophonatronum acetioxidans |
| A0A426H481 | Peribacillus simplex |
| A0A428J5Z4 | Bacillus sp. HMF5848 |
| A0A428M8Y9 | Herbinix hemicellulosilytica |
| A0A428MVW9 | Bacillus salarius |
| A0A429X2H0 | Bacillus terrae |
| A0A429Y6F5 | Bacillus acidinfaciens |
| A0A432LCA4 | Lysinibacillus antri |
| A0A437SAY1 | Bacillus thuringiensis |
| A0A443T7P7 | Bacillus mycoides |
| A0A446IE11 | Paeniclostridium sordellii 8483 |
| A0A450CL81 | Clostridioides difficile |
| A0A480BIT0 | Paenibacillus naphthalenovorans |
| A0A494WRZ0 | Desulfofundulus salinum |
| A0A494XL46 | Cohnella endophytica |
| A0A494YYE2 | Lysinibacillus endophyticus |
| A0A494Z6S2 | Oceanobacillus bengalensis |
| A0A495A7J9 | Oceanobacillus halophilus |
| A0A498CPN2 | Anaerotruncus sp. 22A2-44 |
| A0A498DJM8 | Oceanobacillus piezotolerans |
| A0A4D7AN98 | Dysosmobacter welbionis |
| A0A4E7PMH5 | Bacillus thuringiensis subsp. israelensis |
| A0A4P6B805 | Moorella sp. E306M |
| A0A4P6BAA4 | Moorella sp. E306M |
| A0A4P6HJ06 | Desulfovibrio carbinolicus |
| A0A4P6UTD4 | Ureibacillus thermophilus |
| A0A4P7A078 | Paenisporosarcina antarctica |
| A0A4P7GT46 | Thermaerobacter sp. FW80 |
| A0A4P8SCE3 | Lysinibacillus sp. SGAir0095 |
| A0A4P9F4I9 | Bacillus paranthracis |
| A0A4Q0I7B0 | Acetivibrio mesophilus |
| A0A4Q0V0X7 | Clostridium tetani |
| A0A4Q0VDB4 | Clostridium tetani |
| A0A4Q0VNV3 | Anaerobacillus alkaliphilus |
| A0A4Q1SZX3 | Ammoniphilus sp. CFH 90114 |
| A0A4Q7QAB4 | Fictibacillus sp. BK138 |
| A0A4Q9DRH6 | Paenibacillus thalictri |
| A0A4Q9YBD1 | Bacillus mycoides |
| A0A4Q9Z630 | Bacillus mycoides |
| A0A4R1B4W5 | Cytobacillus praedii |
| A0A4R1MSC2 | Natranaerovirga hydrolytica |
| A0A4R1QUX6 | Fournierella massiliensis |
| A0A4R2BDS4 | Mesobacillus foraminis |
| A0A4R2L4A5 | Marinisporobacter balticus |
| A0A4R2LZB3 | Flavonifractor plautii DSM 6740 |
| A0A4R2M644 | Flavonifractor plautii DSM 6740 |
| A0A4R2NZI0 | Scopulibacillus darangshiensis |
| A0A4R2P5G1 | Scopulibacillus darangshiensis |
| A0A4R2RTW7 | Heliophilum fasciatum |
| A0A4R2TW42 | Serpentinicella alkaliphila |
| A0A4R2XET2 | Bacillus sp. OK085 |
| A0A4R3KDC4 | Tepidibacillus fermentans |
| A0A4R3KYR5 | Keratinibaculum paraultunense |
| A0A4R3MJI7 | Natranaerovirga pectinivora |
| A0A4R3NC51 | Melghiribacillus thermohalophilus |
| A0A4R4BE05 | Bacillus thuringiensis |
| A0A4R4D1D2 | Dehalobacter sp. 12DCB1 |
| A0A4R4D2N7 | Dehalobacter sp. 12DCB1 |
| A0A4R4EDT6 | Paenibacillus sp. 18JY21-1 |
| A0A4R5KQ08 | Paenibacillus piri |
| A0A4R5VX74 | Bacillus salipaludis |
| A0A4R5XQ65 | Jeotgalibacillus sp. S-D1 |
| A0A4R5ZXS8 | Rhodococcus qingshengii |
| A0A4R6AIJ4 | [Brevibacterium] frigoritolerans |
| A0A4R7KU89 | Fonticella tunisiensis |
| A0A4R7UA66 | Lysinibacillus sp. YR326 |
| A0A4R8GK73 | Cytobacillus oceanisediminis |
| A0A4R8H065 | Orenia marismortui |
| A0A4S2DDX9 | Clostridium sartagoforme |
| A0A4S4BXB4 | Bacillus sp. DSL-17 |
| A0A4T2AA21 | Marinifilum sp. JC120 |
| A0A4T9WFH9 | Tissierella creatinini |
| A0A4U0F9B4 | Cohnella pontilimi |
| A0A4U1D3C9 | Bacillus kyonggiensis |
| A0A4U2MJ25 | Peribacillus simplex |
| A0A4U2NZ50 | Bacillus cereus |
| A0A4U2Q3W0 | Paenibacillus terrae |
| A0A4U2X7T3 | Bacillus mycoides |
| A0A4U2Y852 | Brevibacillus antibioticus |
| A0A4U2Z3T0 | Lysinibacillus mangiferihumi |
| A0A4U3BHM8 | Bacillus cereus |
| A0A4U8YN28 | Desulfoluna butyratoxydans |
| A0A4U9R496 | Hathewaya histolytica |
| A0A4V1S4J2 | Desulfotomaculum aquiferis |
| A0A4V2KLV9 | Lysinibacillus sp. OL1 |
| A0A4V2ZP85 | Zhaonella formicivorans |
| A0A4V3WZT5 | Bacillus sp. HUB-I-004 |
| A0A4V6ENS0 | Ruminiclostridium herbifermentans |
| A0A4V6RSY5 | Bacillus timonensis |
| A0A4Y3P861 | Brevibacillus parabrevis |
| A0A4Y6F4U6 | Bacillus tropicus |
| A0A4Y7QTW7 | Bacillus sp. BH2 |
| A0A4Y7REQ7 | Pelotomaculum schinkii |
| A0A4Y7RRP9 | Pelotomaculum propionicicum |
| A0A4Y7S1W0 | Pelotomaculum sp. FP |
| A0A4Y8IFE1 | Filobacillus milosensis |
| A0A4Y8UDC8 | [Brevibacterium] frigoritolerans |
| A0A4Y9AER3 | Lentibacillus salicampi |
| A0A4Z0QMV9 | Desulfosporosinus sp. Sb-LF |
| A0A4Z0QX40 | Desulfosporosinus sp. Sb-LF |
| A0A4Z0R2S4 | Desulfosporosinus fructosivorans |
| A0A4Z0Y062 | Caproiciproducens galactitolivorans |
| A0A501UKT3 | Clostridium perfringens |
| A0A502HIG4 | Brevibacillus laterosporus |
| A0A506Q1V4 | Bacillus sp. |
| A0A511BYY0 | Rummeliibacillus stabekisii |
| A0A511UWU4 | Cerasibacillus quisquiliarum |
| A0A511VZV5 | Alkalibacillus haloalkaliphilus |
| A0A511X1E2 | Halolactibacillus alkaliphilus |
| A0A511ZA79 | Sporosarcina luteola |
| A0A511ZD33 | Oceanobacillus sojae |
| A0A513RP23 | Bacillus sp. S3 |
| A0A514LJ53 | Salicibibacter halophilus |
| A0A516QLF1 | Bacillus sp. BD59S |
| A0A517DUM4 | Sporomusa termitida |
| A0A517I9J6 | Brevibacillus brevis |
| A0A518VA86 | Brevibacillus laterosporus |
| A0A521J7V3 | Gottschalkiaceae bacterium |
| A0A521JSA7 | Anaerolineaceae bacterium |
| A0A540V5M7 | Ureibacillus terrenus |
| A0A542ARC9 | Clostridium sp. KNHs216 |
| A0A542GVQ9 | Microbacterium sp. SLBN-1 |
| A0A542SIH3 | Brevibacillus sp. AG162 |
| A0A544SU94 | Psychrobacillus soli |
| A0A544TC41 | Psychrobacillus lasiicapitis |
| A0A544TTL3 | Psychrobacillus vulpis |
| A0A544UF81 | Lysinibacillus sp. SDF0037 |
| A0A544UY67 | Lysinibacillus sp. SDF0063 |
| A0A549YED7 | Lentibacillus cibarius |
| A0A553KS21 | Brevibacillus sp. LEMMJ03 |
| A0A556C5L6 | Bacillus sp. HY001 |
| A0A556PH08 | Allobacillus sp. SKP4-8 |
| A0A556PUB0 | Allobacillus sp. SKP2-8 |
| A0A559J9F4 | Cohnella sp. G13 |
| A0A561CY49 | [Brevibacterium] frigoritolerans |
| A0A561D8A3 | Neobacillus bataviensis |
| A0A561PHU5 | Paenibacillus sp. 597 |
| A0A562GSH5 | Sporomusa sp. KB1 |
| A0A562HD75 | Desulfitobacterium sp. LBE |
| A0A562HLS9 | Desulfitobacterium sp. LBE |
| A0A562JFS2 | Cytobacillus oceanisediminis |
| A0A562JKH7 | Sedimentibacter saalensis |
| A0A562QS94 | Alkalihalobacillus nanhaiisediminis |
| A0A564TJ22 | Faecalibacterium prausnitzii |
| A0A564UBG7 | Faecalibacterium prausnitzii |
| A0A5A9E5S4 | Bacillus sp. CH30_1T |
| A0A5B0CPV8 | Sporosarcina sp. ANT_H38 |
| A0A5B0WWK9 | Paenibacillus sp. B2(2019) |
| A0A5B7TJG4 | Caloramator sp. E03 |
| A0A5B8PCI2 | Bacillus cereus |
| A0A5B8ZE44 | Bacillus dafuensis |
| A0A5B9YJF4 | Cellulosilyticum sp. WCF-2 |
| A0A5C0SF84 | Crassaminicella sp. SY095 |
| A0A5C1F943 | Bacillus sp. JAS24-2 |
| A0A5C1FPL0 | Bacillus mycoides |
| A0A5C4T074 | Paenibacillus hemerocallicola |
| A0A5C4ZHZ6 | Bacillus pacificus |
| A0A5C5A368 | Bacillus tropicus |
| A0A5C5AKT7 | Bacillus sp. CD3-5 |
| A0A5C6W520 | Metabacillus litoralis |
| A0A5D0CN84 | Paenibacillus faecis |
| A0A5D4KDH0 | Rossellomorea vietnamensis |
| A0A5D4KXV8 | Bacillus megaterium |
| A0A5D4M9V1 | Rossellomorea vietnamensis |
| A0A5D4NX00 | Rossellomorea vietnamensis |
| A0A5D4R9H1 | Bacillus infantis |
| A0A5D4S2R5 | Bacillus marisflavi |
| A0A5D4SPQ7 | Bacillus infantis |
| A0A5D4T384 | Bacillus horikoshii |
| A0A5D4T3R3 | Bacillus horikoshii |
| A0A5D4TUM1 | Bacillus aquimaris |
| A0A5D4TVK0 | Bacillus aquimaris |
| A0A5E9IIN0 | Bacillus thuringiensis F14-1 |
| A0A5J5H1Q3 | Bacillus endozanthoxylicus |
| A0A5J6PNV1 | Psychrobacillus sp. AK 1817 |
| A0A5J6SJ47 | Psychrobacillus glaciei |
| A0A5K5IH84 | Bacillus sp. FDAARGOS_235 |
| A0A5M8T9G6 | Bacillus cereus |
| A0A5M9GRJ1 | Bacillus paranthracis |
| A0A5M9NFY2 | Clostridium sp. HV4-5-A1G |
| A0A5P0YGD3 | Fictibacillus phosphorivorans |
| A0A5P3XFA4 | Paraclostridium bifermentans |
| A0A5P9HUS3 | Bacillus sp. THAF10 |
| A0A5P9XX08 | Bacillus cereus |
| A0A5Q2N8S1 | Lysinibacillus pakistanensis |
| A0A5Q2NA92 | Heliorestis convoluta |
| A0A5Q2TFM6 | Gracilibacillus sp. SCU50 |
| A0A5R9FDP0 | Alkalihalobacillus caeni |
| A0A5R9GE95 | Paenibacillus antri |
| A0A5R9MKS4 | Ruminococcus sp. KGMB03662 |
| A0A5S4ZU51 | Desulfallas thermosapovorans DSM 6562 |
| A0A5S5B0G2 | Thermosediminibacter litoriperuensis |
| A0A5S5CLE8 | Paenibacillus methanolicus |
| A0A644SYL7 | bioreactor metagenome |
| A0A644TWX1 | bioreactor metagenome |
| A0A644VJA4 | bioreactor metagenome |
| A0A644W404 | bioreactor metagenome |
| A0A644WWD5 | bioreactor metagenome |
| A0A644YQ45 | bioreactor metagenome |
| A0A645A740 | bioreactor metagenome |
| A0A645AAP8 | bioreactor metagenome |
| A0A645ABM7 | bioreactor metagenome |
| A0A645BWR7 | bioreactor metagenome |
| A0A653T5I2 | Bacillus sp. 349Y |
| A0A653VYQ2 | Bacillus mycoides |
| A0A657P3X8 | Bacillus cereus |
| A0A658J9E0 | Butyricicoccus sp. 1XD8-22 |
| A0A679H561 | Bacillus wiedmannii |
| A0A6A2T2F5 | Bacillus sp. B3-WWTP-C-10-D-3 |
| A0A6A4VE12 | Amphibalanus amphitrite |
| A0A6A8DII4 | Aquibacillus halophilus |
| A0A6A8KEA7 | Faecalibacterium prausnitzii |
| A0A6A8SI29 | Turicibacter sanguinis |
| A0A6A8V818 | Pseudoflavonifractor sp. BIOML-A6 |
| A0A6A8V830 | Pseudoflavonifractor sp. BIOML-A6 |
| A0A6B1UB15 | Bittarella massiliensis |
| A0A6B3TQF1 | Neobacillus thermocopriae |
| A0A6B3VUV5 | Bacillus aquiflavi |
| A0A6B3WAP4 | Clostridium botulinum |
| A0A6B3YFS9 | Clostridium botulinum |
| A0A6B3YNZ8 | Clostridium botulinum |
| A0A6B4A337 | Clostridium botulinum |
| A0A6B4GWU8 | Clostridium botulinum |
| A0A6B4HVR8 | Clostridium botulinum |
| A0A6B4IBA8 | Clostridium botulinum |
| A0A6B4JR13 | Clostridium botulinum |
| A0A6B4NDX9 | Clostridium botulinum |
| A0A6B4R0J7 | Clostridium botulinum |
| A0A6B4RTY4 | Clostridium botulinum |
| A0A6B4SA94 | Clostridium botulinum |
| A0A6B4VHG7 | Clostridium botulinum |
| A0A6B4YFX7 | Clostridium botulinum |
| A0A6B4ZZP1 | Clostridium botulinum |
| A0A6B8K598 | Bacillus sp. N3536 |
| A0A6B9YG48 | Virgibacillus sp. MSP4-1 |
| A0A6C0FZY1 | Paenibacillus lycopersici |
| A0A6D1SXL4 | Bacillus sp. BH32 |
| A0A6G1X5J6 | Salinibacillus xinjiangensis |
| A0A6G2CJC6 | Turicibacter sanguinis |
| A0A6G3LZ91 | Pseudoflavonifractor sp. 60 |
| A0A6G4Z9R8 | Clostridium perfringens |
| A0A6H0TKF9 | Bacillus thuringiensis serovar andalousiensis |
| A0A6H1NX95 | Bacillus megaterium |
| A0A6H1X3M4 | Romboutsia sp. CE17 |
| A0A6H3AAH2 | Bacillus anthracis |
| A0A6H9IE09 | Bacillus sp. AY1-10 |
| A0A6H9IPI7 | Bacillus sp. BPN334 |
| A0A6H9JK55 | Bacillus sp. AY2-1 |
| A0A6I0BHK4 | Bacillus sp. CH126_4D |
| A0A6I0F2A0 | Heliorestis acidaminivorans |
| A0A6I0FLR3 | Alkaliphilus pronyensis |
| A0A6I1FGY3 | Bacillus aerolatus |
| A0A6I2AHC9 | Bacillus thuringiensis |
| A0A6I2ETP7 | Bacillus thuringiensis |
| A0A6I2M920 | Bacillus idriensis |
| A0A6I2RNR0 | Flavonifractor plautii |
| A0A6I3Q644 | Ruthenibacterium lactatiformans |
| A0A6I3R4S9 | Pseudoflavonifractor sp. BIOML-A4 |
| A0A6I3RBZ7 | Pseudoflavonifractor sp. BIOML-A4 |
| A0A6I5ZLZ1 | Moorella glycerini |
| A0A6I5ZUP3 | Moorella glycerini |
| A0A6I6DHN4 | Candidatus Syntrophocurvum alkaliphilum |
| A0A6I6DMY8 | Candidatus Syntrophocurvum alkaliphilum |
| A0A6I6F0W7 | Clostridium bovifaecis |
| A0A6I6PVT3 | Bacillus marisflavi |
| A0A6I6UPJ5 | Rossellomorea vietnamensis |
| A0A6I6YBY6 | Bacillus paranthracis |
| A0A6I7FRP3 | Bacillus sp. NSP2.1 |
| A0A6I7GP76 | Flavonifractor plautii |
| A0A6I7XUL2 | Bacillus sp. SH7-1 |
| A0A6L3BP21 | Bacillus sp. TE8-1 |
| A0A6L3BXH4 | Bacillus sp. BB081 |
| A0A6L3V7K7 | Cytobacillus depressus |
| A0A6L3WQK2 | Bacillus cereus |
| A0A6L5B0U8 | Bacillus sp. ZZV12-4809 |
| A0A6L5LFR6 | Bacillus thuringiensis |
| A0A6L5PX38 | Bacillus sp. RIT694 |
| A0A6L6LRT4 | Ruthenibacterium lactatiformans |
| A0A6L8JRB9 | Virgibacillus halodenitrificans |
| A0A6L8P0Y2 | Bacillus anthracis |
| A0A6M0GZ06 | Clostridium senegalense |
| A0A6M0Q472 | Bacillus mesophilus |
| A0A6M0R927 | Clostridium niameyense |
| A0A6M0SVI1 | Clostridium botulinum |
| A0A6M0XFW4 | Clostridium botulinum |
| A0A6M0XZF1 | Clostridium sporogenes |
| A0A6M0YNS1 | Clostridium botulinum |
| A0A6M1VSI4 | Clostridium perfringens |
| A0A6M1WJW8 | Clostridium perfringens |
| A0A6M8DFM4 | Bacillus cereus |
| A0A6M8GS97 | Arthrobacter citreus |
| A0A6N2HHA1 | Bacillus sp. AR8-1 |
| A0A6N2V9I7 | uncultured Anaerotruncus sp |
| A0A6N3AVX5 | uncultured Clostridium sp |
| A0A6N3BM97 | Clostridium paraputrificum |
| A0A6N3CGG9 | uncultured Clostridium sp |
| A0A6N3D3J9 | Intestinibacter bartlettii |
| A0A6N3D3M1 | Flavonifractor plautii |
| A0A6N3GUU3 | Clostridium tertium |
| A0A6N7AYE7 | Firmicutes bacterium |
| A0A6N7B269 | Firmicutes bacterium |
| A0A6N7ISZ7 | Desulfofundulus thermobenzoicus |
| A0A6N7R0Z7 | Gracilibacillus thailandensis |
| A0A6N7TN75 | Faecalibacterium sp. BIOML-A1 |
| A0A6N7XNX5 | Tissierella pigra |
| A0A6N8B4R0 | Firmicutes bacterium |
| A0A6N8BDN2 | Firmicutes bacterium |
| A0A6N8BEI8 | Firmicutes bacterium |
| A0A6N8BNZ2 | Desulfovibrio sp |
| A0A6N8BU81 | Clostridiaceae bacterium |
| A0A6N8C219 | Clostridiaceae bacterium |
| A0A6N8CTT3 | Terrilactibacillus tamarindi |
| A0A6N8FIJ6 | Ornithinibacillus caprae |
| A0A6N8HS02 | Lentibacillus sp. JNUCC-1 |
| A0A6N9PSR5 | Neglecta sp. 59 |
| A0A6N9QHI8 | Clostridiales bacterium |
| A0A6P1AUE7 | bacterium LRH843 |
| A0A6P1HH38 | Pontibacillus sp. HMF3514 |
| A0A6P1MJ93 | Aminipila sp. CBA3637 |
| A0A6P1YC00 | Caloranaerobacter azorensis |
| A0A6S6XN32 | Ruminococcaceae bacterium BL-6 |
| A0A6S6YDF0 | Ruminococcaceae bacterium BL-4 |
| A0A7C1K7S8 | Firmicutes bacterium |
| A0A7C6CL76 | Firmicutes bacterium |
| A0A7C6DDK9 | Clostridiales bacterium |
| A0A7C6DDZ0 | Clostridiaceae bacterium |
| A0A7C6GDH1 | Firmicutes bacterium |
| A0A7C6GNF6 | Firmicutes bacterium |
| A0A7C6GQS9 | Clostridiales bacterium |
| A0A7C6H0R5 | Firmicutes bacterium |
| A0A7C6HBU7 | Firmicutes bacterium |
| A0A7C6I248 | bacterium |
| A0A7C6I7L3 | Clostridia bacterium |
| A0A7C6J2V5 | Firmicutes bacterium |
| A0A7C6JG03 | Halanaerobiaceae bacterium |
| A0A7C6JTG4 | Firmicutes bacterium |
| A0A7C6K588 | Tissierellia bacterium |
| A0A7C6KMN9 | Firmicutes bacterium |
| A0A7C6KMV2 | Firmicutes bacterium |
| A0A7C6KQF9 | Firmicutes bacterium |
| A0A7C6KQL7 | Peptococcaceae bacterium |
| A0A7C6KT62 | Peptococcaceae bacterium |
| A0A7C6KVP1 | Clostridia bacterium |
| A0A7C6LA05 | Firmicutes bacterium |
| A0A7C6LGA3 | Firmicutes bacterium |
| A0A7C6LMA9 | Syntrophomonadaceae bacterium |
| A0A7C6LXM1 | Clostridiales bacterium |
| A0A7C6MQM7 | Natronincola sp |
| A0A7C6MWU2 | Clostridiales bacterium |
| A0A7C6MWY4 | Thermoanaerobacterales bacterium |
| A0A7C6N070 | Clostridiales bacterium |
| A0A7C6NBF0 | Peptococcaceae bacterium |
| A0A7C6PA07 | Syntrophomonadaceae bacterium |
| A0A7C6PC58 | Syntrophomonadaceae bacterium |
| A0A7C6QL67 | Syntrophomonadaceae bacterium |
| A0A7C6QMM1 | Firmicutes bacterium |
| A0A7C6QZ05 | Syntrophomonadaceae bacterium |
| A0A7C6R2F0 | Desulfotomaculum sp |
| A0A7C6R779 | Clostridiales bacterium |
| A0A7C6RBJ1 | Firmicutes bacterium |
| A0A7C6S8L4 | Firmicutes bacterium |
| A0A7C6UCY2 | Firmicutes bacterium |
| A0A7C6UP72 | Clostridia bacterium |
| A0A7C6UQ56 | Firmicutes bacterium |
| A0A7C6UX56 | Clostridiaceae bacterium |
| A0A7C6V1S1 | Syntrophomonadaceae bacterium |
| A0A7C6V2J3 | Syntrophomonadaceae bacterium |
| A0A7C6VSC9 | Epulopiscium sp |
| A0A7C6VV10 | Firmicutes bacterium |
| A0A7C6WKY1 | Clostridiales bacterium |
| A0A7C6WXJ4 | Thermoanaerobacterales bacterium |
| A0A7C6X1Y6 | Firmicutes bacterium |
| A0A7C6X4B9 | Firmicutes bacterium |
| A0A7C6XXJ1 | Clostridiales bacterium |
| A0A7C6Y7E3 | Firmicutes bacterium |
| A0A7C6YKV7 | Thermoanaerobacterales bacterium |
| A0A7C6YXI0 | Firmicutes bacterium |
| A0A7C6ZDK9 | Firmicutes bacterium |
| A0A7C6ZFI6 | Firmicutes bacterium |
| A0A7C6ZH21 | Clostridia bacterium |
| A0A7C6ZIJ8 | Clostridia bacterium |
| A0A7C6ZKX3 | Syntrophaceticus sp |
| A0A7C6ZKZ4 | Syntrophaceticus sp |
| A0A7C6ZPG7 | Firmicutes bacterium |
| A0A7C7AH63 | Tissierellia bacterium |
| A0A7C7D3B7 | Desulfitobacterium dehalogenans |
| A0A7C7D8F6 | Desulfitobacterium dehalogenans |
| A0A7C7DDM2 | Firmicutes bacterium |
| A0A7C7E6Q8 | Clostridiales bacterium |
| A0A7C7E9Q5 | Thermoanaerobacterales bacterium |
| A0A7C7EEY8 | Clostridiales bacterium |
| A0A7C7ES76 | Tissierellia bacterium |
| A0A7C8HEL4 | Defluviitalea raffinosedens |
| A0A7C9H7Q3 | Firmicutes bacterium |
| A0A7C9LNW5 | Firmicutes bacterium |
| A0A7D3YTS7 | Bacillus cereus |
| A0A7D6W0W0 | Clostridium intestinale |
| A0A7G6DYX6 | Thermoanaerosceptrum fracticalcis |
| A0A7G7FZN6 | Metabacillus sp. KUDC1714 |
| A0A7G8U143 | Brevibacterium sp. PAMC23299 |
| A0A7G8UM29 | Paenibacillus sp. PAMC21692 |
| A0A7G8X521 | Sporosarcina sp. resist |
| A0A7G9B1C0 | Oscillibacter sp. NSJ-62 |
| A0A7G9WA91 | Alkalicella caledoniensis |
| A0A7H8S7I8 | Lentibacillus sp. CBA3610 |
| A0A7H8SBS3 | Lentibacillus sp. CBA3610 |
| A0A7I8CZX6 | Solibaculum mannosilyticum |
| A0A7M1PWT5 | Clostridium sp. 'deep sea' |
| A0A7M1SCY3 | Bacillus sp. HD4P25 |
| A0A7M2AUG6 | Brevibacillus sp. JNUCC-41 |
| A0A7M2B8L1 | Brevibacterium sp. JNUCC-42 |
| A0A7M2QY21 | Viridibacillus sp. JNUCC-6 |
| A0A7S7J0H7 | Clostridiales bacterium |
| A0A7S7L5L3 | Anaerobacillus isosaccharinicus |
| A0A7T0GK46 | Sarcina sp. JB2 |
| A0A7T2CM09 | Lysinibacillus sp. JNUCC-51 |
| A0A7T2CWN8 | Lysinibacillus sp. JNUCC-52 |
| A0A7T3MQN5 | Bacillus thuringiensis |
| A0A7T5JMM6 | Brevibacillus sp. FJAT-54423 |
| A0A7T6Z6I6 | Salicibibacter cibarius |
| A0A7T6ZDJ5 | Salicibibacter cibi |
| A0A7T8FS40 | Lysinibacillus sp. FJAT-51161 |
| A0A7T8LT30 | Bacillus sp. TK-2 |
| A0A7T9LNC4 | Weizmannia coagulans |
| A0A7T9M1K1 | Peribacillus psychrosaccharolyticus |
| A0A7T9WWL7 | Bacillus cereus |
| A0A7U1C0Y7 | Bacillus vini |
| A0A7U1GM15 | Bacillus oleronius |
| A0A7U5GXH5 | Sporosarcina sp. P37 |
| A0A7U6BS14 | Peribacillus butanolivorans |
| A0A7U6BYI3 | Bacillus anthracis |
| A0A7U6KG44 | Lachnospiraceae bacterium KM106-2 |
| A0A7U9C9P3 | Clostridium sporogenes (strain ATCC 7955 / DSM 767 / NBRC 16411 / NCIMB 8053 / NCTC 8594 / PA 3679) |
| A0A7U9IA45 | Clostridium sp. (strain ATCC 29733 / VPI C48-50) |
| A0A7V3T2L0 | Clostridia bacterium |
| A0A7V6GCS0 | Clostridia bacterium |
| A0A7V6HMG2 | Bacilli bacterium |
| A0A7V6JHG5 | Clostridiaceae bacterium |
| A0A7V6M949 | bacterium |
| A0A7V6MTU2 | Thermoanaerobacterales bacterium |
| A0A7V6MY74 | Thermoanaerobacterales bacterium |
| A0A7V6PWL0 | Clostridiaceae bacterium |
| A0A7V6QRQ9 | Bacillales bacterium |
| A0A7V6RKU2 | Syntrophomonadaceae bacterium |
| A0A7V6RN39 | Syntrophomonadaceae bacterium |
| A0A7V6TLZ4 | Tissierellia bacterium |
| A0A7V6TV71 | Clostridiaceae bacterium |
| A0A7V6UD60 | Thermoanaerobacterales bacterium |
| A0A7V6UY66 | Clostridiaceae bacterium |
| A0A7V6X306 | Clostridia bacterium |
| A0A7V6Y6I5 | Thermoanaerobacterales bacterium |
| A0A7V6YDU2 | Clostridia bacterium |
| A0A7V6ZW88 | Clostridia bacterium |
| A0A7V7BGU3 | Clostridia bacterium |
| A0A7V7BLQ0 | Bacillus sp. |
| A0A7V7HHU0 | Bacillus sp. AR2-1 |
| A0A7V7HIR9 | Bacillus sp. SH5-2 |
| A0A7V7IBB4 | Bacillus sp. BB56-3 |
| A0A7V7L1U8 | Bacillus sp. BF2-3 |
| A0A7V7LCJ6 | Bacillus sp. BB51/4 |
| A0A7V7V767 | Bacillus luti |
| A0A7V7YXD2 | Bacillus sp. B1-WWTP-T-0.5-Post-4 |
| A0A7V8ICA6 | Clostridium sp. NCR |
| A0A7W0HLA7 | Desulfosalsimonas propionicica |
| A0A7W3SPF6 | Fontibacillus solani |
| A0A7W4L921 | Bacillus sp. APMAM |
| A0A7W5B4Z0 | Paenibacillus phyllosphaerae |
| A0A7W7X7L1 | Bacillus toyonensis |
| A0A7X0HT98 | Bacillus benzoevorans |
| A0A7X0RQH6 | Cohnella nanjingensis |
| A0A7X0S9B5 | Clostridium gasigenes |
| A0A7X0VL37 | Clostridium algidicarnis |
| A0A7X1IC48 | Bittarella massiliensis |
| A0A7X2AYS0 | Bacillus thuringiensis |
| A0A7X2DN79 | Bacillus thuringiensis |
| A0A7X2EBL1 | Bacillus thuringiensis |
| A0A7X2IWM3 | Bacillus lacus |
| A0A7X2JV26 | Bacillus thuringiensis |
| A0A7X2MZ49 | Inconstantimicrobium porci |
| A0A7X2Z2I4 | Paenibacillus woosongensis |
| A0A7X3DJR4 | Oscillibacter sp |
| A0A7X3IEC6 | Paenibacillus sp. HJL G12 |
| A0A7X5PCB6 | Clostridium sporogenes |
| A0A7X6SBF8 | Tissierellia bacterium |
| A0A7X6UIS6 | Syntrophomonadaceae bacterium |
| A0A7X6UIZ4 | Syntrophomonadaceae bacterium |
| A0A7X6VD94 | Clostridiaceae bacterium |
| A0A7X6VI85 | Clostridiaceae bacterium |
| A0A7X6VSZ7 | Syntrophomonadaceae bacterium |
| A0A7X6WLS9 | Syntrophomonadaceae bacterium |
| A0A7X6XUD8 | Thermoanaerobacterales bacterium |
| A0A7X6Y3N0 | Clostridiaceae bacterium |
| A0A7X6Y8B8 | Clostridia bacterium |
| A0A7X7AP46 | Clostridia bacterium |
| A0A7X7CH54 | Clostridiaceae bacterium |
| A0A7X7EJ03 | Clostridiaceae bacterium |
| A0A7X7GZ90 | Bacilli bacterium |
| A0A7X7MUM9 | Clostridia bacterium |
| A0A7X7MVJ8 | Clostridia bacterium |
| A0A7X7NZ92 | Syntrophomonadaceae bacterium |
| A0A7X7Q1I2 | Bacilli bacterium |
| A0A7X7R440 | Syntrophomonadaceae bacterium |
| A0A7X7R4F8 | Syntrophomonadaceae bacterium |
| A0A7X7RDS6 | Clostridia bacterium |
| A0A7X7S9Z9 | Syntrophomonadaceae bacterium |
| A0A7X7SXE4 | Clostridiaceae bacterium |
| A0A7X7TKW1 | Syntrophomonadaceae bacterium |
| A0A7X7VXL3 | Clostridiaceae bacterium |
| A0A7X7W023 | Clostridiaceae bacterium |
| A0A7X7X2I2 | Clostridiaceae bacterium |
| A0A7X7XYV0 | Clostridium sp. |
| A0A7X7Y1U9 | Tissierellia bacterium |
| A0A7X7YMR1 | Syntrophomonadaceae bacterium |
| A0A7X8ATQ3 | Clostridium sp. |
| A0A7X8AXV7 | Bacilli bacterium |
| A0A7X8DGT4 | Peptococcaceae bacterium |
| A0A7X8DRN7 | Halanaerobiaceae bacterium |
| A0A7X8E5I6 | Tissierellia bacterium |
| A0A7X8EB61 | Clostridia bacterium |
| A0A7X8EQQ8 | Epulopiscium sp |
| A0A7X8G314 | Tissierellia bacterium |
| A0A7X8HVB8 | Epulopiscium sp |
| A0A7X8ILD3 | Syntrophomonadaceae bacterium |
| A0A7X8IYI9 | Tissierellia bacterium |
| A0A7X8J0G8 | Clostridiaceae bacterium |
| A0A7X8JGE8 | Clostridiaceae bacterium |
| A0A7X8JPL1 | Epulopiscium sp |
| A0A7X8JV28 | Clostridiaceae bacterium |
| A0A7X8JVM6 | Clostridiaceae bacterium |
| A0A7X8K0M5 | Clostridia bacterium |
| A0A7X8KU57 | Peptococcaceae bacterium |
| A0A7X8KVQ6 | Peptococcaceae bacterium |
| A0A7X8KXH3 | Tissierellia bacterium |
| A0A7X8LBC6 | Epulopiscium sp |
| A0A7X8LE64 | Tissierellia bacterium |
| A0A7X8LKM9 | Syntrophomonadaceae bacterium |
| A0A7X8LLI0 | Syntrophomonadaceae bacterium |
| A0A7X8LZ60 | Clostridiaceae bacterium |
| A0A7X8M5S1 | Clostridiaceae bacterium |
| A0A7X8ME38 | Clostridium sp. |
| A0A7X8MKV5 | Syntrophomonadaceae bacterium |
| A0A7X8MKW5 | Syntrophomonadaceae bacterium |
| A0A7X8MTC7 | Clostridia bacterium |
| A0A7X8QKK6 | Clostridiaceae bacterium |
| A0A7X8QZ39 | Clostridiaceae bacterium |
| A0A7X8RI82 | Bacillus sp. RO1 |
| A0A7X8U432 | Syntrophomonadaceae bacterium |
| A0A7X8U9N5 | Syntrophomonadaceae bacterium |
| A0A7X8UAT8 | Syntrophomonadaceae bacterium |
| A0A7X8UAU5 | Syntrophomonadaceae bacterium |
| A0A7X8V2N3 | Clostridia bacterium |
| A0A7X8V4D6 | Clostridia bacterium |
| A0A7X8VE48 | Thermoanaerobacteraceae bacterium |
| A0A7X8X4U8 | Clostridia bacterium |
| A0A7X8YPE3 | Syntrophomonadaceae bacterium |
| A0A7X9A2W8 | Clostridia bacterium |
| A0A7X9A3N7 | Clostridia bacterium |
| A0A7X9AK19 | Clostridiaceae bacterium |
| A0A7X9BDX9 | Peptococcaceae bacterium |
| A0A7X9BN40 | Syntrophomonadaceae bacterium |
| A0A7X9BY17 | Desulfitobacterium sp |
| A0A7X9BYB6 | Desulfitobacterium sp |
| A0A7X9CMI0 | Tissierellia bacterium |
| A0A7X9GIJ8 | Syntrophomonadaceae bacterium |
| A0A7X9GX79 | Tissierellia bacterium |
| A0A7X9H752 | Clostridium sp. |
| A0A7X9HV19 | Clostridiaceae bacterium |
| A0A7X9M2Y9 | Bacillus sp. DNRA2 |
| A0A7X9MUW7 | Psychrobacillus sp. BL-248-WT-3 |
| A0A7X9XEY1 | Clostridium sp. SM-530-WT-3G |
| A0A7Y0HRM8 | Clostridium sp. P21 |
| A0A7Y0L2Y7 | Sulfobacillus sp. DSM 109850 |
| A0A7Y0QFI2 | Clostridioides difficile |
| A0A7Y0R3C4 | Clostridioides difficile |
| A0A7Y3V7F3 | Clostridium cochlearium |
| A0A7Y8S572 | Bacillus sp. |
| A0A7Y9BFJ1 | Bacillus sp. EB106-08-02-XG196 |
| A0A7Z0PMB4 | Bacillus sp. Gen3 |
| A0A7Z1FW67 | Bacillus pseudomycoides |
| A0A7Z1G0F8 | Bacillus pseudomycoides |
| A0A7Z1H6N7 | Bacillus pseudomycoides |
| A0A7Z8GW09 | Bacillus sp. 007/AIA-02/001 |
| A0A7Z8RK23 | Bacillus sp. AY18-3 |
| A0A7Z8RWA2 | Bacillus sp. AR13-1 |
| A0A7Z8S410 | Bacillus sp. BF9-10 |
| A0A800N994 | Cytobacillus firmus |
| A0A806I8Q0 | Bacillus thuringiensis HD-789 |
| A0A806LMP8 | Lysinibacillus varians |
| A0A806Q4X9 | Clostridium botulinum CDC_297 |
| A0A810Q5T9 | Oscillibacter sp. MM59 |
| A0A822Q2Z0 | Paeniclostridium sordellii |
| A0A822ULY4 | Clostridium perfringens |
| A0A826HM10 | Bacillus thuringiensis Bt407 |
| A0A828S1L8 | Turicibacter sp. HGF1 |
| A0A828XKZ7 | Bacillus cereus BAG4X12-1 |
| A0A828ZDY2 | Lysinibacillus fusiformis ZB2 |
| A0A829ZSL2 | Thermanaeromonas sp. C210 |
| A0A829ZUL2 | Thermanaeromonas sp. C210 |
| A0A833HN03 | Alkaliphilus serpentinus |
| A0A837GI25 | Bacillaceae bacterium MTCC 10057 |
| A0A837KMQ6 | Brevibacillus formosus |
| A0A837YUD3 | Bacillus badius |
| A0A838X3K7 | Brevibacillus halotolerans |
| A0A839TQS9 | Paenibacillus rhizosphaerae |
| A0A840KLU9 | Sporosarcina luteola |
| A0A840PPP3 | Ureibacillus thermosphaericus |
| A0A840QSY3 | Texcoconibacillus texcoconensis |
| A0A841KV83 | Anaerosolibacter carboniphilus |
| A0A841Q0X3 | Geomicrobium halophilum |
| A0A841Q651 | Salirhabdus euzebyi |
| A0A844DH03 | Faecalibacterium prausnitzii |
| A0A844DYE9 | Faecalibacterium prausnitzii |
| A0A844E9W1 | Faecalibacterium sp. BIOML-A3 |
| A0A844FHR5 | Anaerosalibacter bizertensis |
| A0A844K0X8 | Pseudoflavonifractor sp. BIOML-A18 |
| A0A844K2L4 | Pseudoflavonifractor sp. BIOML-A18 |
| A0A844LWF0 | Virgibacillus dakarensis |
| A0A845QQR3 | Anaerotruncus colihominis |
| A0A845QT30 | Senegalia massiliensis |
| A0A845R4V3 | Colidextribacter sp. OB.20 |
| A0A845RA57 | Colidextribacter sp. OB.20 |
| A0A845RL93 | Anaerotruncus colihominis |
| A0A845RRR7 | Dehalobacter sp. 4CP |
| A0A845RT85 | Dehalobacter sp. 4CP |
| A0A846HS47 | Clostridium botulinum |
| A0A846J4Y7 | Clostridium botulinum |
| A0A846JZ08 | Clostridium botulinum |
| A0A846TKE1 | Mesobacillus selenatarsenatis |
| A0A847AWZ6 | Tissierellia bacterium |
| A0A847CA00 | Oscillospiraceae bacterium |
| A0A847KCU1 | Peptococcaceae bacterium |
| A0A847NAY7 | Gracilibacteraceae bacterium |
| A0A847NCY6 | Gracilibacteraceae bacterium |
| A0A847NMB1 | Tissierellia bacterium |
| A0A847QJ83 | Veillonellaceae bacterium |
| A0A847QK74 | Peptococcaceae bacterium |
| A0A847QQQ4 | Peptococcaceae bacterium |
| A0A847QRM5 | Epulopiscium sp |
| A0A847WA91 | Papillibacter sp |
| A0A847WEB3 | Epulopiscium sp |
| A0A847XCR4 | Tissierellia bacterium |
| A0A848BLW0 | Paraclostridium bifermentans |
| A0A850EN10 | Paenibacillus sp. JW14 |
| A0A852TDM5 | Neobacillus niacini |
| A0A852UDW1 | Sporosarcina sp. JAI121 |
| A0A853WAF5 | Moorella thermoacetica |
| A0A853XD93 | Bacillus sp. L27 |
| A0A853XNP5 | Bacillus pacificus |
| A0A853Y7D0 | Bacillus mobilis |
| A0A854AZS9 | Bacillus toyonensis |
| A0A854DGV6 | Bacillus thuringiensis |
| A0A854KQ33 | Bacillus thuringiensis serovar shandongiensis |
| A0A854U9Y6 | Peribacillus simplex |
| A0A855AQQ2 | Bacillus thuringiensis |
| A0A855AUN2 | Bacillus cereus |
| A0A855BI38 | Bacillus cereus |
| A0A855E4Z1 | Bacillus thuringiensis |
| A0A855KY33 | Bacillus sp. AKBS9 |
| A0A857DDP5 | Dehalobacter restrictus |
| A0A857DMW9 | Dehalobacter restrictus |
| A0A858BUP0 | Aminipila butyrica |
| A0A858BWL1 | Aminipila butyrica |
| A0A8A3NRN9 | thermophilic bacterium 3443-3Ac |
| A0A8A5Z6E9 | Cohnella sp. LGH |
| A0A8A7KNN4 | Halanaerobiaceae bacterium NS-1 |
| A0A8A9G9B2 | Bacillus cytotoxicus |
| A0A8B2VMS5 | Bacillus sp. dmp5 |
| A0A8B4BZD4 | Bacillus coagulans DSM 1 = ATCC 7050 |
| A0A8B5XTP7 | Peribacillus simplex |
| A0A8B6J9B3 | Clostridioides difficile |
| A0PYH5 | Clostridium novyi (strain NT) |
| A0R959 | Bacillus thuringiensis (strain Al Hakam) |
| A1HS32 | Thermosinus carboxydivorans Nor1 |
| A3DCJ3 | Acetivibrio thermocellus (strain ATCC 27405 / DSM 1237 / JCM 9322 / NBRC 103400 / NCIMB 10682 / NRRL B-4536 / VPI 7372) |
| A4J8C7 | Desulfotomaculum reducens (strain MI-1) |
| A5D4E2 | Pelotomaculum thermopropionicum (strain DSM 13744 / JCM 10971 / SI) |
| A5D4H5 | Pelotomaculum thermopropionicum (strain DSM 13744 / JCM 10971 / SI) |
| A5I764 | Clostridium botulinum (strain Hall / ATCC 3502 / NCTC 13319 / Type A) |
| A6CL50 | Bacillus sp. SG-1 |
| A6P157 | Pseudoflavonifractor capillosus ATCC 29799 |
| A6P165 | Pseudoflavonifractor capillosus ATCC 29799 |
| A6TSS7 | Alkaliphilus metalliredigens (strain QYMF) |
| A7GIS6 | Clostridium botulinum (strain Langeland / NCTC 10281 / Type F) |
| A7GKN8 | Bacillus cytotoxicus (strain DSM 22905 / CIP 110041 / 391-98 / NVH 391-98) |
| A7VP38 | [Clostridium] leptum DSM 753 |
| A8MF94 | Alkaliphilus oremlandii (strain OhILAs) |
| A8SGA0 | Faecalibacterium prausnitzii M21/2 |
| A9KQQ0 | Lachnoclostridium phytofermentans (strain ATCC 700394 / DSM 18823 / ISDg) |
| A9VRL8 | Bacillus mycoides (strain KBAB4) |
| B0A768 | Intestinibacter bartlettii DSM 16795 |
| B0PGQ2 | Anaerotruncus colihominis DSM 17241 |
| B1B7L8 | Clostridium botulinum C str. Eklund |
| B1BUY9 | Clostridium perfringens E str. JGS1987 |
| B1HZY4 | Lysinibacillus sphaericus (strain C3-41) |
| B1I0Y4 | Desulforudis audaxviator (strain MP104C) |
| B1IFH8 | Clostridium botulinum (strain Okra / Type B1) |
| B1L220 | Clostridium botulinum (strain Loch Maree / Type A3) |
| B1V147 | Clostridium perfringens D str. JGS1721 |
| B2A234 | Natranaerobius thermophilus (strain ATCC BAA-1301 / DSM 18059 / JW/NM-WN-LF) |
| B2TR81 | Clostridium botulinum (strain Eklund 17B / Type B) |
| B7H7E6 | Bacillus cereus (strain B4264) |
| B7HT42 | Bacillus cereus (strain AH187) |
| B7IV22 | Bacillus cereus (strain G9842) |
| B7JN48 | Bacillus cereus (strain AH820) |
| B8D024 | Halothermothrix orenii (strain H 168 / OCM 544 / DSM 9562) |
| B8FWH0 | Desulfitobacterium hafniense (strain DSM 10664 / DCB-2) |
| B8FYY0 | Desulfitobacterium hafniense (strain DSM 10664 / DCB-2) |
| B8I518 | Ruminiclostridium cellulolyticum (strain ATCC 35319 / DSM 5812 / JCM 6584 / H10) |
| B9J2I6 | Bacillus cereus (strain Q1) |
| C0EC80 | [Clostridium] methylpentosum DSM 5476 |
| C0GEK8 | Dethiobacter alkaliphilus AHT 1 |
| C0GKQ5 | Dethiobacter alkaliphilus AHT 1 |
| C0Z7Y0 | Brevibacillus brevis (strain 47 / JCM 6285 / NBRC 100599) |
| C1FM06 | Clostridium botulinum (strain Kyoto / Type A2) |
| C2MFH9 | Bacillus cereus m1293 |
| C2NCC9 | Bacillus cereus BGSC 6E1 |
| C2NTK7 | Bacillus cereus 172560W |
| C2P9Q4 | Bacillus wiedmannii |
| C2Q4Q4 | Bacillus mycoides |
| C2R2U8 | Bacillus cereus m1550 |
| C2RYC7 | Bacillus cereus BDRD-ST26 |
| C2SET4 | Bacillus cereus BDRD-ST196 |
| C2SVM5 | Bacillus cereus BDRD-Cer4 |
| C2U8N8 | Bacillus cereus Rock1-15 |
| C2UQ90 | Bacillus cereus Rock3-28 |
| C2V6L5 | Bacillus cereus Rock3-29 |
| C2VND4 | Bacillus cereus Rock3-42 |
| C2W3H0 | Bacillus cereus Rock3-44 |
| C2WH64 | Bacillus cereus Rock4-2 |
| C2X6J6 | Bacillus cereus F65185 |
| C2XNT4 | Bacillus mycoides |
| C2YLE7 | Bacillus cereus AH1271 |
| C2Z2I2 | Bacillus cereus AH1272 |
| C3AGV8 | Bacillus pseudomycoides |
| C3BF79 | Bacillus pseudomycoides DSM 12442 |
| C3DEH4 | Bacillus thuringiensis serovar sotto str. T04001 |
| C3DYB8 | Bacillus thuringiensis serovar pakistani str. T13001 |
| C3FXV5 | Bacillus thuringiensis serovar andalousiensis BGSC 4AW1 |
| C3GDL8 | Bacillus thuringiensis serovar pondicheriensis BGSC 4BA1 |
| C3GVU6 | Bacillus thuringiensis serovar huazhongensis BGSC 4BD1 |
| C6BUC2 | Desulfovibrio salexigens (strain ATCC 14822 / DSM 2638 / NCIMB 8403 / VKM B-1763) |
| C6CZL4 | Paenibacillus sp. (strain JDR-2) |
| C6PNR2 | Clostridium carboxidivorans P7 |
| C7H4L3 | Faecalibacterium prausnitzii (strain DSM 17677 / JCM 31915 / A2-165) |
| D3EJ60 | Geobacillus sp. (strain Y412MC10) |
| D3FTK9 | Alkalihalobacillus pseudofirmus (strain ATCC BAA-2126 / JCM 17055 / OF4) |
| D4K392 | Faecalibacterium prausnitzii L2-6 |
| D4K5Q4 | Faecalibacterium prausnitzii SL3/3 |
| D4LCC1 | Ruminococcus champanellensis (strain DSM 18848 / JCM 17042 / KCTC 15320 / 18P13) |
| D5Q7I4 | Clostridioides difficile NAP08 |
| D5XCB8 | Thermincola potens (strain JR) |
| D7CMV3 | Syntrophothermus lipocalidus (strain DSM 12680 / TGB-C1) |
| D7CMW6 | Syntrophothermus lipocalidus (strain DSM 12680 / TGB-C1) |
| D7CPI5 | Syntrophothermus lipocalidus (strain DSM 12680 / TGB-C1) |
| D8GZN9 | Bacillus cereus var. anthracis (strain CI) |
| D9QSU7 | Acetohalobium arabaticum (strain ATCC 49924 / DSM 5501 / Z-7288) |
| D9S3F2 | Thermosediminibacter oceani (strain ATCC BAA-1034 / DSM 16646 / JW/IW-1228P) |
| D9SU30 | Clostridium cellulovorans (strain ATCC 35296 / DSM 3052 / OCM 3 / 743B) |
| E1JTH6 | Desulfovibrio fructosivorans JJ |
| E2ZIP0 | Faecalibacterium cf. prausnitzii KLE1255 |
| E3GYH6 | Methanothermus fervidus (strain ATCC 43054 / DSM 2088 / JCM 10308 / V24 S) |
| E5WH22 | Bacillus sp. 2_A_57_CT2 |
| E6SGM7 | Thermaerobacter marianensis (strain ATCC 700841 / DSM 12885 / JCM 10246 / 7p75a) |
| E6U242 | Bacillus cellulosilyticus (strain ATCC 21833 / DSM 2522 / FERM P-1141 / JCM 9156 / N-4) |
| E6U7S8 | Ethanoligenens harbinense (strain DSM 18485 / JCM 12961 / CGMCC 1.5033 / YUAN-3) |
| E6UIA5 | Ruminococcus albus (strain ATCC 27210 / DSM 20455 / JCM 14654 / NCDO 2250 / 7) |
| E9SC08 | Ruminococcus albus 8 |
| F0T282 | Syntrophobotulus glycolicus (strain DSM 8271 / FlGlyR) |
| F1TD32 | Ruminiclostridium papyrosolvens DSM 2782 |
| F2F5B4 | Solibacillus silvestris (strain StLB046) |
| F2JMP3 | Cellulosilyticum lentocellum (strain ATCC 49066 / DSM 5427 / NCIMB 11756 / RHM5) |
| F3M7Z8 | Paenibacillus sp. HGF5 |
| F3ZYE5 | Mahella australiensis (strain DSM 15567 / CIP 107919 / 50-1 BON) |
| F4LUV1 | Tepidanaerobacter acetatoxydans (strain DSM 21804 / JCM 16047 / Re1) |
| F4XDD8 | Ruminococcaceae bacterium D16 |
| F5L908 | Caldalkalibacillus thermarum (strain TA2.A1) |
| F6B7M9 | Desulfotomaculum nigrificans (strain DSM 14880 / VKM B-2319 / CO-1-SRB) |
| F6DV32 | Desulfotomaculum ruminis (strain ATCC 23193 / DSM 2154 / NCIMB 8452 / DL) |
| F7YZF7 | Bacillus coagulans (strain 2-6) |
| F9DRH8 | Sporosarcina newyorkensis 2681 |
| G2FZ51 | Desulfosporosinus sp. OT |
| G2IFS5 | Candidatus Arthromitus sp. SFB-rat-Yit |
| G2TI38 | Bacillus coagulans 36D1 |
| G4HDB3 | Paenibacillus lactis 154 |
| G4KNY6 | Oscillibacter valericigenes (strain DSM 18026 / NBRC 101213 / Sjm18-20) |
| G7VTU2 | Paenibacillus terrae (strain HPL-003) |
| G7WGC8 | Desulfosporosinus orientis (strain ATCC 19365 / DSM 765 / NCIMB 8382 / VKM B-1628 / Singapore I) |
| G8LX97 | Hungateiclostridium clariflavum (strain DSM 19732 / NBRC 101661 / EBR45) |
| G8LXQ4 | Hungateiclostridium clariflavum (strain DSM 19732 / NBRC 101661 / EBR45) |
| G8U0K6 | Sulfobacillus acidophilus (strain ATCC 700253 / DSM 10332 / NAL) |
| G9QEC9 | Bacillus sp. 7_6_55CFAA_CT2 |
| G9RY24 | Subdoligranulum sp. 4_3_54A2FAA |
| G9RY65 | Subdoligranulum sp. 4_3_54A2FAA |
| G9XMB5 | Desulfitobacterium hafniense DP7 |
| G9XTI7 | Desulfitobacterium hafniense DP7 |
| G9YNB0 | Flavonifractor plautii ATCC 29863 |
| G9YTS8 | Flavonifractor plautii ATCC 29863 |
| H0UHL3 | Brevibacillus laterosporus GI-9 |
| H1CJV2 | Lachnospiraceae bacterium 7_1_58FAA |
| H1CM65 | Lachnospiraceae bacterium 7_1_58FAA |
| H2JGD2 | Clostridium sp. BNL1100 |
| H5Y0H2 | Desulfosporosinus youngiae DSM 17734 |
| H5Y4C0 | Desulfosporosinus youngiae DSM 17734 |
| I3EBS4 | Bacillus methanolicus (strain MGA3 / ATCC 53907) |
| I4A9C6 | Desulfitobacterium dehalogenans (strain ATCC 51507 / DSM 9161 / JW/IU-DC1) |
| I4AEC7 | Desulfitobacterium dehalogenans (strain ATCC 51507 / DSM 9161 / JW/IU-DC1) |
| I4DCA6 | Desulfosporosinus acidiphilus (strain DSM 22704 / JCM 16185 / SJ4) |
| I7K7N2 | Caloramator australicus RC3 |
| I8UKI9 | Fictibacillus macauensis ZFHKF-1 |
| J0MVU4 | Clostridium sp. MSTE9 |
| J2HF29 | Brevibacillus sp. CF112 |
| J3A0M7 | Brevibacillus sp. BC25 |
| J3UQG8 | Bacillus thuringiensis HD-771 |
| J7J1X7 | Desulfosporosinus meridiei (strain ATCC BAA-275 / DSM 13257 / KCTC 12902 / NCIMB 13706 / S10) |
| J7TF58 | Clostridium sporogenes (strain ATCC 15579) |
| J7VVX4 | Bacillus cereus VD142 |
| J7W203 | Bacillus cereus VD022 |
| J8AN70 | Bacillus cereus BAG5X1-1 |
| J8B291 | Bacillus cereus BAG6X1-2 |
| J8CYF2 | Bacillus cereus HuA4-10 |
| J8EX71 | Bacillus cereus MC67 |
| J8GEI1 | Bacillus cereus MSX-D12 |
| J8HFT4 | Bacillus cereus VD014 |
| J8HJP5 | Bacillus cereus VD048 |
| J8HJY3 | Bacillus mycoides |
| J8HNV0 | Bacillus cereus VD045 |
| J8J8Y7 | Bacillus cereus VD107 |
| J8JXA8 | Bacillus cereus VD115 |
| J8KEJ8 | Bacillus cereus VD154 |
| J8L079 | Bacillus cereus VD166 |
| J8WRZ7 | Bacillus cereus BAG6O-2 |
| J9A3N1 | Bacillus wiedmannii |
| J9BNE1 | Bacillus cereus HuA2-1 |
| K0AYL8 | Gottschalkia acidurici (strain ATCC 7906 / DSM 604 / BCRC 14475 / CIP 104303 / KCTC 5404 / NCIMB 10678 / 9a) |
| K0FYJ3 | Bacillus thuringiensis MC28 |
| K0J7W6 | Amphibacillus xylanus (strain ATCC 51415 / DSM 6626 / JCM 7361 / LMG 17667 / NBRC 15112 / Ep01) |
| K1KWR9 | Solibacillus isronensis B3W22 |
| K1T604 | human gut metagenome |
| K4L163 | Dehalobacter sp. CF |
| K4L8A9 | Dehalobacter sp. CF |
| K4LJE9 | Thermacetogenium phaeum (strain ATCC BAA-254 / DSM 26808 / PB) |
| K4LJG8 | Thermacetogenium phaeum (strain ATCC BAA-254 / DSM 26808 / PB) |
| K4LS31 | Thermacetogenium phaeum (strain ATCC BAA-254 / DSM 26808 / PB) |
| K6CDL7 | Bacillus bataviensis LMG 21833 |
| K6DFM4 | Bacillus azotoformans LMG 9581 |
| K6PNB1 | Thermaerobacter subterraneus DSM 13965 |
| K8EHB8 | Desulfotomaculum hydrothermale Lam5 = DSM 18033 |
| L0F8H2 | Desulfitobacterium dichloroeliminans (strain LMG P-21439 / DCA1) |
| L0FCV5 | Desulfitobacterium dichloroeliminans (strain LMG P-21439 / DCA1) |
| L0K8D4 | Halobacteroides halobius (strain ATCC 35273 / DSM 5150 / MD-1) |
| L1QMY1 | Clostridium celatum DSM 1785 |
| L7VSI4 | Thermoclostridium stercorarium (strain ATCC 35414 / DSM 8532 / NCIMB 11754) |
| M1Q2M6 | uncultured organism |
| M1QFP2 | Bacillus thuringiensis serovar thuringiensis str. IS5056 |
| M1ZFQ2 | [Clostridium] ultunense Esp |
| M1ZWC4 | Clostridium botulinum CFSAN001627 |
| M7NH17 | Bhargavaea cecembensis DSE10 |
| M8ECB6 | Brevibacillus borstelensis AK1 |
| N1LY38 | Bacillus sp. GeD10 |
| N9WJ53 | Clostridium thermobutyricum |
| Q0AYS9 | Syntrophomonas wolfei subsp. wolfei (strain DSM 2245B / Goettingen) |
| Q0B0A3 | Syntrophomonas wolfei subsp. wolfei (strain DSM 2245B / Goettingen) |
| Q0SSR9 | Clostridium perfringens (strain SM101 / Type A) |
| Q180E2 | Clostridioides difficile (strain 630) |
| Q24MU3 | Desulfitobacterium hafniense (strain Y51) |
| Q24X43 | Desulfitobacterium hafniense (strain Y51) |
| Q2BDL5 | Bacillus sp. NRRL B-14911 |
| Q2RHM2 | Moorella thermoacetica (strain ATCC 39073 / JCM 9320) |
| Q2RIX9 | Moorella thermoacetica (strain ATCC 39073 / JCM 9320) |
| Q3EXK5 | Bacillus thuringiensis serovar israelensis ATCC 35646 |
| Q63GL9 | Bacillus cereus (strain ZK / E33L) |
| Q67P03 | Symbiobacterium thermophilum (strain T / IAM 14863) |
| Q6HP40 | Bacillus thuringiensis subsp. konkukian (strain 97-27) |
| Q73E97 | Bacillus cereus (strain ATCC 10987 / NRS 248) |
| Q81IJ0 | Bacillus cereus (strain ATCC 14579 / DSM 31 / CCUG 7414 / JCM 2152 / NBRC 15305 / NCIMB 9373 / NCTC 2599 / NRRL B-3711) |
| Q891C1 | Clostridium tetani (strain Massachusetts / E88) |
| Q8EMK7 | Oceanobacillus iheyensis (strain DSM 14371 / CIP 107618 / JCM 11309 / KCTC 3954 / HTE831) |
| Q8XK53 | Clostridium perfringens (strain 13 / Type A) |
| R1ATL3 | Caldisalinibacter kiritimatiensis |
| R1ATL9 | Caldisalinibacter kiritimatiensis |
| R4KF34 | Desulfallas gibsoniae DSM 7213 |
| R5AR75 | Firmicutes bacterium CAG:103 |
| R5D6S9 | Firmicutes bacterium CAG:555 |
| R5FGW9 | Faecalibacterium sp. CAG:1138 |
| R5H8Q9 | Firmicutes bacterium CAG:114 |
| R5IEV0 | Firmicutes bacterium CAG:124 |
| R5MXU2 | Eubacterium sp. CAG:180 |
| R5Q2B7 | Ruminococcus sp. CAG:724 |
| R5S576 | Firmicutes bacterium CAG:129 |
| R5VJ62 | Ruminococcus sp. CAG:254 |
| R5X722 | Clostridium bartlettii CAG:1329 |
| R5YSI8 | Ruminococcus sp. CAG:488 |
| R6BCN3 | Clostridium sp. CAG:169 |
| R6CKL4 | Clostridium sp. CAG:242 |
| R6DPJ1 | Firmicutes bacterium CAG:238 |
| R6DXU7 | Ruminococcus sp. CAG:563 |
| R6EN63 | Firmicutes bacterium CAG:145 |
| R6FYV5 | Clostridium sp. CAG:221 |
| R6IUK7 | Ruminococcus sp. CAG:177 |
| R6J393 | Firmicutes bacterium CAG:240 |
| R6KVE6 | Clostridium sp. CAG:265 |
| R6LRV5 | Firmicutes bacterium CAG:170 |
| R6NFJ6 | Clostridium sp. CAG:413 |
| R6P1I3 | Clostridium leptum CAG:27 |
| R6PYN2 | Faecalibacterium sp. CAG:82 |
| R6TCL0 | Ruminococcus sp. CAG:57 |
| R6U446 | Firmicutes bacterium CAG:272 |
| R6UPR3 | Oscillibacter sp. CAG:155 |
| R6VWU8 | Ruminococcus sp. CAG:382 |
| R6XR55 | Clostridium sp. CAG:349 |
| R7A017 | Ruminococcus sp. CAG:379 |
| R7BS74 | Firmicutes bacterium CAG:475 |
| R7FV06 | Eubacterium sp. CAG:841 |
| R7H5S4 | Ruminococcus sp. CAG:403 |
| R7K9N9 | Acidaminococcus sp. CAG:917 |
| R7L2L0 | Ruminococcus sp. CAG:353 |
| R7RPZ7 | Thermobrachium celere DSM 8682 |
| R7RQ10 | Thermobrachium celere DSM 8682 |
| R7ZIJ4 | Lysinibacillus sphaericus OT4b.31 |
| R8CKT0 | Bacillus cereus HuA3-9 |
| R8E529 | Bacillus cereus VD133 |
| R8GZ74 | Bacillus cereus VD196 |
| R8H0E9 | Bacillus cereus VD021 |
| R8I707 | Bacillus cereus BAG1O-1 |
| R8NGJ2 | Bacillus cereus HuB13-1 |
| R8NLF3 | Bacillus cereus (strain VD146) |
| R8PRB9 | Bacillus cereus VD136 |
| R8PTX8 | Bacillus cereus VDM053 |
| R8QZW5 | Bacillus cereus VD118 |
| R8SS86 | Bacillus cereus HuB4-4 |
| R8UIT8 | Bacillus cereus VD184 |
| R9CGG2 | Clostridium sartagoforme AAU1 |
| R9LI20 | Anaerotruncus sp. G3(2012) |
| R9LII4 | Anaerotruncus sp. G3(2012) |
| R9M946 | Oscillibacter sp. 1-3 |
| S0FWC8 | Ruminiclostridium cellobioparum subsp. termitidis CT1112 |
| S2YHJ1 | Paenisporosarcina sp. HGH0030 |
| S6EHR8 | Clostridium chauvoei JF4335 |
| T0JDL2 | Dehalobacter sp. UNSWDHB |
| T0PB17 | Clostridium sp. BL8 |
| T2RE34 | Paeniclostridium sordellii (strain ATCC 9714 / DSM 2141 / JCM 3814 / LMG 15708 / NCIMB 10717 / 211) |
| T2RMA2 | Dehalobacter sp. UNSWDHB |
| T3D8U9 | Clostridioides difficile CD160 |
| T4VKR6 | Paraclostridium bifermentans ATCC 638 |
| T4VVP0 | Paraclostridium bifermentans ATCC 19299 |
| U2CV98 | Clostridiales bacterium oral taxon 876 str. F0540 |
| U2DE89 | Clostridium sp. ATCC BAA-442 |
| U2EBP6 | Haloplasma contractile SSD-17B |
| U2N011 | Clostridium intestinale URNW |
| U2SIV8 | Oscillibacter sp. KLE 1745 |
| U4QZW2 | Ruminiclostridium papyrosolvens C7 |
| U5LGV4 | Bacillus infantis NRRL B-14911 |
| U6SKK9 | Bacillus marmarensis DSM 21297 |
| V2Y0S3 | Firmicutes bacterium ASF500 |
| V2YJ22 | Firmicutes bacterium ASF500 |
| V5M3G1 | Bacillus thuringiensis YBT-1518 |
| V6M837 | Brevibacillus panacihumi W25 |
| V6TC90 | Bacillus sp. 17376 |
| V9H1S1 | Clostridium sp. 7_2_43FAA |
| W0EBV3 | Desulfitobacterium metallireducens DSM 15288 |
| W0ECK7 | Desulfitobacterium metallireducens DSM 15288 |
| W0U3U4 | Ruminococcus bicirculans |
| W1SEP7 | Bacillus vireti LMG 21834 |
| W4C6U1 | Paenibacillus sp. FSL R7-269 |
| W4DW27 | Paenibacillus sp. FSL R7-277 |
| W4EZQ7 | Viridibacillus arenosi FSL R5-213 |
| W4Q9Q5 | Bacillus wakoensis JCM 9140 |
| W4QQM3 | Bacillus akibai (strain ATCC 43226 / DSM 21942 / JCM 9157 / 1139) |
| W4RL53 | Bacillus boroniphilus JCM 21738 |
| W4V9Q1 | Hungateiclostridium straminisolvens JCM 21531 |
| W7KY04 | Bacillus firmus DS1 |
| W7SEJ5 | Lysinibacillus sphaericus CBAM5 |
| W7YG53 | Paenibacillus pini JCM 16418 |
| W8Y4X0 | Bacillus thuringiensis DB27 |
| W9BER1 | Oceanobacillus picturae |
