## Supplement Data Table2 for "Dual-wield NTPases: a novel protein family mined from AlphaFold DB"

Supplement Data Table 2: List of ESM metagenomic Atlas entries structurally related to dwNTPase. Note that the target database was culled by sequence similarity of 30% identity and pLDDT threshold.

MGYP000005464415

MGYP000011566305

MGYP000011743424

MGYP000013930452

MGYP000021985546

MGYP000022775823

MGYP000031243067

MGYP000032816584

MGYP000035039818

MGYP000039754873

MGYP000046075001

MGYP000046921890

MGYP000050773439

MGYP000056714802

MGYP000058475240

MGYP000063702171

MGYP000072106316

MGYP000073817642

MGYP000073976865

MGYP000074926707

MGYP000087961665

MGYP000090814326

MGYP000095660081

MGYP000097934424

MGYP000100781166

MGYP000110243900

MGYP000121404939

MGYP000132674256

MGYP000133036970

MGYP000137384458

MGYP000137884487

MGYP000138095260

MGYP000144053071

MGYP000149276925

MGYP000152158533

MGYP000155467865

MGYP000158031104

MGYP000158701616

MGYP000161370762

MGYP000172069787

MGYP000179168143

MGYP000182400474

MGYP000187260450

MGYP000196473938

MGYP000203777768

MGYP000206671460

MGYP000209962063

MGYP000213213017

MGYP000214913979

MGYP000223653340

MGYP000226615218

MGYP000229661469

MGYP000233862775

MGYP000260203210

MGYP000267655121

MGYP000276180941

MGYP000281675394

MGYP000283494147

MGYP000289560546

MGYP000297351230

MGYP000307241419

MGYP000316400542

MGYP000317455607

MGYP000329090296

MGYP000329155775

MGYP000332792456

MGYP000338656182

MGYP000338766891

MGYP000338885734

MGYP000339528302

MGYP000341965729

MGYP000344960023

MGYP000349730980

MGYP000356359273

MGYP000357322122

MGYP000362356478

MGYP000365138801

MGYP000366121951

MGYP000369246366

MGYP000369248057

MGYP000374059993

MGYP000376728391

MGYP000377349853

MGYP000380150569

MGYP000380918257

MGYP000387215517

MGYP000389137078

MGYP000392093694

MGYP000392322862

MGYP000397624302

MGYP000412013239

MGYP000414862069

MGYP000415449340

MGYP000415610050

MGYP000418125265

MGYP000424919781

MGYP000425215761

MGYP000427578989

MGYP000432980634

MGYP000434790480

MGYP000440675606

MGYP000449267722

MGYP000453218129

MGYP000468120591

MGYP000479063216

MGYP000481223319

MGYP000495801975

MGYP000501602264

MGYP000516580646

MGYP000518541956

MGYP000522568599

MGYP000540603521

MGYP000544980939

MGYP000554953503

MGYP000560810235

MGYP000563868109

MGYP000565819564

MGYP000570892856

MGYP000573565277

MGYP000580686002

MGYP000583596164

MGYP000598313075

MGYP000609899509

MGYP000610054608

MGYP000619824976

MGYP000621160421

MGYP000624467883

MGYP000624854459

MGYP000626659218

MGYP000628622447

MGYP000633643098

MGYP000635340560

MGYP000637753403

MGYP000640418712

MGYP000651216849

MGYP000656668235

MGYP000665329988

MGYP000668644108

MGYP000671723777

MGYP000672861390

MGYP000674781713

MGYP000680700556

MGYP000680727538

MGYP000687400265

MGYP000688418485

MGYP000692424004

MGYP000694260436

MGYP000712421253

MGYP000712598401

MGYP000715366489

MGYP000716100718

MGYP000716110458

MGYP000719069569

MGYP000719259666

MGYP000720987161

MGYP000722397558

MGYP000737789640

MGYP000745950594

MGYP000760084081

MGYP000766970156

MGYP000773444244

MGYP000776980989

MGYP000779413157

MGYP000783606972

MGYP000794234970

MGYP000794583975

MGYP000796842517

MGYP000800695632

MGYP000818155992

MGYP000819680346

MGYP000829006276

MGYP000833698882

MGYP000837620560

MGYP000845675048

MGYP000846291321

MGYP000846738100

MGYP000847433322

MGYP000850522706

MGYP000854750270

MGYP000855463610

MGYP000856935708

MGYP000859444552

MGYP000860510931

MGYP000862261671

MGYP000862978595

MGYP000863910226

MGYP000864615565

MGYP000865798123

MGYP000866159967

MGYP000870042540

MGYP000872154735

MGYP000873111992

MGYP000873114169

MGYP000873298085

MGYP000873570358

MGYP000874498255

MGYP000875228748

MGYP000876619059

MGYP000881585069

MGYP000882995862

MGYP000883449302

MGYP000884151462

MGYP000884157304

MGYP000886298914

MGYP000886527711

MGYP000887942385

MGYP000889347843

MGYP000891638238

MGYP000893577897

MGYP000893585008

MGYP000896138925

MGYP000896152022

MGYP000902764005

MGYP000905320016

MGYP000905590446

MGYP000906464546

MGYP000911693974

MGYP000912395745

MGYP000913366715

MGYP000917325413

MGYP000918987734

MGYP000920861653

MGYP000927488903

MGYP000929340741

MGYP000929694189

MGYP000932862644

MGYP000934286412

MGYP000937807594

MGYP000938080560

MGYP000939256813

MGYP000943814674

MGYP000944892989

MGYP000946285437

MGYP000948302140

MGYP000948387078

MGYP000948665050

MGYP000948773478

MGYP000949039413

MGYP000950522601

MGYP000950781180

MGYP000952638058

MGYP000953940834

MGYP000954761325

MGYP000956436473

MGYP000965532308

MGYP000966778367

MGYP000967381621

MGYP000967452070

MGYP000969341437

MGYP000971973798

MGYP000981829549

MGYP000982030885

MGYP000982564017

MGYP000983360511

MGYP000983698559

MGYP000983968897

MGYP000984007444

MGYP000984069175

MGYP000986005084

MGYP000986772698

MGYP000987235152

MGYP000987950999

MGYP000989346284

MGYP000991037961

MGYP000993126788

MGYP000996428994

MGYP000998089494

MGYP000999504741

MGYP001000915644

MGYP001001350963

MGYP001002397009

MGYP001003736678

MGYP001004190013

MGYP001004907671

MGYP001008415023

MGYP001009845231

MGYP001010016555

MGYP001010081099

MGYP001010801421

MGYP001012929272

MGYP001015136735

MGYP001018576274

MGYP001019263009

MGYP001019466318

MGYP001022181482

MGYP001022569621

MGYP001023499417

MGYP001024206109

MGYP001031223130

MGYP001032263427

MGYP001033019173

MGYP001036146819

MGYP001037691505

MGYP001041136564

MGYP001044408008

MGYP001044653672

MGYP001046357902

MGYP001051366889

MGYP001051616043

MGYP001052389888

MGYP001053934307

MGYP001054709820

MGYP001055124250

MGYP001057389601

MGYP001057713172

MGYP001057939588

MGYP001058076732

MGYP001066949444

MGYP001067596079

MGYP001067597097

MGYP001070558268

MGYP001071062081

MGYP001072290625

MGYP001072290727

MGYP001072684865

MGYP001074592748

MGYP001074594893

MGYP001075631168

MGYP001076246188

MGYP001076286658

MGYP001077760211

MGYP001078469938

MGYP001081539167

MGYP001083094903

MGYP001086549199

MGYP001087267426

MGYP001089574896

MGYP001095254367

MGYP001097077469

MGYP001099651039

MGYP001101482462

MGYP001102279279

MGYP001104087068

MGYP001105578980

MGYP001111995788

MGYP001118306118

MGYP001119547012

MGYP001120777949

MGYP001122123002

MGYP001124793146

MGYP001130624379

MGYP001132418681

MGYP001136708707

MGYP001139646309

MGYP001140236529

MGYP001140486808

MGYP001141881270

MGYP001144037523

MGYP001147184671

MGYP001147701674

MGYP001148728985

MGYP001151314202

MGYP001153095883

MGYP001153238483

MGYP001153393980

MGYP001153741911

MGYP001158822045

MGYP001163045142

MGYP001167325490

MGYP001170022268

MGYP001173886188

MGYP001178305136

MGYP001190237233

MGYP001193054289

MGYP001198073675

MGYP001199854867

MGYP001203888545

MGYP001205242401

MGYP001208954122

MGYP001212037890

MGYP001216256983

MGYP001226913156

MGYP001229757033

MGYP001230977503

MGYP001236415831

MGYP001238894987

MGYP001240458690

MGYP001241439385

MGYP001241830453

MGYP001244424814

MGYP001245476879

MGYP001246862665

MGYP001251410889

MGYP001256856623

MGYP001258079375

MGYP001260779301

MGYP001266339757

MGYP001270284586

MGYP001274359445

MGYP001282449832

MGYP001293271363

MGYP001298300523

MGYP001298735028

MGYP001306662269

MGYP001308373670

MGYP001310676587

MGYP001316291143

MGYP001317270614

MGYP001317651948

MGYP001321326270

MGYP001324221372

MGYP001325391279

MGYP001328123680

MGYP001335462562

MGYP001343381432

MGYP001343531413

MGYP001347097104

MGYP001349776869

MGYP001350155789

MGYP001370477403

MGYP001375895633

MGYP001375895852

MGYP001377036890

MGYP001377380736

MGYP001386727829

MGYP001389434110

MGYP001390933540

MGYP001394707573

MGYP001397346626

MGYP001400061178

MGYP001407994366

MGYP001408430282

MGYP001410709774

MGYP001411239173

MGYP001416624695

MGYP001430084181

MGYP001431026079

MGYP001431417008

MGYP001432564324

MGYP001437982201

MGYP001446108616

MGYP001464086068

MGYP001465057784

MGYP001470679893

MGYP001472048744

MGYP001480157240

MGYP001481512249

MGYP001482463118

MGYP001484212884

MGYP001484214231

MGYP001486922552

MGYP001487875866

MGYP001516528669

MGYP001522023819

MGYP001535160546

MGYP001537081954

MGYP001538263980

MGYP001598895366

MGYP001601149468

MGYP001620285363

MGYP001624020629

MGYP001625272499

MGYP001630033458

MGYP001630426298

MGYP001630889881

MGYP001632728344

MGYP001638034195

MGYP001639344602

MGYP001657894984

MGYP001770710926

MGYP001771164780

MGYP001771283061

MGYP001774661584

MGYP001775303000

MGYP001776077675

MGYP001777284512

MGYP001777798491

MGYP001777999770

MGYP001778362933

MGYP001780621392

MGYP001781026291

MGYP001783202305

MGYP001783919854

MGYP001829439330

MGYP001851345070

MGYP001852003032

MGYP001869686281

MGYP001944123563

MGYP001944327984

MGYP001952770736

MGYP001953138799

MGYP001953984248

MGYP001954818258

MGYP002268305163

MGYP002348050470

MGYP002388813658

MGYP002398377235

MGYP002407614603

MGYP002409113681

MGYP002410704882

MGYP002505411068

MGYP002508460163

MGYP002508579337

MGYP002509702026

MGYP002509779980

MGYP002510528239

MGYP002512426959

MGYP002512774708

MGYP002513981819

MGYP002514523331

MGYP002514735286

MGYP002515637646

MGYP002516001785

MGYP002516138547

MGYP002516672976

MGYP002516965467

MGYP002517707201

MGYP002518695557

MGYP002518877164

MGYP002518946026

MGYP002519335693

MGYP002519536267

MGYP002519891592

MGYP002520059971

MGYP002520388537

MGYP002520843664

MGYP002521152857

MGYP002521245368

MGYP002521741868

MGYP002521823067

MGYP002522324169

MGYP002523679548

MGYP002526212556

MGYP002531393319

MGYP002535785097

MGYP002536951133

MGYP002538392298

MGYP002540006505

MGYP002542236210

MGYP002544833878

MGYP002545730500

MGYP002546544705

MGYP002547882387

MGYP002548274122

MGYP002548845381

MGYP002550710106

MGYP002551048834

MGYP002552576849

MGYP002553774068

MGYP002554244177

MGYP002557624238

MGYP002563089458

MGYP002563477401

MGYP002564862066

MGYP002565572074

MGYP002565576012

MGYP002569519302

MGYP002569928787

MGYP002570950553

MGYP002572209789

MGYP002580689582

MGYP002584680938

MGYP002584950757

MGYP002604693242

MGYP002625390995

MGYP002626635166

MGYP002647315051

MGYP002647372959

MGYP002648415025

MGYP002655976423

MGYP002672574807

MGYP002672701855

MGYP002675175697

MGYP002679840418

MGYP002705934524

MGYP002710575024

MGYP002728503105

MGYP002731537162

MGYP002732731958

MGYP002733154163

MGYP002733396199

MGYP002735076069

MGYP002748286501

MGYP002751119144

MGYP002753153100

MGYP002797240956

MGYP002798388229

MGYP002802551986

MGYP002857010239

MGYP002867796410

MGYP002903955578

MGYP002960176274

MGYP002972649115

MGYP003013740277

MGYP003101485480

MGYP003104526251

MGYP003162439181

MGYP003181990754

MGYP003184855784

MGYP003185590153

MGYP003195452491

MGYP003209727232

MGYP003210669313

MGYP003220877510

MGYP003266687897

MGYP003280304731

MGYP003283676434

MGYP003288713578

MGYP003288879220

MGYP003289010650

MGYP003289121590

MGYP003289460875

MGYP003289554861

MGYP003289654488

MGYP003289736391

MGYP003290012312

MGYP003290061622

MGYP003290476675

MGYP003290532250

MGYP003290763530

MGYP003290847745

MGYP003290873295

MGYP003290982398

MGYP003291326079

MGYP003291517417

MGYP003291711009

MGYP003291766561

MGYP003292095451

MGYP003292202049

MGYP003292347188

MGYP003293279942

MGYP003293463256

MGYP003293661037

MGYP003294162239

MGYP003294245754

MGYP003294696003

MGYP003295022881

MGYP003295543339

MGYP003295677572

MGYP003295683491

MGYP003296276390

MGYP003296445675

MGYP003297612339

MGYP003297636011

MGYP003298888900

MGYP003299105919

MGYP003299134100

MGYP003299421817

MGYP003299624148

MGYP003299798750

MGYP003300215213

MGYP003301305664

MGYP003301463800

MGYP003302090907

MGYP003302257712

MGYP003302551956

MGYP003302844486

MGYP003303284055

MGYP003303453267

MGYP003303602865

MGYP003303613744

MGYP003303857016

MGYP003303883274

MGYP003304453194

MGYP003304735657

MGYP003305165249

MGYP003305203406

MGYP003305708281

MGYP003306062031

MGYP003307021237

MGYP003307705220

MGYP003307919367

MGYP003308042158

MGYP003308684829

MGYP003308840078

MGYP003308867545

MGYP003308877326

MGYP003309211794

MGYP003309457844

MGYP003309930354

MGYP003310390468

MGYP003310719773

MGYP003311209964

MGYP003311456034

MGYP003311669213

MGYP003315797261

MGYP003315959051

MGYP003316642960

MGYP003316655320

MGYP003317707224

MGYP003318230597

MGYP003318922623

MGYP003319073510

MGYP003319556249

MGYP003319980098

MGYP003324755819

MGYP003329257096

MGYP003339773965

MGYP003345614305

MGYP003362406859

MGYP003365329244

MGYP003369291436

MGYP003369572669

MGYP003369774084

MGYP003369841141

MGYP003371456443

MGYP003373848938

MGYP003375948820

MGYP003376599254

MGYP003377696461

MGYP003378316808

MGYP003379681987

MGYP003385241863

MGYP003385333612

MGYP003393809899

MGYP003400006688

MGYP003421748675

MGYP003427310168

MGYP003430666403

MGYP003430775767

MGYP003435514729

MGYP003447008246

MGYP003466133340

MGYP003467745854

MGYP003474320691

MGYP003482910043

MGYP003484242520

MGYP003508593118

MGYP003509481182

MGYP003510406608

MGYP003515515405

MGYP003518594222

MGYP003520657654

MGYP003522064156

MGYP003526856239

MGYP003528103500

MGYP003530985645

MGYP003531931634

MGYP003533196885

MGYP003533852953

MGYP003542505439

MGYP003547168537

MGYP003550500512

MGYP003552847930

MGYP003558080982

MGYP003558878903

MGYP003564231365

MGYP003571335074

MGYP003585492775

MGYP003592361735

MGYP003623391267

MGYP003623576751

MGYP003623868684

MGYP003764291581

MGYP003783283803

MGYP003819602827

MGYP003900866323

MGYP003901033641
